# AlphaGenome 2 predicts regulatory activity and variant effects across cell states

**DOI:** 10.64898/2026.11.11.765857

**Authors:** Adam Gayoso, Anne Mottram, Alexander Karollus, Guido Novati, Pavol Drotár, John Reid, Laetitia Meng-Papaxanthos, Adam Kosiorek, Eirini Arvaniti, Vincent Dutordoir, Soham De, Tom Ward, Lai Hong Wong, Matteo Perino, Kyle Taylor, Jun Cheng, Natasha Latysheva, Joshua Pan, Dhavanthi Hariharan, Clare Bycroft, Lauren Nicolaisen, Demis Hassabis, Pushmeet Kohli, Žiga Avsec

**Affiliations:** Google DeepMind

## Abstract

Most genetic variants associated with human traits and diseases lie in non-coding DNA, where they can alter gene regulation in specific cell types and states. Single-cell atlases chart these states across tissues and development, but rarely profile enough donors to associate variants with expression directly. Deep learning models could bridge this gap by predicting variant effects from DNA sequence, yet they poorly distinguish related cell states and require fine-tuning for new datasets. Here we present AlphaGenome 2 (AG2), which substantially advances cell-type-specific prediction of regulatory activity and variant effects for gene expression and chromatin accessibility. Trained on organism-wide human and mouse single-cell atlases and bulk functional genomics data, AG2 takes as input a 1-Mb DNA sequence and a condition specifying a target tissue, cell type, or cell state. Conditioning enables AG2 to impute unmeasured epigenomic tracks and predict multimodal variant effects across cell types. Inferring this condition directly from gene expression extends AG2 to unseen datasets and cell states without fine-tuning. AG2 outperforms fine-tuned AlphaGenome 1 and specialized models trained on target datasets, doubling recall of causal single-cell eQTLs at 90% precision. AG2 also achieves state-of-the-art enhancer–gene linking and can pinpoint causal variants at complex trait loci. Across hundreds of human cell types, we generated an atlas of enhancer–gene links and scores for 7 million common variants. As single-cell profiling expands across development, tissues, and disease cohorts, AG2 provides a blueprint for turning observational atlases into variant effect maps.

## Introduction

A single human genome gives rise to hundreds of distinct cell types and states, whose different gene expression programs are driven by selective utilization of non-coding regulatory sequences. Single-cell initiatives such as the Human Cell Atlas^1^ and repositories such as CELLxGENE^2^ have rapidly cataloged transcriptional and chromatin states across primary tissues, development, and disease^3–6^, but our understanding of how non-coding regulatory variation affects these cell states has lagged behind. Observational single-cell atlases typically sample too few donors to map genetic effects directly. Resolving variant effects through single-cell quantitative trait locus (QTL) mapping requires larger cohorts, as recently exemplified by TenK10K^7^. Even at this scale, QTL studies do not clearly separate causal variants from those in linkage disequilibrium (LD), lack statistical power for rare variants, and remain largely restricted to few accessible tissues. These factors contribute to a systematic gap between molecular QTLs and complex-trait loci from genome-wide association studies (GWAS)^8^. Experimental perturbations face complementary constraints: massively parallel reporter assays (MPRAs) test sequences outside native chromatin context^9,10^, whereas endogenous editing approaches scale to only a limited set of loci and tractable cell types^11–13^.

Deep learning sequence-to-function (S2F) models could complement these experimental constraints by predicting regulatory activity and variant effects in different cell types directly from DNA sequence. We recently described AlphaGenome^14^ (hereafter AlphaGenome 1, or AG1), a state-of-the-art unified S2F model that predicts diverse molecular readouts at up to single-base resolution from a 1-Mb human or mouse DNA sequence. Yet like earlier multitask S2F models^15,16^, AlphaGenome 1 relies predominantly on bulk tissues and cell lines, predicts a fixed catalog of experimentally measured tracks, and struggles to distinguish closely related cell types. Recent studies have trained S2F models directly on single-cell data^17–21^ or conditioned models on tissues, cell types, or cell states^21–24^ to impute unmeasured modalities^22,24^ and generalize to unseen samples^23,24^. However, neither single-cell training nor conditioning alone has been shown to substantially improve cell-type specificity or variant effect prediction over multitask models such as Borzoi^16^ and AlphaGenome 1. Weak cell-type specificity across training in turn constrains generalization to unseen cell types and states. Therefore, advances in both predictive performance and data coverage are required to enable cell-type-resolved variant interpretation for single-cell atlases.

Here we present AlphaGenome 2 (AG2), a conditional S2F model that advances both predictive performance and coverage across cell types and states, predicting regulatory activity and variant effects from a 1-Mb human or mouse DNA sequence. To improve performance across closely related cell types, AlphaGenome 2 represents a specified tissue, cell line, cell type, or cell state (which we refer to as a condition) as an embedding that modulates sequence representations throughout the network, starting from the earliest layers. We trained AlphaGenome 2 across bulk functional genomics data, cell-type-resolved DNA methylation, and organism-wide human and mouse single-cell RNA-seq (scRNA-seq) and ATAC-seq (scATAC-seq) atlases. Together, early conditioning and expanded training data substantially sharpen cell-type specificity relative to unconditioned multitask and late-conditioned models, enabling AlphaGenome 2 to outperform fine-tuned AlphaGenome 1 and specialized single-cell models^18,21^ on expression QTL (eQTL) variant effect prediction and distal enhancer–gene (E2G) linking. To expand coverage across modalities and unseen cell states, AlphaGenome 2 decouples conditions from output heads to impute unmeasured epigenomic assays in primary cells, and introduces a post-training stage that aligns a single-cell expression autoencoder trained on over 100 million human scRNA-seq profiles into the frozen sequence model’s condition space. Inferring condition embeddings directly from scRNA-seq extends AlphaGenome 2 to newly profiled cohorts and rare cell states without fine-tuning, outperforming models trained directly on those cohorts^18,21^. Leveraging AlphaGenome 2, we predict lineage-specific distal E2G links genome-wide and score nearly 7 million trait-associated and common variants to help annotate complex traits with putatively causal variants, target genes with direction of effect, and affected cell states.

## Results

### Condition-specific sequence modeling

AlphaGenome 2 is a deep learning model that takes as input a 1-Mb DNA sequence, a target condition (tissue, cell line, cell type, or cell state), and a species identifier (human or mouse) and predicts genomic tracks spanning diverse molecular readouts (Fig. 1a–c). Built on a U-Net-style backbone^25^ similar to AG1^14^, the model integrates proximal and distal sequence context across the 1-Mb window (Fig. 1b). To capture condition-specific regulation, adaptive normalization layers^26^ modulate intermediate sequence representations using a condition embedding, which is either selected from a learned dictionary of ontology-defined reference conditions or generated from scRNA-seq profiles via a cell encoder (Fig. 1a; Methods). For a given condition, these predictions take the form of base- and bin-resolution tracks (spanning RNA coverage, transcription initiation (CAGE), RNA splicing, chromatin accessibility, histone modifications, transcription factor (TF) binding, and CpG methylation), scalar gene expression values, and pairwise interactions representing chromatin contact maps and splice junctions (Fig. 1c).

**Figure 1.**
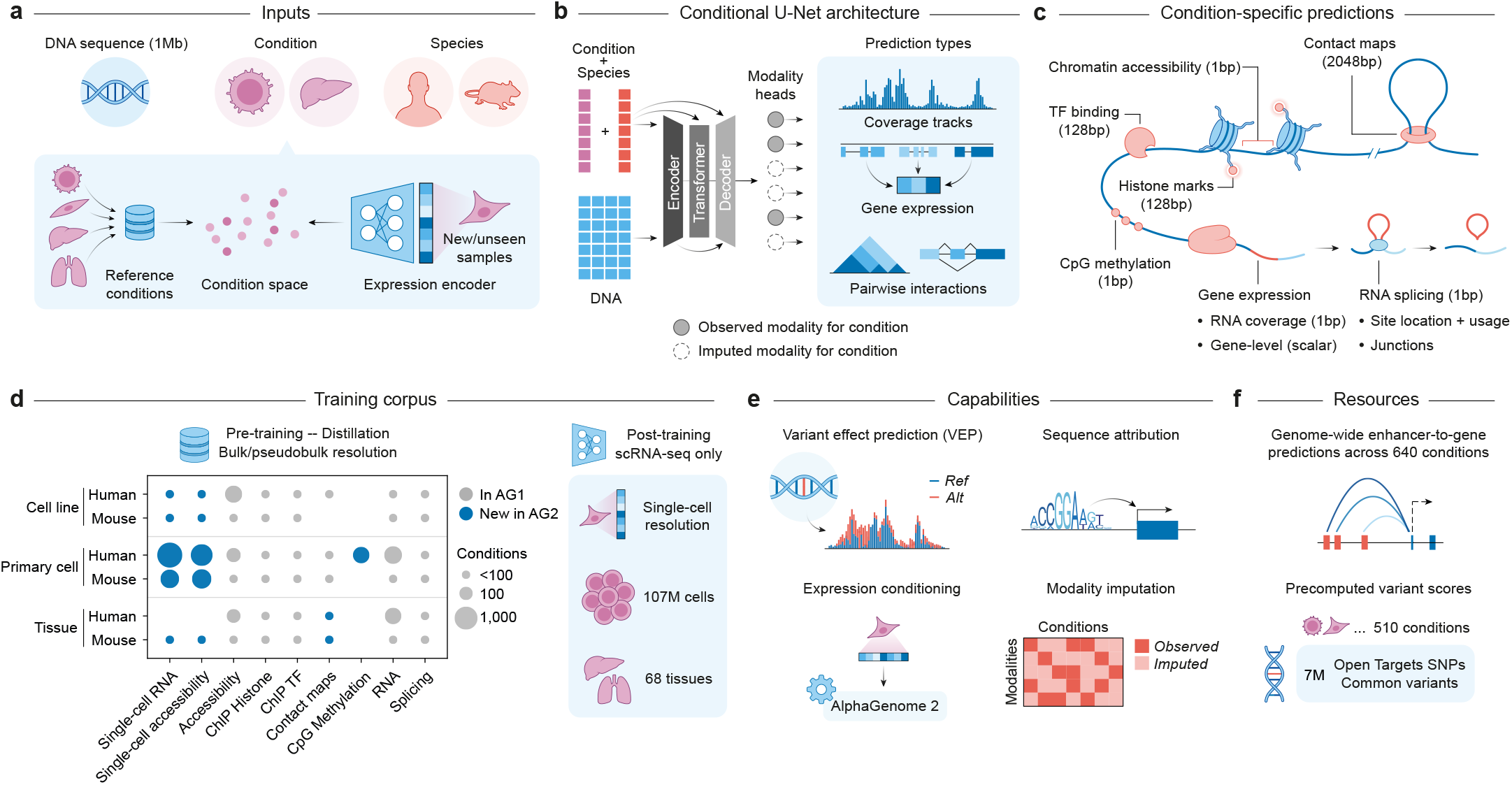
AlphaGenome 2 adapts sequence representations to each condition to predict multimodal genomic readouts in observed and unseen cell states. **(a)** AG2 takes as input a 1-Mb DNA sequence, a target condition, and species identifier (human or mouse). Biological conditioning is specified either from a predetermined set of discrete reference conditions or from single-cell gene expression data for generalization to unseen primary cell types and samples (Methods). **(b)** Biological and species conditioning occurs throughout the architecture of AG2, modulating sequence representations across the U-Net encoder, central transformer tower, and decoder. For each condition, task-specific modality heads predict both experimentally observed modalities and imputed modalities, which span coverage tracks, scalar gene-level expression, and pairwise chromatin interactions. **(c)** For each condition, AG2 predicts transcription factor (TF) binding (128-bp resolution), histone modifications (128 bp), chromatin accessibility (ATAC-seq and DNase-seq, 1 bp), CpG DNA methylation (1 bp), chromatin contact maps (2048 bp), gene expression (1-bp stranded CAGE and RNA-seq coverage and scalar RNA-seq gene-level counts), and RNA splicing (1-bp splice site location, usage, and splice junctions). **(d)** AG2 extends the AG1 bulk training corpus with single-cell RNA-seq, single-cell ATAC-seq, and CpG methylation datasets. Additionally, its single-cell expression encoder was trained on over 100 million individual scRNA-seq profiles across 68 tissues. **(e)** Capabilities of the model include cell-type-resolved variant effect prediction (VEP; comparing reference and alternative allele profiles), nucleotide-resolution sequence attribution via *in silico* mutagenesis (ISM), zero-shot generalization to unseen conditions, and cross-modality epigenomic imputation. **(f)** Precomputed predictions accompanying release include an enhancer–gene resource (genome-wide predictions across 640 conditions) and a GWAS-centric variant effect resource (covering 7 million variants including Open Targets Platform^27^ single-nucleotide polymorphisms (SNPs) and common variants, across 510 conditions).

AG2 was trained in four stages (pre-training, condition refinement, distillation, and post-training) on the reference human and mouse genomes and an expansive corpus of bulk and single-cell genomics data (Fig. 1d; Supplementary Fig. 1; Methods). Compared to AG1, which was trained exclusively on bulk assays, AG2 introduces 3977 single-cell RNA-seq and ATAC-seq pseudobulk tracks across 36 human and mouse studies (many of which featured profiling of both modalities; Supplementary Table 1), adds cell-type-resolved CpG methylation spanning 185 human reference conditions^28,29^, and post-trains on single cells sampled from 107 million human scRNA-seq profiles (Supplementary Tables 2 and 3). Across this corpus, bulk assays and cell-type-aggregated single-cell pseudobulks are indexed by standard ontology terms into a fixed catalog of reference conditions, each representing a distinct human or mouse tissue, cell line, or cell type and grouping all modalities measured for that condition (Methods). Eight randomly initialized AG2 models were first pre-trained across these reference conditions and then refined by further optimizing their reference condition embeddings and modulation layers with the rest of the model frozen, yielding eight “teacher” models. For distillation, a single AG2 “student” model sharing the same architecture was trained to reproduce the ensemble-averaged predictions of the teachers on randomly mutated input sequences, transferring the robustness and variant effect prediction accuracy of the teacher ensemble into a single model^14,30^. Finally, the student model was post-trained at the single-cell level on human scRNA-seq profiles from CELLxGENE^2^ and our human single-cell pre-training corpus, freezing the distilled model and mapping embeddings from a separately trained single-cell expression autoencoder into AG2’s condition space.

Because conditions are decoupled from shared modality heads and the distilled student backbone remains frozen during post-training, a single AG2 model operates across both reference and expression conditioning. This design supports cross-modality assay imputation in sparsely profiled conditions, adaptation to unseen scRNA-seq cohorts without retraining, and variant effect scoring and sequence attribution across modalities (Fig. 1e). Across our benchmarks, we evaluate dedicated models trained with genomic intervals, experimental tracks, or single-cell datasets held out (Methods), while applying the full-genome AG2 model across hundreds of human conditions to construct precomputed atlases of distal E2G links and GWAS variant effect predictions (Fig. 1f).

### Improved condition-specific predictions

Accurately resolving differences in gene expression and chromatin accessibility across tissues and cell types remains a major challenge for S2F models^31–33^, and at the single-cell level this difficulty is compounded by the technical noise of single-cell assays. To evaluate AG2’s capacity to address these limitations, we assessed predictions across the set of reference biological conditions using a teacher model trained with test genome intervals held out. We constructed a directly comparable baseline (AG1-sc) by fine-tuning AG1 on the single-cell reference conditions in our training corpus using the same genome split (Methods). Because AG1 outperforms Borzoi at predicting condition-specific gene expression^14^, AG1-sc also provides a strong baseline relative to Borzoi-based single-cell models.

We first evaluated the prediction of gene expression log fold changes (LFCs), computed per gene relative to the mean across bulk tissues or across pseudobulked cell types within individual tissues. This approach isolates condition-specific variation from overall expression level (Methods). Across GTEx tissues^34^, AG2 outperformed AG1, yielding a median per-tissue Pearson correlation of *r* = 0.71 (vs. 0.61) and explaining 53% of cross-tissue expression variance relative to the tissue mean (41%) (Fig. 2a). We observed comparable improvements on pseudobulk scRNA-seq data across human and mouse tissues, where AG2 explained a median of 34% of within-tissue cell-type-specific expression variance across comparison groups, compared with 23% for AG1-sc (Fig. 2b, Supplementary Fig. 3a). Extending this analysis to LFCs of single-cell chromatin accessibility signal across regulatory peaks, AG2 similarly captured a larger fraction of cell-type-specific variance (median 60% vs. 51%; Fig. 2b, Supplementary Fig. 3a).

**Figure 2.**
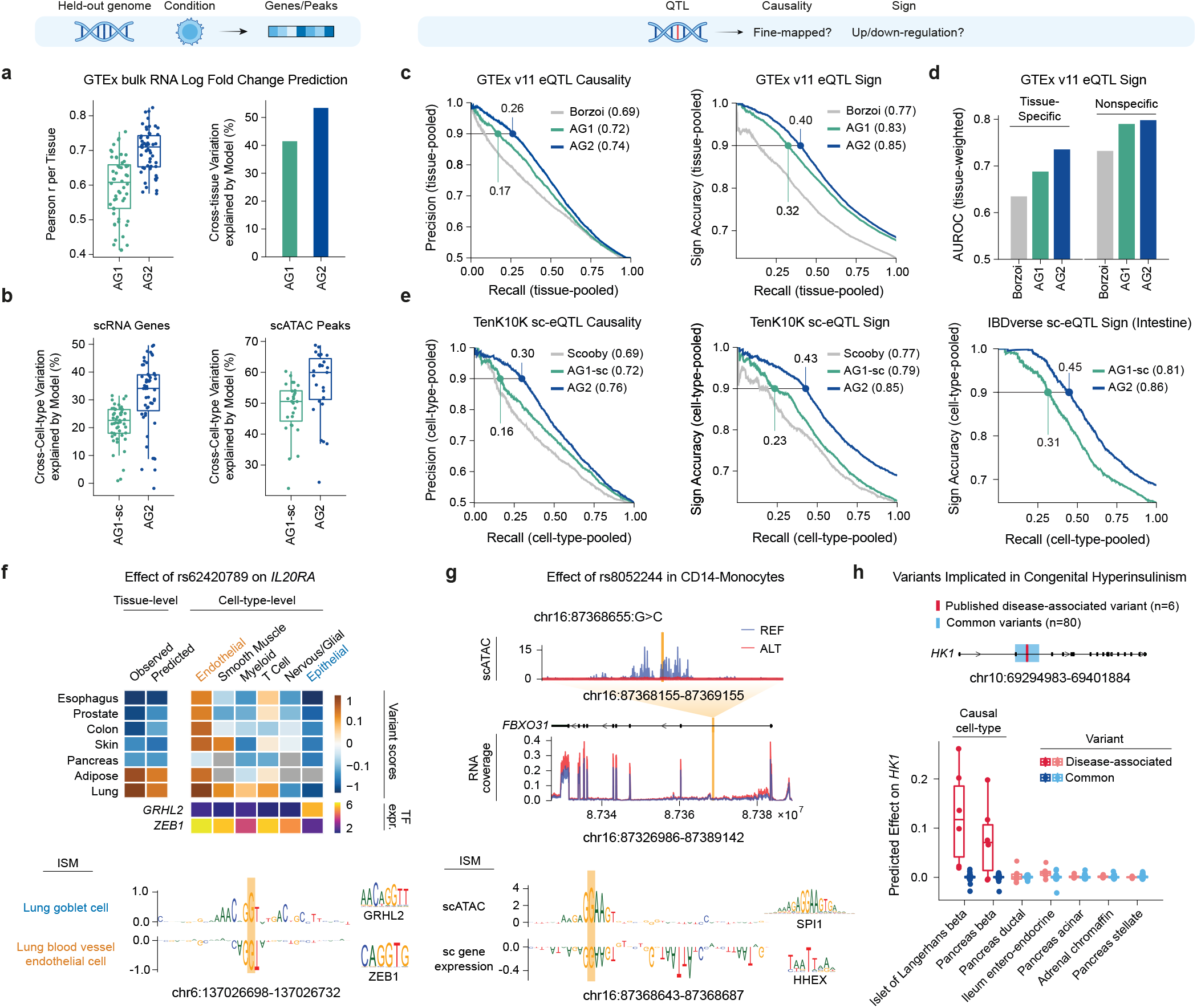
AlphaGenome 2 improves condition-specific gene expression and regulatory variant effect prediction across tissues and primary cell types. **(a)** Bulk tissue gene expression prediction on held-out genomic intervals across GTEx tissues (*n* = 52 tissues; 1885 genes) for AlphaGenome 1 (AG1) and AlphaGenome 2 (AG2). Left: Per-tissue Pearson correlation (*r*) between observed and predicted cross-tissue gene log fold changes (LFCs). Right: Percentage of cross-tissue gene expression variation explained. Box plots (left) display the median (center line), the 25th and 75th percentiles (box bounds), and the whiskers extend to 1.5 times the interquartile range from the box bounds; all data points are overlaid. **(b)** Percentage of within-tissue cross-cell-type variation explained on held-out genomic intervals by AG1 fine-tuned on single-cell reference conditions (AG1-sc) and AG2 for single-cell gene expression (left) and single-cell chromatin accessibility peaks (right). Each point represents a tissue/dataset comparison group (Methods). Left (right): *n* = 51 (27) groups, comprising 11 (10) mouse and 40 (17) human. Box plots are defined as in **a. (c)** GTEx v11 bulk tissue eQTL evaluation across 49 tissues comparing Borzoi, AG1, and AG2. Left: Tissue-pooled precision–recall curves for distinguishing fine-mapped eQTLs (*n* = 33,837 unique variant–tissue–gene combinations) from distance-matched negative controls at a 1:1 ratio (causality; AUPRC in legend). Right: Cumulative sign accuracy versus recall (sign; area under curve in legend, 51.3% positive, *n* = 33,829). Point callouts indicate recall at 90% precision (left) or 90% sign accuracy (right). **(d)** Tissue-weighted mean AUROC for GTEx v11 eQTL sign prediction stratified into tissue-specific versus nonspecific eQTLs (Methods; *n* = 7169 specific and *n* = 26,660 nonspecific eQTLs). **(e)** Single-cell QTL evaluation across primary peripheral blood immune (TenK10K) and intestinal (IBDverse) cell types. Left and middle: Cell-type-pooled precision–recall (causality, 1:1 negative-to-positive ratio) and sign accuracy–recall (sign, 47.8% positive) curves for 6346 causal sc-eQTLs across 28 cell types, comparing Scooby, AG1-sc, and AG2. Right: Sign accuracy–recall (sign, 48.5% positive) curves for 3524 causal sc-eQTLs across 47 cell types. Point callouts indicate recall at 90% precision or 90% sign accuracy. **(f)** Opposing cell-type-specific effects of rs62420789 on *IL20RA*. Top: Observed and AG2-predicted tissue-level eQTL scores across seven GTEx tissues (left), AG2 cell-type-level variant scores across constituent lineages (right), and expression of *GRHL2* and *ZEB1* (bottom rows). Bottom: *In silico* mutagenesis (ISM) attribution logos (variant shaded in orange) with matched reference motifs (right) in lung goblet cells (activating GRHL2 motif, JASPAR^36^ MA1105.3) and lung blood vessel endothelial cells (repressive ZEB1 motif, JASPAR MA0103.4). **(g)** Multimodal interpretation of rs8052244 (`chr16:87368655:G>C`) in CD14^+^ monocytes at the *FBXO31* locus. Top: Predicted reference (REF) and alternative (ALT) allele profiles for local scATAC-seq and bulk *FBXO31* RNA-seq coverage conditioned on the CD14^+^ monocyte reference condition. Bottom: ISM attribution logos for scATAC-seq accessibility and single-cell *FBXO31* gene expression with matched motifs (right), showing disruption of an ETS motif (orange shading, JASPAR SPI1 MA0080.5) adjacent to a repressive homeodomain motif (JASPAR HHEX MA0183.1). **(h)** Cell-type prioritization for rare non-coding variants in congenital hyperinsulinism. Top: *HK1* locus showing the intronic regulatory region containing *n* = 6 disease-associated *de novo* variants and *n* = 80 common variants. Bottom: Box plots of AG2-predicted variant effects (log fold change) on *HK1* expression for the disease-associated and common variants across seven pancreatic and neuroendocrine primary cell types (darker colors highlight causal *β*-cell conditions). Box plots are defined as in **a**.

These improvements were evident at individual loci on held-out sequences. For the mouse gene *Trpm5*, which is required for chemosensing and is selectively expressed in intestinal tuft cells^35^, AG2 accurately predicted selective *Trpm5* expression and tuft-cell-specific accessible chromatin peaks at the *Trpm5* locus across mouse cell types, whereas AG1-sc failed to resolve either signal (Supplementary Fig. 3b). Similarly, across human hematopoietic differentiation, AG2 recovered continuous gene expression trajectories on held-out sequences for key lineage-defining regulators from hematopoietic stem cells toward erythroid and B-cell lineages with higher fidelity than AG1-sc (Supplementary Fig. 3c).

We ablated AG2 to determine whether these improvements stemmed from the expanded training data or the architectural differences relative to AG1 (Supplementary Fig. 4a,b). On the same genome split, we compared pre-trained AG2 models against an unconditioned AG1 multitask model retrained on matched bulk datasets and against restricted-conditioning AG2 configurations pre-trained on the identical full corpus (Methods). Pre-trained AG2 consistently improved gene expression LFC prediction across cell lines and GTEx tissues and condition-specific peak prediction for histone and TF ChIP-seq (Supplementary Fig. 4b). Restricting where condition embeddings enter the network showed that conditioning only at the output or decoder layers yielded minimal gains over the multitask baseline, whereas conditioning the transformer and encoder drove the primary improvements (Supplementary Fig. 4a,b).

### Improved prediction of non-coding variant effects

We next evaluated whether improved condition-specific track modeling translates to more accurate prediction of genetic variant effects using fine-mapped QTLs on held-out test chromosomes, scoring variants with the full-genome distilled AG2 model (Methods). Across benchmarks, we assessed whether models could distinguish putatively causal fine-mapped variants from matched negative controls (causality task) and predict the direction (sign task) and magnitude (coefficient task) of allelic effects.

On bulk tissue eQTLs from GTEx across 49 tissues, AG2 improved prediction of allelic effect direction and magnitude as well as prioritization of putatively causal variants compared with AG1 and Borzoi^16^ (Fig. 2c, Supplementary Fig. 5a). These gains were concentrated in the high-precision regime most relevant for variant prioritization: for causality prediction, AG2 increased recall at 90% precision from 17% for AG1 to 26% (overall area under the precision–recall curve, AUPRC, 0.74 vs. 0.72 for AG1 and 0.69 for Borzoi), and for sign prediction, it increased recall at 90% accuracy from 32% to 40% (overall area under the sign accuracy–recall curve, AUC, 0.85 vs. 0.83 and 0.77; Fig. 2c). Stratifying variants revealed that AG2’s improvement over AG1 was driven primarily by eQTLs with putative tissue-specific effects (Fig. 2d; Methods), consistent with improved prediction of cross-tissue expression differences carrying over to tissue-specific variant effects. Across the remaining bulk splicing, chromatin accessibility, DNase sensitivity, and polyadenylation QTL benchmarks from AG1^14^, AG2 performed comparably to AG1 (Supplementary Fig. 2).

To evaluate variant effect prediction at the cell-type level, we benchmarked models on single-cell QTL from two studies: TenK10K, which mapped variant effects on expression (sc-eQTLs; 28 cell types) and chromatin accessibility (sc-caQTLs; 24 cell types) across peripheral blood mononuclear cells^7,37^, and IBDverse, which mapped sc-eQTL in intestinal epithelial and immune cells (47 cell types, excluding blood)^38,39^. Both cohorts are scored using matched reference conditions from our training corpus (Methods). Across the causality, sign, and coefficient tasks constructed from fine-mapped variants, AG2 outperformed baseline models on all sc-eQTL tasks and sc-caQTL tasks (Fig. 2e, Supplementary Fig. 5b,c,d). Gains were again largest in the 90% precision and accuracy regime: AG2 increased recall from 16% for AG1-sc to 30% for TenK10K sc-eQTL causality (overall AUPRC 0.76 vs. 0.72 for AG1-sc and 0.69 for Scooby), from 23% to 43% for TenK10K sc-eQTL sign prediction (overall AUC 0.85 vs. 0.79 and 0.77), and from 31% to 45% for IBDverse sc-eQTL sign (overall AUC 0.86 vs. 0.81; Fig. 2e). A full-genome distilled AG2 model trained with all bulk PBMC tracks held out (“AG2 (no bulk PBMC)”; Methods) showed largely unchanged single-cell QTL performance on TenK10K, with a small decrease in effect-size correlation (Spearman *ρ* 0.47 to 0.45; Supplementary Fig. 5e), demonstrating that these predictions do not depend on bulk PBMC training data.

Previous evaluations of S2F models observed a steep decline in eQTL prediction accuracy with distance from the transcription start site (TSS) and concluded that such models largely fail to capture distal regulatory elements^33^. However, stratifying fine-mapped eQTLs by whether they overlap accessible chromatin reveals a sharp distinction between in-peak and outside-peak variants that is masked in unstratified analyses (Supplementary Fig. 6a,b,c). For TenK10K sc-eQTLs located within accessible primary blood scATAC-seq peaks, predictive performance remained strong even at longer distances (>35 kb from the TSS); at these distances AG2 retained a clear advantage over AG1-sc and Scooby in sign prediction (area under the receiver operating characteristic curve, AUROC, 0.86 vs. 0.78 for AG1-sc) and effect-size prediction (Spearman *ρ* 0.57 vs. 0.47; Supplementary Fig. 6a). By contrast, outside accessible peaks, AG1-sc and Scooby predicted sign at close to random chance level, and AG2 retained only modest accuracy (sign AUROC up to 0.63; Supplementary Fig. 5e). Repeating this stratification on GTEx bulk eQTLs using ENCODE candidate *cis*-regulatory elements (cCREs)^40^ as a proxy for accessible chromatin showed a consistent but weaker difference between in-cCRE and outside-cCRE variants (Supplementary Fig. 6b), indicating that S2F models can capture distal regulatory variant effects when variants reside within active regulatory elements. Stratifying IBDverse sc-eQTL within proxy peaks (Methods), yielded similar results (Supplementary Fig. 6c).

To illustrate how cell-type-resolved predictions clarify bulk tissue associations, we examined the fine-mapped GTEx eQTL rs62420789 (`chr6:137026715:G>A`), which increases *IL20RA* expression in lung and adipose tissue but decreases its expression in esophagus mucosa and other tissues (Fig. 2f). AG2 resolved these opposing tissue-level effects into distinct cell-type-specific mechanisms: it predicted that rs62420789 decreases *IL20RA* expression in epithelial cells (such as lung goblet cells), where nucleotide-resolution *in silico* mutagenesis (ISM) indicated disruption of an activating GRHL2 binding motif, but increases expression in endothelial cells (such as lung blood vessel endothelial cells), where ISM indicated disruption of a repressive ZEB1 motif (Fig. 2f). Consistent with these motif attributions, *GRHL2* is selectively expressed in epithelial lineages, where *ZEB1* is selectively absent (Fig. 2f). Because bulk tissue RNA-seq measures the average expression across a mixture of constituent cell types, a variant that acts in opposite directions in epithelial and endothelial cells can yield opposite bulk eQTL signs depending on which cell population dominates a given tissue sample, explaining the apparent tissue-specific reversal across GTEx tissues as a consequence of cellular composition.

Unlike models trained solely on chromatin accessibility, AG2 jointly models chromatin and gene expression to capture variant effects across both layers of regulation simultaneously, even when they act in opposite directions. For example, the fine-mapped QTL rs8052244 (`chr16:87368655:G>C`) in CD14^+^ monocytes is associated with decreased local chromatin accessibility but increased expression of *FBXO31* (Fig. 2g). AG2 recapitulated both effects, and ISM attribution across the two modalities suggested that the variant disrupts an ETS-like motif that establishes local accessibility, whereas this motif, together with an adjacent homeodomain motif, is predicted to drive repression of *FBXO31* (Fig. 2g). Independent macrophage QTL data^41^ further show that the alternate allele is associated with reduced levels of the repressive mark H3K27me3 along the *FBXO31* gene body, consistent with a repressive role for this element.

Finally, we tested whether AG2 can retrospectively prioritize known disease-associated variants and their relevant cell type in a published rare disease locus. In congenital hyperinsulinism, rare *de novo* non-coding variants (*n* = 6) within an intronic regulatory region of *HK1* cause inappropriate insulin secretion by selectively upregulating *HK1* in pancreatic *β* cells^42^ (Fig. 2h). AG2 predicted strong gain-of-expression effects for these individual variants specifically in pancreatic islet *β* cells (islet of Langerhans beta and pancreas beta reference conditions), while predicting near-zero effects in non-endocrine pancreatic lineages, off-target endocrine cells, and *n* = 80 common population variants within the surrounding regulatory region (Fig. 2h). Together, these results show that AG2 can accurately resolve regulatory variant effects and their underlying mechanisms across both bulk tissues and individual cell types.

### Imputing unmeasured epigenomic assays

Chromatin accessibility and histone modifications mark active and repressed regulatory elements across cell types, yet profiling them across hundreds of human tissues and primary cell types is impractical due to cost and sample constraints. As a result, even the largest epigenomic compendia remain incomplete, missing tens of thousands of possible experiments^43^. This gap is widest in single-cell atlases, where most studies profile only scRNA-seq because parallel or joint scATAC-seq is costly. Histone modifications are rarer still, as bulk ChIP-seq requires purified cell populations and single-cell assays capture only one to a few marks per experiment with sparse per-cell coverage and specialized protocols^44^. Because AG2 was trained on bulk and single-cell data together, every assay measured in a condition informs a shared condition embedding. Thus, in principle, the model can impute unmeasured modalities from any combination of observed assays and predict variant effects in unmeasured assays (Fig. 1b; Methods).

To benchmark imputation performance, we trained a full-genome distilled AG2 model (the imputation-split model; Methods) holding out a test set of 184 bulk tracks across 30 conditions, spanning chromatin accessibility, RNA-seq, and ChIP-seq for both TFs and histone modifications (Supplementary Fig. 7a, Supplementary Table 4). The other assays measured in each condition remained in training, reflecting the common situation in which a condition has been profiled with some assays but not others. For a consistent comparison against a baseline method, we retrained Avocado^45^, a tensor factorization method, on the same split after a comprehensive hyperparameter search (Methods). We also compared performance against two baselines that copy signal from training tracks of the same assay: their mean, and an oracle that uses the held-out track to select and average the five most correlated training tracks (closest), thereby approximating an upper bound for transferring signal from similar conditions (Methods). Because most signal is shared across conditions, genome-wide correlation with the observed track largely reflects this shared signal: the mean, which contains no condition-specific information, reached Pearson *r* of 0.47–0.72 across chromatin assays, similar to Avocado (0.39–0.73; Supplementary Fig. 8f). Across the consensus peak catalog for each assay, we therefore scored each region by its mean predicted coverage and evaluated how accurately this classified whether the region carried a called peak in the held-out condition (peak AUPRC; Methods). Peaks shared by most conditions can be predicted by averaging other conditions, so we also stratified consensus peaks by the fraction of conditions in which they are called. For RNA-seq, whose signal is likewise dominated by genes expressed in all conditions, we compared predicted and observed expression differences between held-out conditions (gene log fold changes; Methods).

AG2 recovered held-out peaks more accurately than all three alternatives for each chromatin assay (peak AUPRC of 0.69–0.79; Supplementary Fig. 7b), better predicted expression differences between conditions for RNA-seq (Pearson *r* = 0.74, compared with 0.58 for Avocado and 0.69 for closest; Supplementary Fig. 7b), and outperformed Avocado on 166 of 181 evaluated tracks (Supplementary Fig. 8a). Gains were largest on the rarest peaks (≤12.5% of conditions), where AG2 achieved a 1.7- to 2.9-fold higher AUPRC than the better of Avocado and closest, and negligible on peaks shared across nearly all conditions (Supplementary Fig. 8b–e). For TF ChIP-seq (66 held-out tracks across 43 factors), both Avocado and closest performed near or below random on these rare peaks (Supplementary Fig. 8b). These results are consistent with AG2 inferring condition-specific peaks from DNA sequence rather than copying signal from related conditions, as illustrated by held-out H3K4me1 enhancer peaks at the *LMO1* locus in SK-N-SH cells (Supplementary Fig. 7c). We note that each held-out TF track comes from a condition in which ChIP-seq for that factor was performed, and thus in which the factor is expressed; we expect that imputing binding in conditions where a factor is not expressed or otherwise inactive will yield poor performance with false-positive peaks due to a lack of negative feedback provided to the model.

We then tested whether imputation can annotate enhancers in primary human immune cells profiled only with single-cell assays. Using the imputation-split model, which also held out all bulk PBMC and astrocyte tracks, we imputed H3K27ac across nine PBMC conditions with observed scRNA-seq and scATAC-seq and combined it with observed accessibility to classify open non-promoter representative DNase I hypersensitive sites (rDHSs) with enhancer-like signatures (Methods). Compared with held-out primary PBMC Mint-ChIP H3K27ac profiles^46,47^, imputed H3K27ac identified enhancer-like elements more accurately across all nine cell types (AUPRC 0.89–0.98) than either observed primary scATAC-seq alone (0.62–0.83) or observed H3K27ac ChIP-seq from lineage-matched cell lines (0.76–0.93; Supplementary Fig. 7d), and predicted cell-type differences in H3K27ac across genome-wide rDHSs (*r* = 0.44; Supplementary Fig. 7e). Thus, imputing H3K27ac from single-cell accessibility and expression can help annotate enhancers in primary cell types.

### Generalizing to new cell states

Fixed-track S2F models such as AG1 and CREsted^18^ must be trained or fine-tuned for new datasets and cell states, whereas cell-conditioned models such as Scooby^21^ are trained on individual datasets and do not generalize across all cell types. To generalize well to new samples across human biology without additional training, we introduced a post-training stage that derives AG2 conditions from scRNA-seq profiles (Fig. 3a; Methods). We first trained an scVI-inspired single-cell expression autoencoder^51^ on human scRNA-seq data largely sourced from CELLxGENE^2^ (Methods). We then post-trained the distilled AG2 student model to predict each cell’s autoencoder reconstruction from DNA sequence, conditioned on that cell’s embedding. Because these reconstructions are denoised estimates of expression rather than sparse counts, they serve as pseudobulk-like targets at single-cell resolution, exposing AG2 to the full diversity of profiled human cell states without collapsing cells into discrete pseudobulks. At inference, rather than running the sequence model on every cell, we condition AG2 on a cell population (such as a cell type or metacell) by encoding its cells and averaging their embeddings (Fig. 3d; Methods).

**Figure 3.**
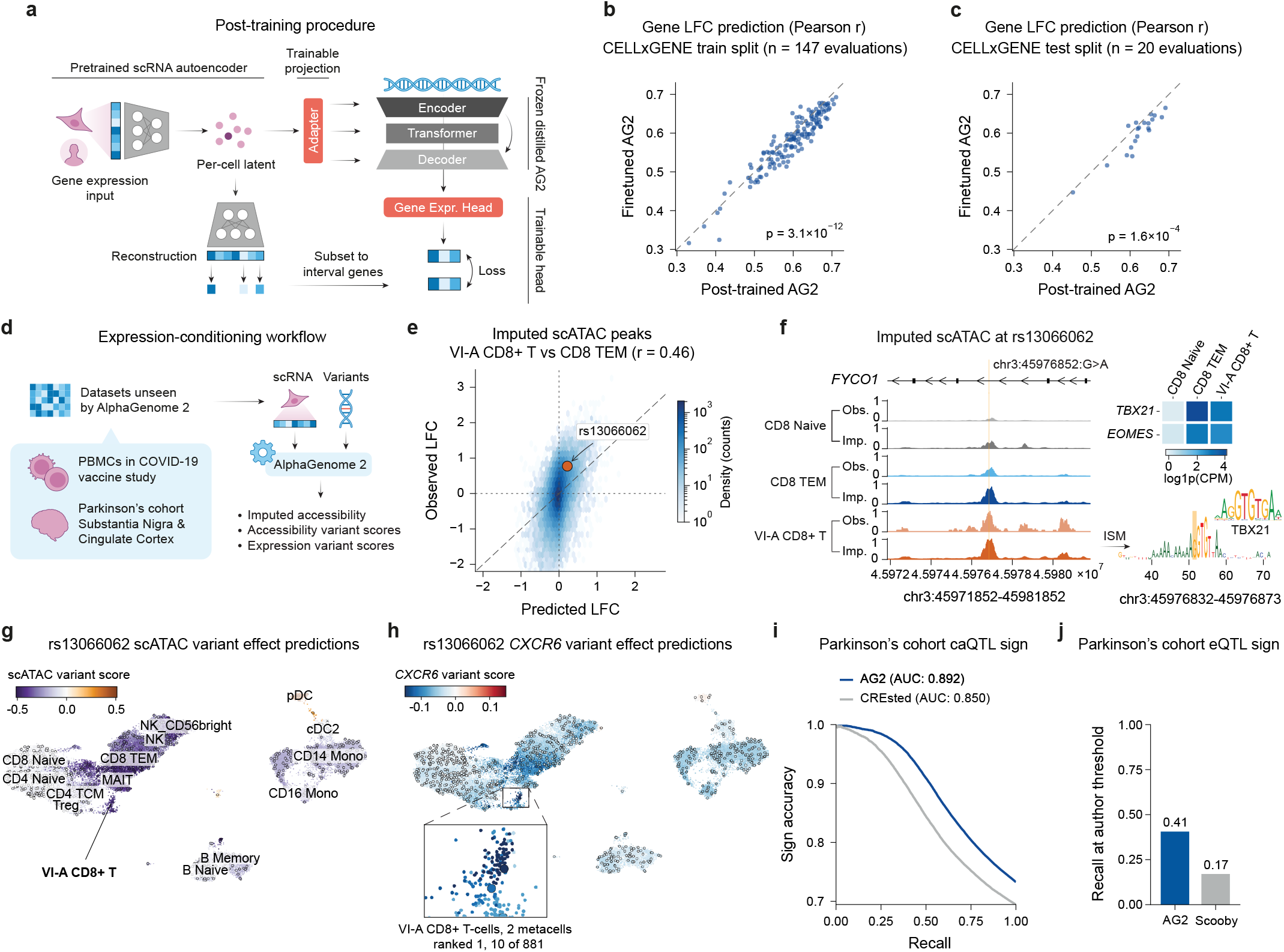
Expression conditioning extends AlphaGenome 2 to unseen single-cell datasets and cell states. **(a)** Post-training procedure (Methods). A pre-trained scRNA-seq autoencoder embeds single cells and reconstructs denoised gene expression. With the autoencoder and distilled AG2 frozen, a trainable adapter maps cell embeddings into AG2’s condition space and a trainable head predicts each cell’s reconstructed expression from DNA sequence. **(b,c)** Cell-type-specific gene expression prediction on held-out genomic intervals comparing post-trained and dataset-fine-tuned AG2 across CELLxGENE evaluation groups from the training split (**b**; *n* = 147) and a chronologically held-out test split (**c**; *n* = 20). Each point shows the mean cross-gene Pearson *r* between predicted and observed LFCs (per cell type relative to the mean) within 5–10 cell types from the same lineage, tissue, and dataset (Methods; dashed line, *y* = *x*; one-sided paired Wilcoxon signed-rank test). **(d)** Expression-conditioning workflow on unseen studies. Single cells from post-vaccination PBMCs^48^ and from substantia nigra and cingulate cortex in a Parkinson’s disease cohort^49^ are embedded with the cell encoder and averaged per cell type or metacell to impute chromatin accessibility and score variant effects on accessibility and expression. **(e–h)** SARS-CoV-2 vaccination PBMC cohort (days 11 and 28 pooled; AG2 conditioned on scRNA-seq, observed accessibility from separate scATAC-seq cells; Methods)^48^. **(e)** Predicted versus observed scATAC-seq LFC between vaccine-induced group A CD8^+^ T (VI-A CD8^+^ T) and effector memory CD8^+^ T (CD8 TEM) cells across predicted accessible peaks (*n* = 37,035; Pearson *r* = 0.46; orange circle, peak containing rs13066062; dashed line, *y* = *x*). **(f)** Left: Observed (Obs.) and imputed (Imp.) scATAC-seq profiles (independently min–max scaled across states) in naive CD8^+^ T (CD8 Naive), CD8 TEM, and VI-A CD8^+^ T cells around rs13066062 (`chr3:45976852:G>A`, orange line) at *FYCO1*. Top right: Observed expression of *TBX21* and *EOMES* (log(CPM + 1); CPM, counts per million). Bottom right: scATAC-seq ISM logo around rs13066062 (orange shading) in VI-A CD8^+^ T cells with the matched TBX21 motif (JASPAR MA0690.1). **(g,h)** Predicted effect of rs13066062 on local scATAC-seq (**g**) and *CXCR6* expression (**h**) across 881 PBMC metacells conditioned on their mean cell embeddings, shown on a uniform manifold approximation and projection (UMAP)^50^ of cell-encoder embeddings (cells colored by metacell score; outlined circles, metacells; inset in **h**, VI-A CD8^+^ T metacells). **(i,j)** Variant effect direction across brain regions and cell types in the Parkinson’s disease cohort^49^ (Methods). **(i)** Sign accuracy versus recall across concordant chromatin accessibility QTL and allele-specific chromatin accessibility variants (caQTL-ASCA; *n* = 86,596) for expression-conditioned AG2 and cohort-trained CREsted. **(j)** Recall of overlapping eQTLs (*n* = 1620) matched at the 87.3% sign accuracy achieved by cohort-fine-tuned Scooby at its *|*LFC*|* > 0.05 reporting threshold.

We first evaluated two aspects of the autoencoder: its cell embeddings and its expression reconstructions. To test the cell embeddings on unseen data, we applied the encoder to PBMC scRNA-seq profiles from a SARS-CoV-2 vaccination study^48^; without dataset-specific training, the embeddings preserved cell-type structure on par with an scVI model trained on this dataset, with similar integration benchmarking scores in this cohort with limited batch effects^52^ (Supplementary Fig. 9a,b@). To test the reconstructions in the realistic setting of newly generated studies, we used a temporal split of CELLxGENE and evaluated across 16 held-out test datasets (Methods). For each cell type, we averaged the embeddings of *k* cells and decoded this mean embedding into a single expression profile (Supplementary Fig. 9c). Decoding a single cell’s embedding (*k* = 1) already recovered observed pseudobulk gene LFCs across cell types (median Pearson *r* = 0.61), confirming that single-cell reconstructions provide effective pseudobulk-like targets for post-training. Moreover, although the autoencoder was trained on individual cells, decoding the mean embedding of an increasing number of cells improved agreement with observed pseudobulk LFCs up to a median *r* of 0.87 across all cells (Supplementary Fig. 9c), suggesting that the embedding space is locally linear and demonstrating that averaging cell embeddings denoises individual cells to represent a cell population at inference.

We next asked whether expression conditioning could replace dataset-specific fine-tuning. Starting from a distilled student model trained with genomic intervals held out (and a cell encoder trained without access to genes in those intervals), we compared post-training against fine-tuning the same student model on pseudobulk expression from each CELLxGENE dataset (Methods). On the held-out intervals, expression conditioning predicted cell-type-specific expression more accurately than fine-tuning across 147 evaluation groups of related cell types from training-split datasets (*P* = 3.1 × 10*^−12^*; one-sided paired Wilcoxon signed-rank test; Fig. 3b) and across 20 evaluation groups from test-split datasets unseen during autoencoder training and post-training (*P* = 1.6 × 10*^−^*^4^; one-sided paired Wilcoxon signed-rank test; Fig. 3c), demonstrating that a cell representation learned across the CELLxGENE corpus shares information across datasets and cell states more effectively than fitting each dataset separately. Post-training this distilled student model also outperformed post-training a single pre-trained model (Supplementary Fig. 9d,e). Expression conditioning also preserved variant effect prediction: conditioning the full-genome AG2 model on scRNA-seq datasets excluded from reference-condition pre-training and distillation^38,53^, we found that expression conditioning matched reference conditioning on sc-eQTL sign prediction (Supplementary Fig. 10a,c) and causality prediction (Supplementary Fig. 10b,d), on both the TenK10K and IBDverse datasets. Additionally, expression-conditioning maintained performance on TenK10K sc-eQTL tasks even when taking subsamples of 128, 16, or even a single cell per cell type (Supplementary Fig. 10e,f).

We then applied expression conditioning with the full-genome AG2 model to two studies unseen by AG2 (Fig. 3d). The first, the SARS-CoV-2 vaccination study above, identified vaccine-induced group A CD8^+^ T (VI-A CD8^+^ T) cells, a rare population of SARS-CoV-2-specific CD8^+^ T cells that expands after booster vaccination^48^. Chromatin accessibility was profiled in only 111 of these cells, separately from scRNA-seq, providing a sparse readout of this state. Conditioned on the mean cell embeddings of each T-cell state from scRNA-seq alone, AG2 imputed accessibility differences between VI-A CD8^+^ T cells and transcriptionally similar effector memory CD8^+^ T (CD8 TEM) cells that correlated with the observed differences (Pearson *r* = 0.46 across 37,035 predicted accessible peaks; Fig. 3e; Methods), indicating that expression conditioning can recover chromatin differences in rare cell states that are difficult to profile directly.

Zhang *et al.* reported that regulatory elements with increased accessibility in VI-A CD8^+^ T cells harbor variants associated with COVID-19 severity, including near *FYCO1* at the 3p21.31 locus, but did not link these variants to target genes^48^. One such variant, rs13066062 (`chr3:45976852:G>A`), lies in a *FYCO1* intron within a peak that was more accessible in VI-A CD8^+^ T cells than in CD8 TEM cells in both observed and imputed data (Fig. 3e,f). ISM in VI-A CD8^+^ T cells indicated that the variant disrupts a T-box motif compatible with TBX21 and EOMES, which are expressed at higher levels in CD8 TEM and VI-A CD8^+^ T cells than in naive CD8^+^ T cells (Fig. 3f). Consistent with the expression of these factors, conditioning AG2 on 881 metacells spanning all post-vaccination PBMCs predicted loss of accessibility concentrated in CD8 TEM, natural killer (NK), and VI-A CD8^+^ T cells, with little effect in naive T, B, and myeloid cells (Fig. 3g; Methods). AG2 also imputed higher H3K27ac at this peak in CD8 TEM and VI-A CD8^+^ T cells than in naive CD8^+^ T cells and predicted that rs13066062 modestly reduces H3K27ac in these states (Supplementary Fig. 10k). Across nearby genes, the variant was predicted to decrease expression of the chemokine receptor gene *CXCR6* more strongly than *FYCO1* (Supplementary Fig. 10i); this effect was concentrated in effector T-cell and NK metacells, with the two VI-A CD8^+^ T metacells ranking first and tenth of 881 (Fig. 3h, Supplementary Fig. 10j). These predictions nominate *CXCR6* as a putative target of rs13066062, aligning with 3p21.31 colocalization and eQTL analyses linking COVID-19 severity risk to reduced *CXCR6* expression in T cells^54^.

The second unseen study profiled the substantia nigra and cingulate cortex of 190 donors with and without Parkinson’s disease by single-nucleus multiome sequencing (paired RNA-seq and ATAC-seq), mapped eQTLs and chromatin accessibility QTLs (caQTLs) alongside allele-specific chromatin accessibility (ASCA), and trained CREsted and fine-tuned Scooby models on these data^49^. Conditioned on the mean single-nucleus expression embeddings of each cell type and brain region, AG2 predicted the effect direction of variants significant in both accessibility analyses (caQTL-ASCA) more accurately than CREsted (sign accuracy–recall AUC 0.892 vs. 0.850; Fig. 3i), and at the sign accuracy Scooby achieves at its reporting threshold for eQTLs (87.3%), AG2 reached more than double the recall (41% vs. 17%, Fig. 3j). On both tasks, expression conditioning also matched or slightly exceeded conditioning on matched reference conditions (caQTL-ASCA AUC 0.892 vs. 0.885; eQTL AUC 0.834 vs. 0.807; Supplementary Fig. 10g,h). Thus, without any training on this cohort, post-trained AG2 outperformed sequence models trained on the cohort’s own data, demonstrating that expression conditioning extends AG2 to unseen datasets without fine-tuning.

### Predicting enhancer–gene links at scale

Mapping distal enhancers to their target genes across human cell types remains challenging: CRISPR interference (CRISPRi) screens are largely limited to immortalized cell lines^11,55,62^, peak–gene correlation methods^63^ require matched chromatin accessibility and expression data and confound regulation within a cell type with co-variation across cell states^64^. Supervised classifiers such as ENCODE-rE2G(-Extended) and scE2G require experimental chromatin assays in each target cell type^55,64^. S2F models instead estimate enhancer importance from the sensitivity of predicted gene expression to the input DNA sequence^14,16^, but prior models are restricted to fixed and mostly bulk training tracks. Because AG2 is trained across hundreds of primary single-cell reference conditions and supports expression conditioning on new scRNA-seq profiles, it can predict cell-type-specific E2G links across primary cell types and unseen single-cell datasets without requiring chromatin assays in the target cell type. To score candidate E2G links in a target condition, we took the gradient of the model’s predicted gene expression with respect to the 1-Mb input sequence centered on each gene’s TSS and aggregated the unnormalized gradient magnitudes across each candidate element (Fig. 4a; Methods).

**Figure 4.**
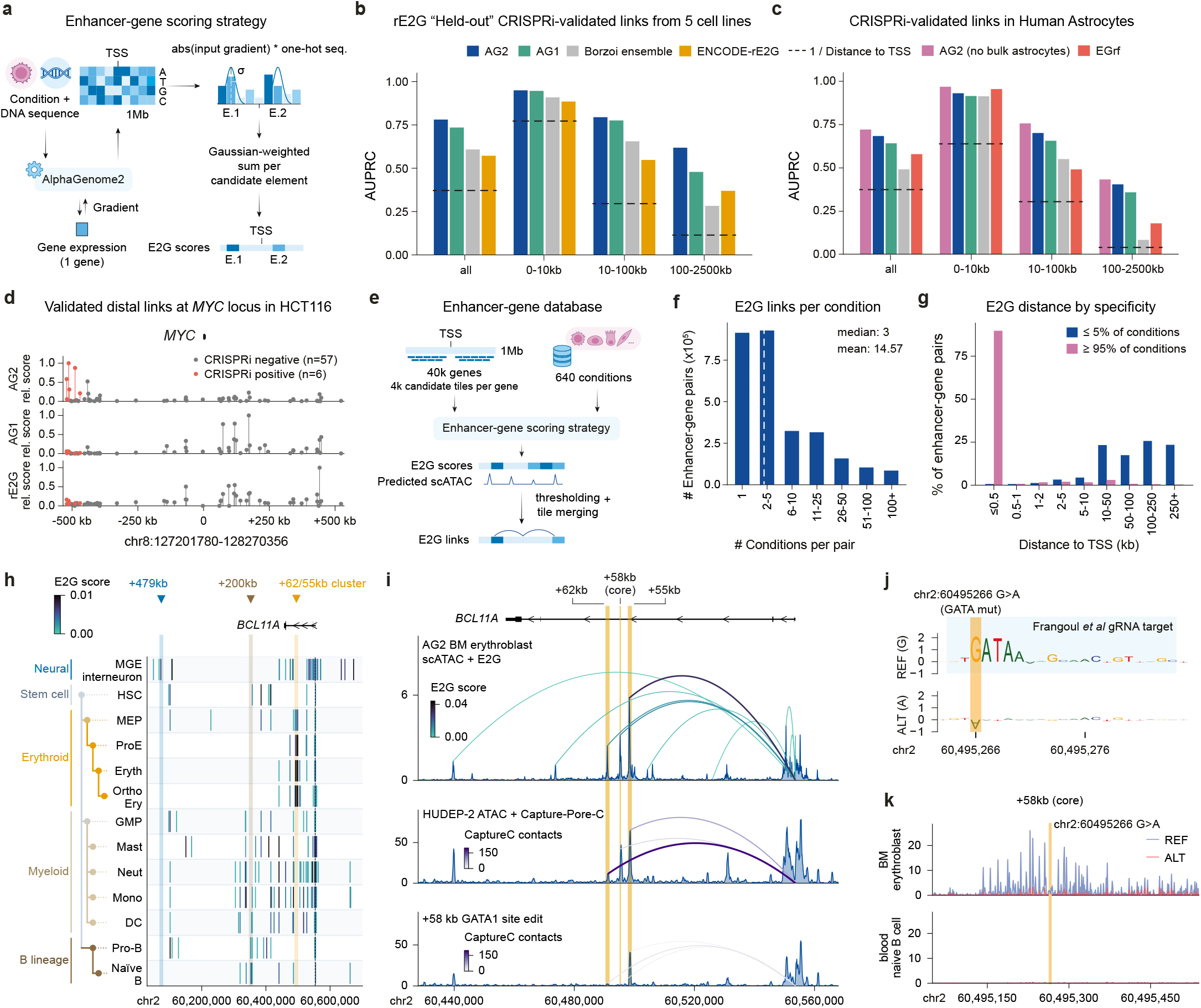
AlphaGenome 2 links enhancers to their target genes in primary cell types. **(a)** E2G scoring strategy (Methods). Conditioned on a target cell type, candidate elements in a 1-Mb sequence centered on a gene’s TSS are scored from the gradient of predicted expression with respect to the input sequence. **(b,c)** E2G linking AUPRC across all element–gene pairs and by TSS distance on **(b)** the held-out ENCODE-rE2G CRISPRi benchmark across five cell lines^55^ (*n* = 4378 pairs, 190 positives; Methods) and **(c)** a human astrocyte CRISPRi screen^56^ (*n* = 1902 pairs, 122 positives), comparing AG2, AG1^14^, a Borzoi ensemble^16^, ENCODE-rE2G (**b**), EGrf (**c**) and AG2 trained without bulk astrocyte tracks (**c**). **(d)** E2G scores normalized to the window maximum for AG2, AG1 and ENCODE-rE2G across *n* = 63 CRISPRi-tested elements (*n* = 6 positives) at the *MYC* locus in HCT116 cells. **(e)** Genome-wide E2G atlas pipeline across 640 primary single-cell reference conditions (Methods). **(f)** Consensus distal E2G pairs by number of active conditions across the 640 reference conditions (dashed line, median). **(g)** TSS distance distribution for condition-specific (*≤*5% of conditions) versus ubiquitous (*≥*95%) consensus E2G pairs. **(h)** Active E2G links across the *BCL11A* locus in 13 primary neural and hematopoietic conditions (500-bp bins colored by gradient score; dashed line, TSS; Methods). Lineage-colored shading and arrowheads mark the +62/55 kb erythroid enhancer cluster^57,58^ and B-cell (+200 kb^58^) and cortical (+479 kb^59^) promoter capture Hi-C contacts. **(i)** Zoomed view of *BCL11A* showing AG2-predicted bone marrow (BM) erythroblast scATAC-seq and active E2G links (top) alongside experimental ATAC-seq and enhancer-probe Capture-Pore-C in wild-type (middle) and `sg1617`-edited (+58 kb GATA1 site; bottom) HUDEP-2 erythroid cells^60^. Experimental arcs show TSS contacts per million genome-wide paired-end tags per baited enhancer (mean of two replicates; Methods); orange shading marks the +62, +58 (core) and +55 kb elements. **(j)** ISM logos of predicted BM erythroblast scATAC-seq across the therapeutic guide RNA (gRNA) target site^61^ for REF and the GATAA-disrupting ALT allele (`chr2:60495266:G>A`; orange shading). **(k)** Base-resolution predicted scATAC-seq across the +58 kb core enhancer for REF and ALT in BM erythroblasts and blood naive B cells (orange shading as in **j**).

Using the distilled AG2 model trained across the full genome, we first evaluated E2G linking on the held-out ENCODE-rE2G CRISPRi benchmark spanning 4378 element–gene pairs across five cell lines outside the ENCODE-rE2G training set^55^. Conditioned on the matching reference conditions, AG2 achieved an AUPRC of 0.78, outperforming AG1 (0.73), a four-model Borzoi ensemble (0.61) and ENCODE-rE2G (0.57; Fig. 4b; K562 benchmark in Supplementary Fig. 11f). Whereas sequence models performed comparably within 10 kb of the TSS (AUPRC 0.91–0.95), AG2’s advantage grew with distance, reaching an AUPRC of 0.62 at 100–2500 kb versus 0.48 for AG1, 0.37 for ENCODE-rE2G and 0.28 for Borzoi (Fig. 4b). On a CRISPRi screen in cultured primary human astrocytes^56^ AG2 achieved an AUPRC of 0.68, surpassing AG1 (0.64), Borzoi (0.49) and the screen’s random-forest predictor EGrf (0.58), again with the largest gain at 100–2500 kb (0.40 versus 0.18 for EGrf; Fig. 4c). Across all three CRISPRi benchmarks, omitting the window-wide attribution normalization used in prior work^14,16^ consistently improved accuracy across sequence models (Supplementary Fig. 11a–c). This performance did not depend on bulk astrocyte data: the imputation-split distilled model, in which all bulk astrocyte tracks were held out so that the astrocyte reference condition was trained only on the screen’s scRNA-seq pseudobulk, reached an AUPRC of 0.72 (Fig. 4c; Methods), and expression conditioning on K562 and astrocyte scRNA-seq profiles matched or exceeded standard reference conditioning (Supplementary Fig. 11d,e). Single-cell expression alone can therefore suffice to map E2G links in primary cells. Likewise, in the unseen Parkinson’s disease brain cohort^49^, expression-conditioned links at caQTL-ASCA peaks showed high Jaccard similarity specifically with predictions from the matching brain reference conditions (Supplementary Fig. 11g).

To illustrate AG2’s gain on long-range enhancers, we examined *MYC* in HCT116 colorectal carcinoma cells from the held-out benchmark. In colorectal carcinoma, *MYC* is regulated across a gene desert by a lineage-specific super-enhancer 500 kb upstream that transcribes *CCAT1* and loops to the *MYC* promoter^65^. All six CRISPRi-validated enhancers among the 63 tested elements at this locus lie 470–521 kb upstream within or adjacent to this super-enhancer (Fig. 4d). AG2 ranked five of these six enhancers among the top seven elements overall (ranks 1, 2, 3, 5, 7 and 24), whereas AG1 (ranks 24–53) and ENCODE-rE2G (ranks 14–51) missed this upstream cluster and prioritized non-functional elements downstream of *MYC* (Fig. 4d).

We next applied AG2 across 640 primary single-cell reference conditions to build an atlas of E2G links spanning 39,323 protein-coding and long non-coding RNA genes (Fig. 4e; Methods). Using thresholds on AG2-predicted target gene expression, local chromatin accessibility, and expression gradient magnitude chosen on the K562 and astrocyte CRISPRi screens to retain nearly all validated enhancers and target genes while maximizing linking *F*_1_ on the astrocyte screen (Supplementary Fig. 12; Methods), we called and summit-merged active 500-bp genome tiles located more than 500 bp from the TSS. This yielded a median of 64,800 distal links per condition at a median TSS distance of 35.7 kb (mean 81 kb), and a median of 4 distal enhancers per regulated gene (Supplementary Fig. 13a–h). When reconciled across conditions into consensus pairs (Methods), distal links proved highly condition-specific: the median number of active conditions per pair was 3, and 65.1% of links were detected in 5 or fewer conditions (90.0% in ≤5% of conditions; Fig. 4f). Including promoter-proximal elements (≤500 bp from the TSS) showed that ubiquitously active links (≥95% of conditions) were predominantly promoter-proximal (89.8%), whereas condition-specific links (≤5% of conditions) were more distal (99.4% >500 bp; Fig. 4g). Consistently, ubiquitously expressed genes almost always had active promoter-proximal elements and, compared with condition-restricted genes, more frequently lacked linked distal enhancers (Supplementary Fig. 13i).

Querying this atlas for a target gene resolves its lineage-specific regulatory architecture, which can then be dissected at base resolution with AG2. We illustrated this at *BCL11A*, a transcription factor whose expression across erythroid, lymphoid, and neural lineages is governed by distinct distal enhancers^57–59^. Across 13 hematopoietic and neural cell types, the atlas recovered lineage-specific sets of distal enhancers that recapitulate experimentally mapped promoter contacts up to nearly 500 kb away, including a +200 kb contact mapped in naive and total B cells, which among the conditions shown AG2 links most strongly in naive B cells, and a +479 kb prefrontal cortex contact in cortical interneurons (Fig. 4h). In erythroid cells, the highest-scoring links converged on the +62/55 kb intronic enhancer cluster (comprising the +62, +58 and +55 kb elements) that drives erythroid *BCL11A* expression to silence fetal hemoglobin^57^. In HUDEP-2 erythroid cells, these elements are accessible and contact the *BCL11A* promoter, and CRISPR–Cas9 editing of the +58 kb enhancer GATA1 site reduces both^60^ (Fig. 4i; Methods). Within the +58 kb enhancer, clinical CRISPR–Cas9 gene therapy for sickle cell disease and *β*-thalassemia disrupts an essential GATA1 motif (GATAA) to reactivate fetal hemoglobin while sparing non-erythroid *BCL11A* function^57,61,66^. *In silico* mutagenesis across the therapeutic guide RNA target site recovered this GATAA motif as the primary sequence determinant of local accessibility in bone marrow erythroblasts, and a single-nucleotide substitution in the motif (`chr2:60495266:G>A`) removed its attribution (Fig. 4j) and reduced predicted accessibility across the +58 kb enhancer in erythroblasts, whereas naive B cells showed little predicted accessibility there for either allele (Fig. 4k).

### Interpreting complex-trait variants across cell types

Genome-wide association studies have identified thousands of loci associated with complex traits and diseases, yet interpreting a non-coding association requires answering three questions that are difficult to resolve at scale: which variant among those in LD is causal, which gene it regulates, and in which cell type it acts^69,70^. Because AG2 predicts cell-type-resolved gene expression across a 1-Mb sequence window, scoring candidate variants with high confidence across primary cell types can address all three questions simultaneously. To retrospectively test this capability at an experimentally dissected disease locus, we examined the 21q22 intergenic locus associated with inflammatory bowel disease (IBD) and other immune-mediated disorders, where Stankey *et al.* combined chromatin conformation capture, CRISPR–Cas9 perturbation, and MPRAs in primary human macrophages to show that rs2836882 drives disease risk by upregulating the distal gene *ETS2*, located approximately 290 kb away, rather than closer candidate genes^67^ (Fig. 5a). Scoring the seven variants in the fine-mapped IBD credible set across primary immune cell types of the ileum, AG2 recovered all three components of this mechanism: it predicted a substantial effect for only rs2836882, linked the variant selectively to *ETS2* despite its 290-kb distance, and confined the effect to monocytes and macrophages (Fig. 5b).

**Figure 5.**
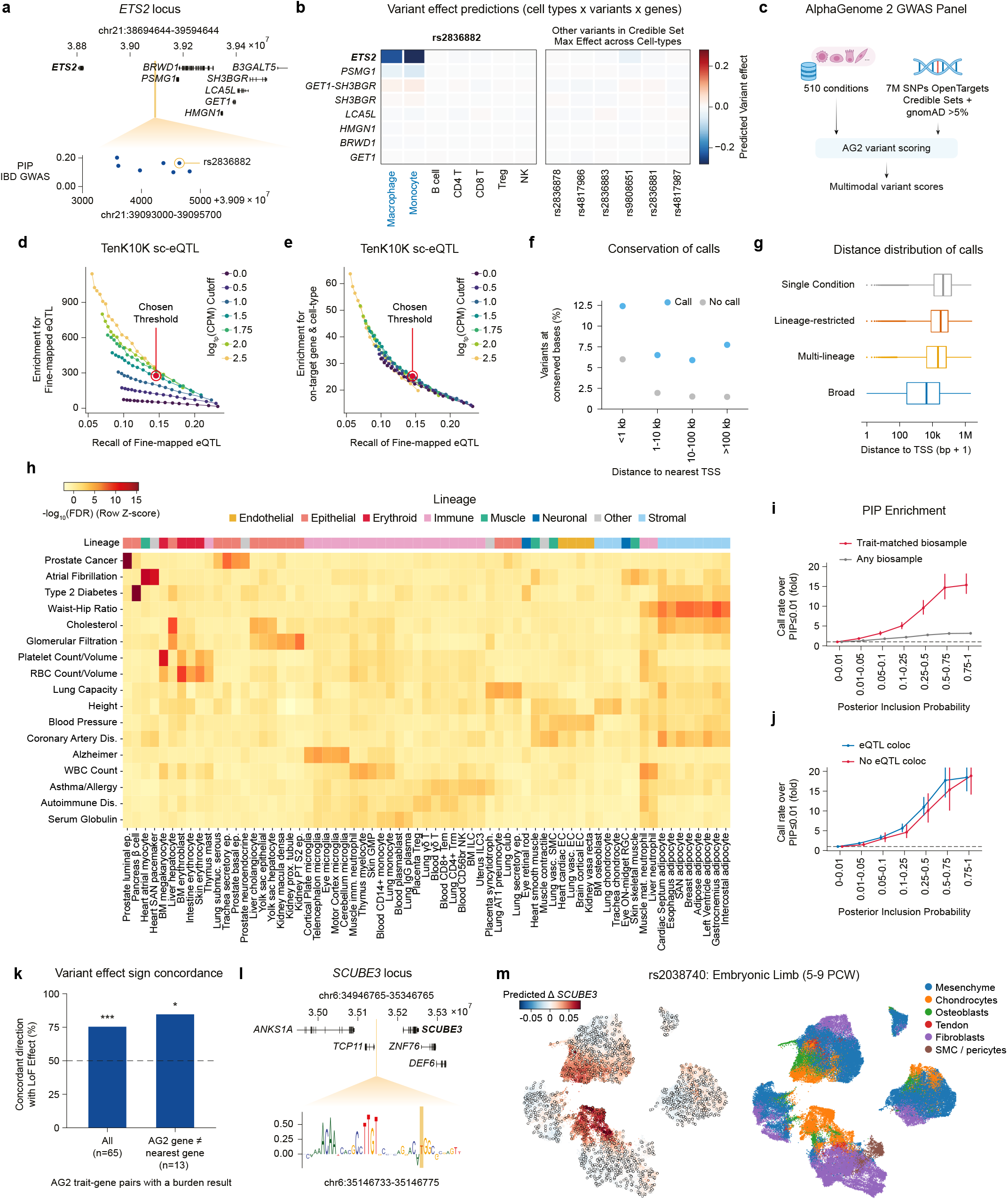
AlphaGenome 2 links non-coding GWAS variants to target genes, cell types and mechanisms. **(a)** Inflammatory bowel disease (IBD) GWAS credible set (*n* = 7 variants) in an intergenic region at 21q22^67^. Top: 900-kb window showing nearby protein-coding genes. Bottom: Zoomed 2.7-kb window showing posterior inclusion probabilities (PIPs), with the experimentally validated causal variant rs2836882 (`chr21:39094644:G>A`) circled. **(b)** (Left) AG2-predicted variant effects (log fold change) on nearby protein-coding genes across primary ileum immune cell types for rs2836882. (Right) Predicted signed effects in ileum immune cells for the remaining six variants in the credible set, computed using the maximum absolute effect per gene. Experimentally validated gene and variant in bold, experimentally validated lineage in blue. **(c)** Overview of the AG2 GWAS panel (Methods). AG2 scores 6,822,130 single-nucleotide variants (SNVs; from Open Targets fine-mapped GWAS credible sets and common gnomAD variants with minor allele frequency (MAF) > 5%) across 510 primary-cell conditions to produce multimodal variant effect scores. **(d)** Calibration of AG2 variant–gene call thresholds using TenK10K single-cell eQTLs across 26 peripheral blood immune conditions^7^ (Methods). Fold enrichment of AG2 calls for fine-mapped eQTLs (PIP > 0.9, *n* = 6362 SNVs) relative to unassociated common background variants (nominal *P* > 0.05, *n* = 1,235,667 SNVs), versus recall of fine-mapped eQTL. Each curve represents one cutoff on the minimum predicted gene expression (log(CPM + 1)) across minimum predicted absolute variant effect cutoffs (*|*LFC*| ∈* [0.05, 0.20]). Red dots mark the operating threshold used throughout (*|*LFC*|* > 0.1, log(CPM + 1) > 1.5). **(e)** Fold enrichment of AG2 calls for on-target (*n* = 14,010) over off-target (nominal *P* > 0.05) pairs of gene and cell type, versus recall of fine-mapped eQTL. Curves and red dot as in **d**. **(f)** Percentage of AG2 called (*n* = 134,813) and uncalled (*n* = 6,687,317) variants which are conserved (phyloP *≥* 2.27, Zoonomia 241 mammal alignment), stratified by distance to the nearest protein-coding transcription start site (TSS). **(g)** Distance to the target gene’s TSS for AG2 variant–gene calls (*n* = 152,849), stratified by cell-type specificity (Methods): single condition (*n* = 65,420), lineage-restricted (1 lineage cluster; *n* = 40,362), multi-lineage (2–3 lineage clusters; *n* = 31,729) and broad (*≥*4 lineage clusters; *n* = 15,338). Box plots display the median (center line), the 25^th^ and 75th percentiles (box bounds), and the whiskers extend to 1.5 times the interquartile range from the box bounds; points beyond whiskers indicate outliers. **(h)** Condition-specific enrichment of GWAS credible-set SNVs relative to common background variants across 17 traits with at least one enriched condition (false discovery rate, FDR *≤* 10*^−^*^7^, one-sided Fisher’s exact test, Benjamini–Hochberg corrected; Methods). Heatmap shows row *z*-scored *−* log­(FDR) for the union of the five most enriched conditions per trait (FDR < 0.05). **(i,j)** Fold enrichment of the AG2 call rate per PIP bin relative to variants with PIP *≤* 0.01 across the 17 traits (*n* = 1,333,700 trait–SNV pairs; Methods). **(i)** Calls in the trait-matched (most significantly enriched) condition per trait versus any of the 510 conditions. **(j)** Calls in the trait-matched condition stratified by whether the variant’s credible set colocalizes with an Open Targets eQTL or sc-eQTL (COLOC *H*­ *≥* 0.8) or not. Dashed lines, fold = 1; error bars, 95% confidence intervals from a 1-Mb genomic block bootstrap. **(k)** Combining the sign of the AG2-predicted expression effect with the GWAS association direction (GWAS *β*) predicts the direction of increased gene expression on the trait, which is compared with the opposite of the rare variant loss-of-function (LoF) burden effect. Percentage of AG2-nominated trait–gene pairs concordant with the LoF burden direction across all pairs with a burden test result (*n* = 65; 75.4%, *P* = 5.1 *×* 10*^−^*^5^) and pairs in which the AG2-nominated gene is not the nearest protein-coding gene (*n* = 13; 84.6%, *P* = 0.022). Dashed line, 50%; *^***^P* < 0.001, *^*^P* < 0.05, two-sided exact binomial test vs. 50%. **(l)** Dissection of the standing-height GWAS variant rs2038740 (`chr6:35146765:C>T`) at the *SCUBE3* locus. Top: 400-kb window showing rs2038740 near *TCP11*, 67 kb upstream of *SCUBE3*. Bottom: *In silico* mutagenesis (ISM) attribution analysis of predicted *SCUBE3* expression change (mean-centered log fold change^14^) in adult bronchus chondrocytes around rs2038740. The alternate allele (rs2038740-T, shaded in orange) is shown. **(m)** Left: Predicted effect of rs2038740 on *SCUBE3* expression (log fold change) for 1575 metacells from the human embryonic limb lateral plate mesoderm (5–9 post-conception weeks (PCW); *n* = 100,935 cells)^68^, each conditioned on the mean embedding of its cells. Cells are shown on a UMAP of cell-encoder embeddings and colored by their metacell’s prediction; dots, metacell mean embeddings. Right: UMAP colored by cell-type labels.

To apply this approach systematically, we used the distilled AG2 model trained across the full genome to score approximately 7 million variants, comprising fine-mapped GWAS credible sets from the Open Targets Platform^27,71^ and common population variants (minor allele frequency > 5%) from gnomAD^72^, across 510 conditions spanning tissue and cell-type combinations (Fig. 5c; Methods). To define a stringent threshold for calling expression-altering variants alongside their target genes and cell types, we calibrated predictions against fine-mapped sc-eQTLs from the TenK10K study^7^, evaluating both enrichment for fine-mapped eQTLs over tested common variants and discrimination of on-target effects from off-target pairs of gene and cell type where the variant had no detectable effect (Methods). Requiring both a minimum predicted effect size and predicted target gene expression in the given condition yielded an approximately 280-fold enrichment for fine-mapped sc-eQTLs and an approximately 25-fold enrichment for on-target over off-target effects (Fig. 5d,e). Variants called at this threshold were predominantly cell-type- or lineage-specific (Supplementary Fig. 14a), overlapped ENCODE cCREs (Supplementary Fig. 14b), and were two- to five-fold more likely than uncalled variants to occupy evolutionarily constrained bases (phyloP ≥ 2.27; Fig. 5f), with more lineage-restricted calls acting at greater distances from the target gene TSS (Fig. 5g). The chosen threshold is likely conservative, because unassociated background variants vastly outnumber fine-mapped eQTLs, so even a small fraction of true regulatory variants missed due to finite sc-eQTL mapping power can deflate the observed enrichment.

We next asked whether variant calls in each condition recover the cell types and states relevant to individual complex traits. Pooling credible-set variants for well-powered traits and testing for enrichment relative to background common variants across all 510 conditions (Methods) identified at least one significantly enriched condition (false discovery rate 10*^−^*^7^; one-sided Fisher’s exact test; Benjamini–Hochberg correction) for 17 (of 28) traits (Fig. 5h). The prioritized cell types recapitulated established trait biology, including implicating prostate luminal epithelial cells in prostate cancer, cardiomyocytes and sinoatrial node pacemaker cells in atrial fibrillation, microglia in Alzheimer’s disease, and diverse immune lineages across autoimmune diseases (Fig. 5h). Across these 17 traits, the fraction of variants called by AG2 is higher for variants with higher posterior inclusion probability (PIP), confirming that the model preferentially prioritizes putatively causal variants over linked bystanders (Fig. 5i). This relationship was substantially stronger within each trait’s top-enriched condition than when aggregating calls across all conditions (Fig. 5i), and held equally for credible sets that do not colocalize with known eQTLs (Fig. 5j), indicating that AG2 provides information beyond existing eQTL studies. Moreover, even when collapsed across conditions and genes to match cell-type-agnostic complex-trait benchmarks such as TraitGym^73^ (Methods), AG2 outperformed AG1^14^ and matched the conservation-based predictor GPN-Star^74^, while a simple combination of AG2 and GPN-Star outperformed all individual methods (Supplementary Fig. 14c).

Beyond nominating causal variants, target genes, and cell types, AG2 predicts the direction of variant-induced expression changes. Combining the predicted expression sign with the direction of the GWAS trait association yields a directional prediction for how increased gene dosage affects the phenotype, which should be opposite to the effect of coding loss-of-function (LoF) variants identified by rare-variant burden tests (Methods). Across nominated trait–gene pairs with rare-variant burden evidence, the direction predicted by AG2 was concordant with LoF burden evidence in approximately 75% of all pairs and in 85% of pairs where the nominated gene was not the nearest gene to the variant (Fig. 5k).

To illustrate how AG2 links non-coding trait variants to distal target genes and developmental cell states, we examined the height-associated variant rs2038740, which lies proximal to the testis-specific gene *TCP11*. Rather than implicating *TCP11*, AG2 predicted that rs2038740 upregulates *SCUBE3*, located 67 kb away, in adult chondrocytes, where the trait-increasing T allele introduces positive sequence attribution immediately adjacent to an intact SOX transcription factor binding motif (Fig. 5l). Because biallelic LoF variants in *SCUBE3* cause a human skeletal dysplasia with short stature and *Scube3*-deficient mice exhibit impaired endochondral bone growth^75^, increased *SCUBE3* expression in chondrocytes provides a plausible mechanism for increased height. Conditioning AG2 on single-cell expression profiles from embryonic human limb development^68^, where longitudinal bone growth is established, localized the predicted effect of rs2038740 to condensed mesenchyme and differentiating chondrocytes (Fig. 5m). Although *SCUBE3* is broadly expressed across early limb-bud populations, the predicted variant effect tracked the expression of *SOX9*, a master regulator of chondrogenesis^76^ (Supplementary Fig. 14d, *ρ* = 0.47). Additional examples of locus-level variant interpretation across complex diseases are provided in Supplementary Note 1.

## Discussion

We have shown that modulating sequence representations early and throughout the network with a target condition, together with training across bulk and organism-wide single-cell atlases, allows AlphaGenome 2 to better resolve regulatory differences across tissues and closely related primary cell types. AlphaGenome 2 improves condition-specific variant effect prediction on gene expression and E2G linking, and enables cross-modality epigenomic imputation, including in unseen cell states profiled only by scRNA-seq. Across hundreds of human cell types, we applied these capabilities to construct precomputed atlases of distal E2G links and variant scores for nearly 7 million trait-associated and common variants, nominating putatively causal variants, target genes with direction of effect, and affected cell states even at GWAS loci lacking known eQTL colocalizations^8^.

Despite these gains, important limitations remain. At high precision, AlphaGenome 2 recovers only a minority of fine-mapped eQTLs and putatively causal GWAS variants. Even in relevant cell states, AlphaGenome 2 can underestimate causal variant effects, particularly for variants outside accessible chromatin, at very distal elements, or acting through post-transcriptional mechanisms. Low recall also limits gene expression prediction across individuals^77–79^. Coverage and resolution across cell states remain incomplete: transient, stimulated, and disease states are underrepresented in training and in the precomputed GWAS panel^80–82^. The model can still struggle to distinguish fine-grained states such as neuronal subpopulations. Because condition embeddings are learned from observational, steady-state profiles, they reflect correlational rather than causal relationships, and it remains unclear whether AlphaGenome 2 can predict regulatory responses to gene knockout, overexpression, or dynamic cell-state transitions.

These limitations present opportunities to improve next-generation models. Closer analysis of why S2F models struggle outside accessible chromatin, and to what extent this gap reflects eQTL fine-mapping errors, could reveal regulatory layers missed by standard chromatin and expression assays. Adding more modalities, such as single-molecule footprinting assays including Fiber-seq^83,84^, could improve low variant recall by covering more biological mechanisms. Cell-state specificity could be improved by expanding the training data, for example to deeply sampled brain atlases^85–87^. Training across additional species could broaden application scope and improve predictive accuracy, as could training on perturbation data, both in *cis* and in *trans*^88^. Therefore, given the flexible architecture of AlphaGenome 2, improvements could be made by extending the training data to additional modalities, cell types, species, or perturbation experiments.

Beyond expanding training data, AlphaGenome 2 provides a modular foundation for downstream extensions. Because expression conditioning enters the frozen sequence model through a lightweight adapter, analogous adapters could map spatial transcriptomics^89^, natural-language descriptions^90,91^, or experimental perturbation profiles into the same condition space. Furthermore, attribution methods^92^ that resolve sequence motifs could be linked to the expression of transcription factors to construct gene regulatory networks underlying scRNA-seq atlases. Although AlphaGenome 2 requires a separate model prediction per condition and is therefore more computationally intensive than fixed-track multitask models, fast attribution and mutagenesis approaches such as Multi-ISM^93^ could scale variant interpretation^94^ across rare variants^95^ and thousands of cell states, including transient and disease states. As single-cell and spatial profiling continue to expand across human development, tissues, and clinical cohorts^1,2^, where sample sizes rarely suffice for direct QTL mapping, AlphaGenome 2 provides a versatile computational approach for bridging single-cell atlases and human genetics.

## Methods

### Terminology

We use the following terms throughout the Methods.

*AG2 and AG1:* Abbreviations for AlphaGenome 2 (this study) and AlphaGenome 1^14^, respectively.

*Condition:* The tissue, cell line, cell type, or cell state for which AG2 makes a prediction. A condition is supplied to the model either as a reference condition or through expression conditioning.

*Reference condition:* A condition defined by a composite ontology identifier (Ontology organization) and repre-sented by a learned entry in the model’s dictionary of reference condition embeddings.

*Expression conditioning:* Conditioning AG2 on cell embeddings obtained by passing single-cell RNA-seq (scRNA-seq) expression profiles through the cell encoder, typically averaged over a group of cells.

*Output head and channel:* AG2 has 14 output heads in Stages 1–3, plus a denoised single-cell gene expression head added in Stage 4 (Output heads). Each head predicts a fixed set of channels, defined by technical or target attributes (such as assay chemistry, suspension type, sequencing endedness, strand or ChIP target) and shared across all conditions.

*Modality:* A class of molecular readout (for example, chromatin accessibility, gene expression, histone modification, CpG methylation or chromatin contact maps), each predicted by one or more output heads.

*Imputation:* Predicting a track for a condition–channel pair that was not observed during training.

### Training data acquisition and processing

AG2 was trained on bulk functional genomics datasets from the AG1 corpus^14^ together with newly curated bulk DNA methylation, scRNA-seq, and single-cell ATAC-seq (scATAC-seq) datasets from public repositories (Supplementary Table 2). For datasets inherited from AG1, we summarize key processing steps and modifications below; full upstream details are given in Avsec *et al*.^14^.

#### Ontology organization

Biological conditions across all experimental assays and species are formalized using a unified multi-ontology indexing scheme. Each biological condition is uniquely represented by a composite 5-tuple of compact uniform resource identifiers (CURIEs):

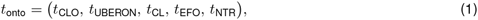

spanning five standard biological ontologies:

- *t*_CLO_: Cell Line Ontology (CLO) identifiers, capturing immortalized and transformed cell lines.
- *t*_UBERON_: Uber-anatomy ontology (UBERON) identifiers, capturing tissues, anatomical systems, and organ structures.
- *t*_CL_: Cell Ontology (CL) identifiers, capturing primary cell types and defined differentiation stages.
- *t*_EFO_: Experimental Factor Ontology (EFO) identifiers, also capturing cell lines.
- *t*_NTR_: New Term Request (NTR) identifiers, accommodating cell states lacking established ontology entries.

Data derived from single-cell modalities, CpG methylation, and primary-cell bulk experiments typically use a combined UBERON term and a CL term, whereas bulk tissue and cell-line conditions typically use a single UBERON, CLO, or EFO identifier.

The composite ontology tuple *t*_onto_ groups experimental datasets that profile the same reference condition under a single shared representation, so a single reference condition may be associated with multiple observed modalities. Within a modality, experiments sharing *t*_onto_ are grouped according to the output channels of that modality’s head (Output heads, Supplementary Tables 5 and 6): experiments differing in channel-defining technical or target attributes map to separate output channels under the same *t*_onto_, whereas replicate experiments sharing both *t*_onto_ and output channel are averaged into a single target track following AG1^14^ (with paired-end and single-end bulk RNA-seq, ATAC-seq, DNase-seq, and ChIP-seq experiments assigned to separate channels rather than pooled).

#### Bulk transcriptomic and epigenomic data

##### RNA-seq

Bulk RNA-seq data were obtained from ENCODE^96^, GTEx^34^, and recount3^97^; apart from a small number of recount3 additions, the corpus matches AG1^14^. Processed base-resolution coverage bigWig files were downloaded from ENCODE and recount3 (including GTEx read coverage files, which were obtained via recount3). These were normalized to a genome-wide sum of 10^8^ (100 reads per million; normalized jointly across forward and reverse files for stranded libraries), and replicate experiments sharing a reference condition and channel attributes (data source, poly(A)^+^ vs. total RNA enrichment, and paired-end vs. single-end sequencing) were averaged into a single track. Alongside base-resolution coverage, gene-level targets for the bulk gene expression head (Output heads) were compiled across GENCODE annotations (v46 for human, vM23 for mouse) as transcripts per million (TPM) for each condition and channel, ignoring strand, and supervised on protein-coding and long non-coding RNA (lncRNA) genes. For ENCODE tracks, we used the TPM values from the ENCODE gene quantification files of every experiment pooled into the track, averaged across files. We excluded 41 ENCODE experiments whose TPM profiles correlated poorly (Pearson *r* < 0.1) with expression derived from their coverage tracks. For GTEx tracks, we used the GTEx v8 gene TPM from the GTEx Portal, averaged across donors within each tissue. For other data from recount3^97^, we averaged the recount3 gene TPM values across the samples of each cell type.

##### ATAC-seq and DNase-seq

Bulk accessibility datasets were obtained from the AG1 corpus^14^ and processed identically except that replicate aggregation within each reference condition separated paired-end and single-end experiments into distinct channels. As in AG1 and ChromBPNet^17^, read 5’ ends were shifted by +4/ 4 bp for ATAC-seq (Tn5 transposase offset) and 0/+1 bp for DNase-seq (DNase I offset), and bigWig coverage tracks were normalized to a genome-wide sum of 10^8^.

##### Transcription factor and histone ChIP-seq

Histone modification and transcription factor (TF) ChIP-seq fold-change-over-control tracks were obtained from ENCODE^96^ via the AG1 corpus^14^, aggregating replicates per reference condition separately for paired-end and single-end experiments.

##### CAGE

CAGE data were obtained from the AG1 corpus with an identical processing pipeline. These data were originally obtained from FANTOM5^98^. Stranded bigWigs were jointly normalized to sum to 100 million.

#### Contact map data

Contact maps were derived from ENCODE Intact Hi-C and 4D Nucleome Micro-C experiments across human and mouse (including the H1 embryonic stem cell and human foreskin fibroblast Micro-C maps used by Orca^99^; 77 maps total across 73 human and 4 mouse reference conditions), selecting a single contact map source per (assembly, reference condition) pair. Each contact map was extracted from its multi-resolution cooler file^100^ at 1000 bp resolution (before training-time interpolation to 2048 bp; see below) and balanced by Iterative Correction and Eigenvector decomposition (ICE)^101^ using cooler. In contrast to AG1 and Orca, we did not apply adaptive coarse-graining, as we found it led to overfitting. Contact map prediction accuracy is therefore not quantitatively comparable to these two models.

##### Preprocessing

For each anchor bin *i*, we stored the balanced contact frequencies with downstream bins *j* = *i*+*d* for *d* ∈ {0, … , *D* – 1}, where *D* = ⌈2^20^/1000⌉ = 1049 diagonal bins cover all pairwise distances within the 1-Mb receptive field at 1000-bp resolution. We then applied observed-over-expected distance normalization: for each contact map, a genome-wide mean profile *m_d_* was computed by averaging strictly positive (>0), non-`NaN` balanced contact frequencies at separation *d* across all chromosomes, and each contact value was normalized as *y_i_*_,*j*_ = *x_i_*_,*j*_/*m_|i−j|_*, so that *y* = 1 represents the genome-wide expectation at that separation, *y* > 1 indicates enriched interactions (such as enhancer–promoter loops), and *y* < 1 indicates depleted or insulated contacts. The normalized data were serialized as per-bin records indexed by global genomic coordinates, with metadata recording *m_d_* and per-chromosome offsets for each (assembly, reference condition) pair.

##### Training-time interpolation

At training time, the stored upper-triangular contacts were reconstructed into a symmetric matrix and interpolated from 1000 bp to the 2048-bp pairwise resolution by area-weighted mean pooling, weighting overlapping source bins by their fractional area of overlap. Unmappable (`NaN`) bins were excluded from the weighted mean, and target bins with >25% `NaN` area overlap were masked as `NaN`.

#### RNA splicing

AG2 predicts the same types of splicing targets as AG1, comprising splice junctions (which model RNA-seq read coverage across junctions), splice site usage (SSU, which quantifies the proportional usage of a splice site), and splice site classification labels.

These targets were derived from the same ENCODE and GTEx RNA-seq datasets used for bulk RNA-seq, and were grouped by ontology CURIEs consistent with the bulk RNA-seq processing pipeline. Unlike the bulk RNA-seq coverage pipeline, the splicing pipeline used only paired-end libraries. GTEx coverage tracks are unstranded, so we realigned the GTEx reads with STAR and inferred junction strand from splice-site dinucleotide motifs (`--outSAMstrandField intronMotif`). This yielded three library types (ENCODE total RNA-seq, ENCODE poly(A)^+^ RNA-seq, and GTEx poly(A)^+^ RNA-seq), each on the forward and reverse strand (Supplementary Table 6).

##### Read alignment

Paired-end RNA-seq reads (converted from name-sorted BAM files to FASTQ where necessary) were aligned to GRCh38.p13 (GENCODE v32) for human and GRCm38.p6 (GENCODE vM23) for mouse using STAR v2.7.11b^102^. Key STAR parameters included: a minimum splice junction overhang of 8 bp (`--alignSJoverhangMin 8`), up to 20 multi-mapped alignments (`--outFilterMultimapNmax 20`), at most 4% mismatches per read length (`--outFilterMismatchNoverReadLmax 0.04`), and strand inference from splice dinucleotide motifs (`--outSAMstrandField intronMotif`). The splice junction table (`SJ.out.tab`) produced by STAR served as the raw input for all subsequent splicing targets.

##### Splice site usage

For each splice junction identified by STAR, the SSU ratio was computed at both the donor and acceptor positions. Following the approach of SpliSER^103^ and AG1, a pileup-based extractor queried the BAM file at each splice site position and counted: *k*, the number of reads spliced at that site (identified via CIGAR *N* operations whose boundaries matched the site coordinate), and *n*, which included reads that spanned the site without splicing and reads that skipped the site via an intron from a competing junction. Reads flagged as PCR duplicates, those with mapping quality below 30, or base quality below 20 were excluded. SSU was computed as SSU = *k*/(*k* + *n*).

##### Per-condition aggregation

Junction counts and SSU counts from all samples belonging to the same ontology CURIE were combined into a single table. STAR junction coordinates were adjusted to 1-based exon-boundary conventions. Junctions on mitochondrial chromosomes, those with unknown splice motifs, and unstranded junctions were discarded.

##### Filtering and normalization

Per-condition junction counts were normalized to a common scale of 10^8^ mapped bases. For both GTEx and ENCODE, the normalization denominator estimated the total mapped bases (approximately total reads read length). The two sources differed only in how this quantity was obtained: from bigWig per-base coverage sums for GTEx, and from STAR read counts multiplied by mapped read length for ENCODE. Each junction was annotated against GENCODE (v46 for human, vM23 for mouse) to identify novel (unannotated) junctions. A minimum average count threshold of 1 normalized read across replicates was applied.

##### Cross-species dataset creation

Per-condition filtered junctions were merged across all tissues within each species into a unified splicing dataset. Junctions were ranked by total evidence (sum of normalized counts number of replicates, summed across all conditions), and the top 600,000 junctions were retained. Similarly, donor and acceptor splice site positions were ranked separately by evidence within each species, retaining the top 300,000 donor sites and top 300,000 acceptor sites alongside all annotated GENCODE sites (v46 for human, vM23 for mouse). When a genomic position appeared as both a donor and an acceptor on different strands, the assignment with higher total evidence was retained, and junctions using the discordant assignment were removed. The final dataset contains, for each species: the normalized junction count per condition, the SSU ratios (SSU_donor_, SSU_acceptor_), and per-junction GENCODE annotation status.

##### Splice site classification labels

Splice site classification labels were computed from the SSU ratios at training time, and were defined per condition rather than per assay. Each splice site has a position, a strand and one SSU ratio per track, where a track is a (condition, assay) pair. A condition can therefore have up to three tracks (ENCODE total RNA-seq, ENCODE poly(A)^+^ RNA-seq and GTEx poly(A)^+^ RNA-seq). As in AG1, we define the donor site as the last nucleotide before the intron and the acceptor site as the first nucleotide after it. For a sampled condition, we took all tracks of that condition and labeled a position as a forward-strand donor, forward-strand acceptor, reverse-strand donor or reverse-strand acceptor if the corresponding site had SSU > 0 (at least one read spliced at the site) in at least one of these tracks. All other positions were labeled as non-splice sites. Conditions without splicing tracks contributed no classification loss. In the imputation split (Imputation split), the classification labels of a condition were dropped entirely if any of its splicing tracks were held out. In contrast, SSU and junction targets remained specific to each assay and strand channel.

#### CpG methylation data

DNA methylation profiles were obtained from two complementary human methylome atlases spanning distinct experimental technologies: bulk whole-genome bisulfite sequencing (WGBS)^28^ and single-nucleus methyl-3C sequencing (snm3C-seq)^29^. Mapping experimental samples from both atlases to composite ontology CURIEs combining an anatomical tissue identifier from UBERON and a cell-type identifier from CL (Ontology organization) yielded 190 methylation profiles (75 bulk WGBS profiles and 115 single-nucleus pseudobulks) across 185 unique human reference conditions.

To process these datasets into a unified representation, all genomic coordinates were aligned to a global reference index of human CpG sites on GRCh38. Bulk WGBS data were obtained from GEO (accession `GSE186458`) in the CpG-centric BETA format^28^, where forward and reverse strand reads at palindromic CpG dinucleotides are inherently combined, and counts were summed across donors for each cell type. For single-nucleus methylomes, raw stranded ALLC files were obtained from Hugging Face (https://huggingface.co/datasets/zhoujt1994/HumanCellEpigenomeAtlas_sc_allc)^29^, processed by extracting CpG contexts with strand merging using ALLCools^104^, grouped into pseudobulks per tissue and cell type, and joined against the global CpG reference index. For each profile, methylated and total read counts were then scaled by a common factor so that total coverage summed to 10^8^ genome-wide (as for the coverage modalities), which preserves per-site methylation fractions. For both technologies, this yielded unstranded CpG dinucleotide profiles, generating paired bigWig tracks that recorded observed methylated read counts and total read coverage separately. In the bigWig tracks, each value spanned the full 2-bp CpG dinucleotide, assigning identical counts to both the cytosine and the paired guanine. Unmeasured CpG sites lacking coverage in a given condition were assigned sentinel values and masked during training.

#### Single-cell RNA-seq and ATAC-seq data

To construct the single-cell gene expression and chromatin accessibility tracks used during AG2 pre-training, we assembled a curated corpus of single-cell and single-nucleus RNA-seq, single-cell ATAC-seq, and paired multiome datasets spanning human and mouse tissues, developmental stages, and reference cell lines (Supplementary Tables 1 and 3).

##### Data sources and processing

Datasets were obtained from public repositories and were either uniformly reprocessed from raw reads or used as processed by their original source (Supplementary Table 3).

###### Reprocessed datasets

For a core set of public human scRNA-seq, scATAC-seq, and 10x Genomics Multiome datasets, raw reads were uniformly aligned to GRCh38.p14 (GENCODE v44) using Cell Ranger v8.0.1 or v9.0.1 (scRNA-seq, intronic reads included), Cell Ranger ATAC v2.1.0 (scATAC-seq), or Cell Ranger ARC v2.0.1 or v2.0.2 (Multiome, aggregated via `cellranger-arc aggr` without depth normalization), with ambient RNA removed by CellBender v0.3.0 (`remove-background`, false-positive rate 0.01, 150 epochs)^105^ for selected scRNA-seq datasets (Supplementary Table 3). We retained RNA cells with >200 detected genes (without additional mitochondrial, upper unique molecular identifier (UMI), or doublet filters) and ATAC cells with 2000– 100,000 fragments and transcription start site (TSS) enrichment ≥5 in SnapATAC2 v2.8.0^106^. Metadata were harmonized to CELLxGENE schema v5.2.0 and published cell annotations were transferred by exact barcode matching; no mouse datasets were reprocessed.

###### Datasets used as released

All other datasets were used as processed by their source: (i) single-cell RNA-seq, ATAC-seq and Multiome data from ENCODE, processed with the ENCODE uniform pipelines; (ii) count matrices from CZ CELLxGENE Discover, for more recent datasets already processed with current alignment pipelines; and (iii) author-processed count matrices and ATAC fragment files from study repositories (GEO, ArrayExpress, Zenodo and consortium portals).

##### Curation

We retained only healthy, unperturbed samples, removing cells from disease samples, genetic perturbation cohorts, or missing tissue, cell-type, or assay metadata. Each cell was assigned a composite ontology label combining its tissue (UBERON) and cell type (Cell Ontology, or an NTR identifier where no Cell Ontology term existed). For datasets distributed through CZ CELLxGENE Discover or ENCODE that already provided curated ontology CURIEs matching the granularity of the authors’ clusters, we retained those pre-assigned ontology terms directly; for study-repository datasets lacking ontology CURIEs, or where standardized repository CURIEs were substantially coarser than the authors’ cluster annotations, we manually mapped the author cluster labels to Cell Ontology terms (Supplementary Table 3). In addition, the two peripheral blood mononuclear cell (PBMC) datasets were re-annotated by label transfer for consistency across modalities: labels from the CITE-seq PBMC reference of Hao *et al.*^107^ were transferred to the Kock *et al.* scRNA-seq dataset^108^ using totalVI^109^, and then from the relabeled Kock *et al.* reference to the Benaglio *et al.* scATAC-seq dataset^110^ using scVI and scANVI (scvi-tools v1.5.1)^51,111,112^. For the Kock *et al.* dataset (restricted to libraries with intronic reads counted), cells were mapped onto a pre-trained Hao *et al.* totalVI reference model^113^ via query-model training (library as batch covariate; protein counts treated as missing; 100 epochs with weight decay 0 and adversarial loss disabled), classified in the totalVI latent space using a balanced-subsample random forest classifier trained on the reference level-2 annotations, and mapped to Cell Ontology terms. For the Benaglio *et al.* dataset, SnapATAC2 paired-insertion gene activity scores (excluding cells without donor IDs) and reference scRNA-seq counts were restricted to 5000 batch-aware highly variable genes in Scanpy^114^ (Seurat v3 flavor^115^, genes detected in ≥5 cells), jointly embedded with a two-layer, 20-dimensional scVI model (negative binomial likelihood with gene-batch dispersion; library and donor as batch covariates; batch size 1024; up to 30 epochs with patience-5 early stopping), and classified using scANVI with a linear classifier trained for 30 epochs (batch size 1024), excluding reference cell types with fewer than 50 cells. Finally, to prevent duplicate representation of the same reference condition across studies, we prioritized dedicated organ-specific atlases over broad multi-organ atlases (retaining multi-organ atlas cells only in tissues lacking a dedicated organ atlas) and kept a single source dataset whenever two studies still profiled the same composite ontology label (Supplementary Table 3).

##### Pseudobulking

For scRNA-seq, raw UMI counts were summed across all cells sharing the same biosample label, assay, suspension type (cell or nucleus), and processing pipeline within each dataset. Pseudobulks from fewer than 50 cells were discarded, with one exception retained for single-cell expression quantitative trait locus (eQTL) evaluation (AXL^+^ dendritic cells, 40 cells). Genes were restricted to those in the GENCODE annotation (v46 for human, vM23 for mouse), and each pseudobulk was normalized to counts per million (CPM).

For scATAC-seq (all derived from nuclear suspensions), fragment files were filtered to quality-controlled cell barcodes (and barcodes shared with scRNA-seq for Multiome datasets). Because public fragment files differ in whether reverse-strand coordinates use a +4/-4 or +4/-5 Tn5 transposase offset, we inspected each dataset and shifted +4/-5 fragment files by +1 bp with a custom script so that all insertion sites followed the +4/-4 cut-site convention^17^. Pseudobulks were then formed without fragment-length filtering by summing Tn5 insertion counts at base-pair resolution across all cells within a dataset sharing the same biosample label, assay and processing pipeline, using SnapATAC2^106^. Pseudobulks from fewer than 50 cells were discarded. Each pseudobulk track was then normalized by its genome-wide total insertion count and scaled to a total of 10^8^ (i.e. 100 reads per million).

#### Single-cell RNA-seq data processing for cell encoder training and post-training

Training the cell encoder and Stage 4 post-training (Stage 4: Post-training for expression conditioning) used individual cells rather than pseudobulks, and underwent less curation. We combined the pre-training scRNA-seq datasets with all human scRNA-seq datasets from CZ CELLxGENE Discover (snapshot date: 16 March 2026), restricted to cells profiled with UMI-based high-throughput assays (10x Genomics RNA assays, SPLiT-seq, ScaleBio, and sci-RNA-seq3). To avoid counting cells that appear in more than one CELLxGENE collection, we kept only cells marked as primary data. Unlike the pre-training corpus, we retained disease samples and applied no tissue deduplication or minimum cell counts; we removed only cells missing tissue, cell-type, or assay annotations. For autoencoder training only, we additionally included unlabeled human basal ganglia single-nucleus RNA-seq profiles (10x Genomics Multiome) from the Human and Mammalian Brain Atlas (HMBA)^116^. Raw UMI counts for each cell were used directly as input.

### Data splits

Evaluating AG2 across sequence prediction, cross-modality imputation, adaptation to unseen single-cell datasets, and variant interpretation requires holding out distinct axes of the data so that no evaluation leaks its target axis during training (Evaluated model configurations). We defined four complementary split schemes, detailed in the paragraphs below:

- **Genomic interval split:** The 8-fold contiguous 1-Mb interval cross-validation split from AG1^14^ and Borzoi^16^, using *fold-1* to evaluate sequence predictions on held-out intervals and *all-folds* when full-genome sequence coverage is required.
- **Track-level imputation split:** Full-genome (*all-folds*) training with designated validation, test, and withheld bulk PBMC and astrocyte tracks masked out, evaluated on *fold-5* intervals.
- **Temporal single-cell dataset split:** A chronological split of CZ CELLxGENE Discover withholding datasets published after 1 October 2025 (plus held-out validation collections) from cell encoder training and Stage 4 post-training of the *fold-1* models to test generalization to newly published datasets, whereas *all-folds* models used all datasets.
- **Variant benchmark split:** The chromosome-level partition from AG1^14^, separating validation chromosomes used for model selection from held-out test chromosomes used for zero-shot variant effect reporting.

#### Reference genomes

Human training targets and evaluations used the GRCh38 reference assembly and GENCODE v46 gene annotations, except for the Parkinson’s disease QTL and enhancer–gene (E2G) evaluations, which were run natively on T2T-CHM13v2.0. Mouse training targets and evaluations used GRCm38 and GENCODE vM23 gene annotations.

#### Genomic interval split

Following AG1^14^ and Borzoi^16^, each reference genome was partitioned into eight contiguous blocks defining eight cross-validation folds (*fold-0* through *fold-7*) over predefined 196,608-bp (3×2^16^ bp) genomic intervals, where each fold assigns six blocks to training, one block to validation, and one block to testing. Each 196,608-bp interval was expanded around its midpoint to a 1-Mb (2^20^ = 1,048,576 bp) model window, excluding any validation or test interval whose 1-Mb window overlapped a training window in the same fold. While training operated across full 1-Mb windows, interval-based evaluations restricted scoring to the central 196,608-bp target window (discarding 425,984 bp from each flank) to avoid sequence-edge effects (Interval evaluations). Interval prediction benchmarks and architectural ablations used models trained on *fold-1* and evaluated on its held-out test intervals, whereas models used for cross-modality imputation, genome-wide variant scoring, and full-atlas inference were trained across all eight blocks (*all-folds*).

#### Imputation split

AG2 can impute unmeasured modalities for a reference condition provided that at least one modality was observed during training. To evaluate common imputation directions (such as predicting chromatin accessibility from RNA-seq or histone ChIP-seq from RNA-seq and accessibility), we designed a track-level imputation split partitioning tracks into training, validation, test, and withheld (labeled “unused” in Supplementary Table 4) sets without holding out genomic intervals during training. Withheld tracks were excluded from training and from the bulk validation/test benchmarks, comprising conditions with all tracks withheld and bulk PBMC and astrocyte tracks withheld for downstream single-cell evaluations. We trained a dedicated *all-folds* model through pre-training, condition refinement, and distillation on the training tracks, and evaluated validation and test tracks on *fold-5* intervals (1749 human intervals).

In the bulk imputation benchmark, we held out 184 test tracks (181 after merging stranded RNA-seq pairs) across 30 human reference conditions (plus 26 validation tracks across 12 conditions; 32 conditions total) spanning bulk RNA-seq, ATAC-seq, DNase-seq, histone ChIP-seq, and TF ChIP-seq (Supplementary Fig. 7a, Supplementary Table 4; Imputation metrics) to compare AG2 against retrained Avocado^45^ and the mean and closest training-track baselines (Supplementary Fig. 7b,c, Supplementary Fig. 8). Withholding all bulk tracks from sorted human PBMC populations and astrocytes likewise enabled evaluating primary-cell H3K27ac enhancer imputation from scRNA-seq and scATAC-seq (Supplementary Fig. 7d,e), TenK10K single-cell QTL prediction without bulk PBMC data (Supplementary Fig. 5e), and E2G linking in the human astrocyte CRISPR interference (CRISPRi) screen from scRNA-seq alone (Fig. 4c).

The number of training and held out tracks across organisms and modalities is summarized in Supplementary Table 4.

#### Encoder post-training data split

The dataset partitioning used for training the single-cell expression autoencoder and Stage 4 post-training is detailed in Single-cell expression autoencoder.

#### Variant benchmarks split

To evaluate generalization to unseen genomic loci and prevent data leakage, all variant effect prediction benchmarks, with the exception of the IBDverse sc-eQTL evaluation, were conducted zero-shot using a chromosome-level partition consistent with AG1^14^. Variants were assigned to validation or test sets based on chromosome:

- **Validation set:** Variants on chromosomes 1, 2, 4, 5, 7, 8, 10, 11, 13, 14, 15, 17, 20, 22, and X were used to guide model selection and hyperparameter tuning.
- **Test set:** Variants on chromosomes 3, 6, 9, 12, 16, 18, 19, and 21 formed the held-out test set on which all final benchmark performances are reported.

For the IBDverse sc-eQTL evaluation, we reported the results on all chromosomes from which we excluded variants present in other QTL validation sets as the smaller IBDverse donor cohort yielded fewer high-confidence fine-mapped variants than GTEx or TenK10K (Single-cell eQTLs). The IBDverse sc-eQTL study was not used for model nor scorer selection.

### Model

AG2 takes as input a 1-Mb DNA sequence (2^20^ = 1,048,576 bp), species identity (human or mouse), and a target condition (tissue, cell line, cell type, or cell state), and makes predictions across diverse genomic modalities. The condition is represented as a condition embedding, obtained in one of two ways. For *reference conditions*, it is taken from a learned dictionary of reference condition embeddings indexed by ontology CURIEs. For *expression conditioning*, cell embeddings are obtained by passing scRNA-seq profiles through the cell encoder of a separately trained single-cell expression autoencoder (resembling scVI^51^; Single-cell expression autoencoder) and mapped to a condition embedding by an adapter module. Whereas reference condition embeddings are learned directly during sequence model pre-training, expression conditioning uses Stage 4 post-training to optimize the adapter and denoised single-cell gene expression head. By modulating intermediate sequence representations via condition embeddings, AG2 explicitly decouples conditions from assay modalities.

Rather than predicting an exhaustive matrix of fixed combinations of condition and modality (as in AG1), each output head predicts only the assay channels for the queried condition, where channels capture technical factors (such as assay differences within a modality) and are reused across conditions. Predictions for a given condition take the form of sequencing-read-based coverage tracks (chromatin accessibility, transcription, transcription factor binding, histone modifications, and CpG methylation), scalar gene expression values, and pairwise interactions representing contact maps and splice junctions.

AG2 builds on the five-component encoder–decoder architecture of AG1^14^ (a convolutional sequence encoder, a transformer tower, pairwise interaction blocks, a convolutional sequence decoder with U-Net skip connections, and modality-specific output heads) with the following architectural changes:

- **Conditioning.** A conditioning vector is constructed from a taxonomy embedding (human or mouse) and a condition embedding obtained either (1) from ontology CURIEs for reference conditions or (2) through expression conditioning, by passing cell embeddings from the cell encoder through the adapter (Condition embeddings). The conditioning vector is broadcast throughout the network via Feature-wise Linear Modulation (FiLM^26^; FiLM conditioning).
- **Normalization.** Batch normalization is replaced everywhere by root-mean-square (RMS) normalization in the encoder and decoder, and by layer normalization in the transformer trunk and pairwise blocks (Sequence encoder and Transformer tower), eliminating running batch statistics.
- **Convolutions.** Weight-standardized convolutions^117^ are replaced with standard 1D convolutions (Sequence encoder).
- **Transformer.** Post-normalization is removed; per-head layer normalization of queries, keys, and values is removed (Transformer tower).
- **Sequence-to-pair projection.** Learned relative positional encodings in the sequence-to-pair projection are replaced with rotary position embeddings (RoPE^118^, matching the transformer tower); the outer sum of projected sequence embeddings is replaced with an outer absolute difference; RMS post-normalization is applied after the projection (Pairwise blocks).
- **New output heads.** A methylation head predicting CpG methylation fractions via a binomial likelihood, and gene-expression heads for both bulk and single-cell RNA are added (Output heads).
- **Channel layout per head.** Each channel within an output head captures technical assay factors (such as 10x Genomics 3’ scRNA-seq with cell versus nuclei suspension) and is reused across conditions, rather than assigning separate channels to individual conditions.
- **Track normalization.** Per-assay scaling replaces per-track scaling (Output heads).
- **Training.** A four-stage training protocol is introduced: pre-training, condition refinement, distillation, and post-training for expression conditioning (Training).

The model was implemented using JAX^119^ and Haiku^120^.

#### Condition embeddings

AG2 explicitly decouples conditions from output assays by conditioning intermediate sequence representations on a *D*-dimensional conditioning vector *c* ∈ ℝ*^D^* (*D* = 256):

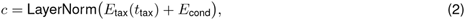

where *E*_tax_ ∈ ℝ^2^*^×D^* embeds species taxonomy (human vs. mouse), and *E*_cond_ ∈ ℝ*^D^* is the condition embedding.

The condition embedding is supplied through one of two pathways:

- **Reference condition embeddings:** Each reference condition is identified by its composite 5-tuple ontology identifier *t*_onto_ (Ontology organization), and each distinct tuple in the vocabulary is assigned its own row in a learned embedding table *E*_onto_ ℝ^*V*_onto_^ *^×D^*, where *V*_onto_ is the number of ontology tuples observed in training.
- **Expression conditioning:** To condition on individual cells across single-cell compendia (such as CELLx-GENE) or unannotated cell states, *E*_cond_ = Adapter(*z*) projects a continuous 256-dimensional cell embedding *z* ∈ ℝ^256^ produced by the cell encoder of our single-cell expression autoencoder (Single-cell expression autoencoder). The adapter is a two-layer projection multilayer perceptron (MLP) with layer normalization:

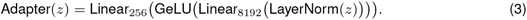

In pseudocode:

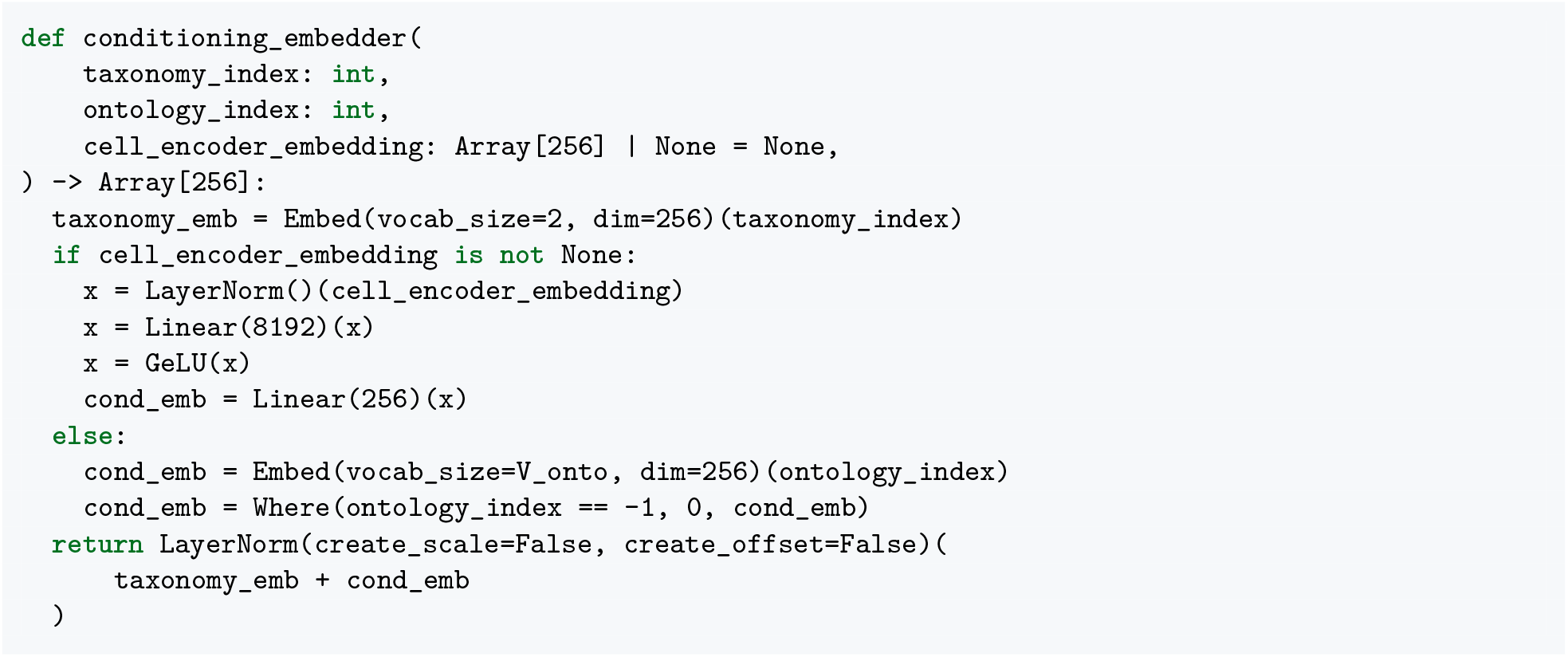

While reference condition embeddings are learned end-to-end during pre-training, the adapter is optimized on individual cell embeddings *z* from the cell encoder during Stage 4 post-training (Stage 4: Post-training for expression conditioning; Single-cell expression autoencoder). At inference, Adapter(·) can be applied either to a single cell’s embedding *z* or, when conditioning on a cell population (such as a cell type, cell state, or metacell), to the mean cell embedding averaged across the constituent cells (Fig. 3d, Supplementary Fig. 10e,f).

#### FiLM conditioning

FiLM^26^ modulates internal representations using the conditioning vector *c* ∈ ℝ*^D^*, computing per-channel scale and offset parameters applied to normalized activations *x*:

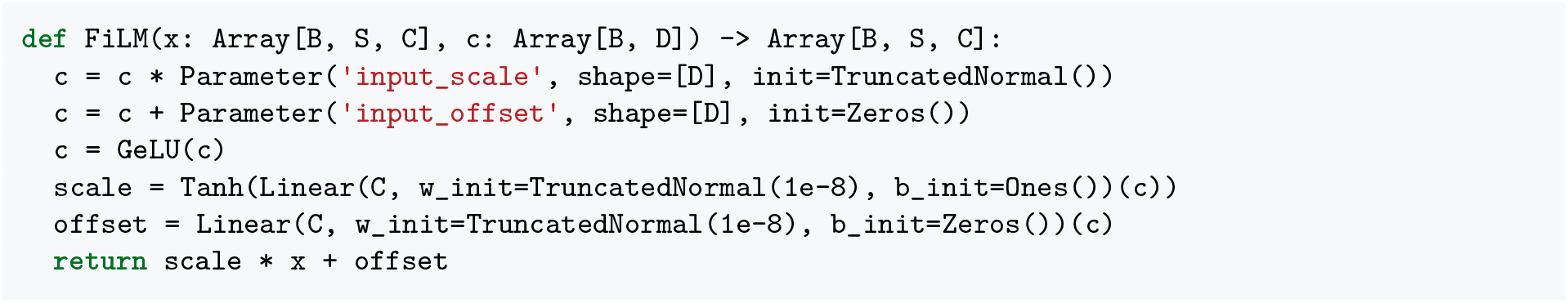

Each FiLM layer has independent parameters so modules can specialize on different features of *c*. Projections are initialized with near-zero weights (10*^−^*^8^), unit pre-activation scale bias and zero offset bias, so FiLM initially scales normalized activations uniformly by tanh(1)≈0.76 with per-channel scales bounded in (−1, 1). FiLM layers are applied immediately after normalization in every convolutional block of the encoder and decoder, and in every transformer block.

When normalization layers (RMSNorm or LayerNorm) are followed by FiLM conditioning, static affine parameters are omitted so that activations are standardized (*x*/RMS(*x*) or (*x* - *μ*)/*σ*) and all per-channel scaling and shifting are supplied by FiLM as a function of *c*. Conversely, in network modules where FiLM conditioning is not applied (such as the pairwise update blocks and output embedders), normalization layers learn static per-channel scale and offset parameters.

#### Sequence encoder

The sequence encoder maps a one-hot DNA input at 1 bp resolution (2^20^ positions) to a condition-modulated sequence embedding at 128 bp resolution (with *S* = 8192 positions and *C* = 1536 channels) through seven stages. Each stage produces a multi-resolution snapshot that is stored for use as a U-Net skip connection in the decoder.

All convolutional blocks follow the same template: RMS normalization, FiLM modulation by the conditioning vector *c*, GeLU activation, and 1D convolution. Relative to AG1, weight-standardized convolutions^117^ (which center and scale kernel weights per output channel) are replaced with ordinary unnormalized 1D convolutions, and batch normalization is replaced with RMSNorm.

The first stage embeds the DNA with a width-15 convolution into 768 channels, followed by a residual convolutional block. Each subsequent stage doubles the spatial bin size via stride-2 max-pooling and applies two residual convolutional blocks, expanding channels linearly by 128 per stage (768, 896, · · · , 1536). Channel-dimension mismatches across residual connections are handled by zero-padding.

In pseudocode:

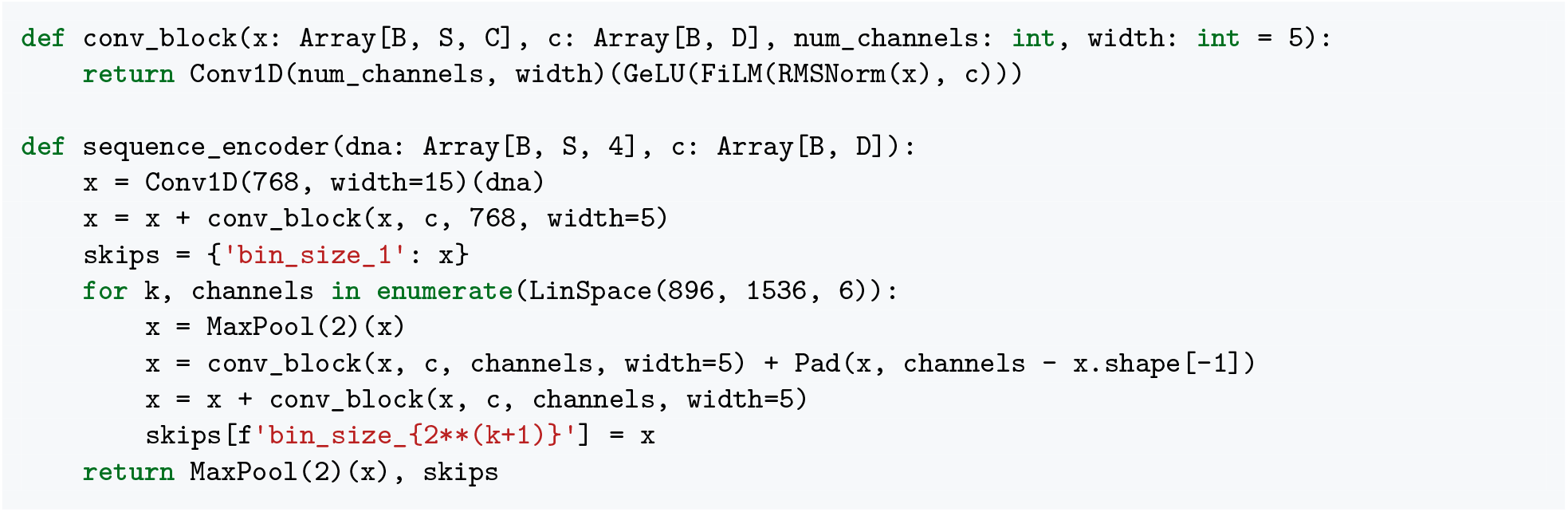

#### Transformer tower

The transformer tower updates the 128-bp sequence embeddings (*S* = 8192, *C* = 1536) through 11 stacked blocks (compared to 9 in AG1). Each block consists of multi-query attention and a feed-forward MLP, both wrapped in residual connections with FiLM-modulated layer pre-normalization. Post-normalization and per-head query/key/value layer normalizations from AG1 are removed.

Attention uses a multi-query layout: 8 query heads (128 dimensions each) share a single key head and a single value head (192 dimensions), projecting back to *C* = 8 x 192 = 1536 channels. Queries and keys are positionally encoded with RoPE. Attention logits are added to a directed pairwise attention bias (described in Pairwise blocks) and soft-clipped via 10 tanh(logits/10). The MLP expands channels 2× with ReLU activation.

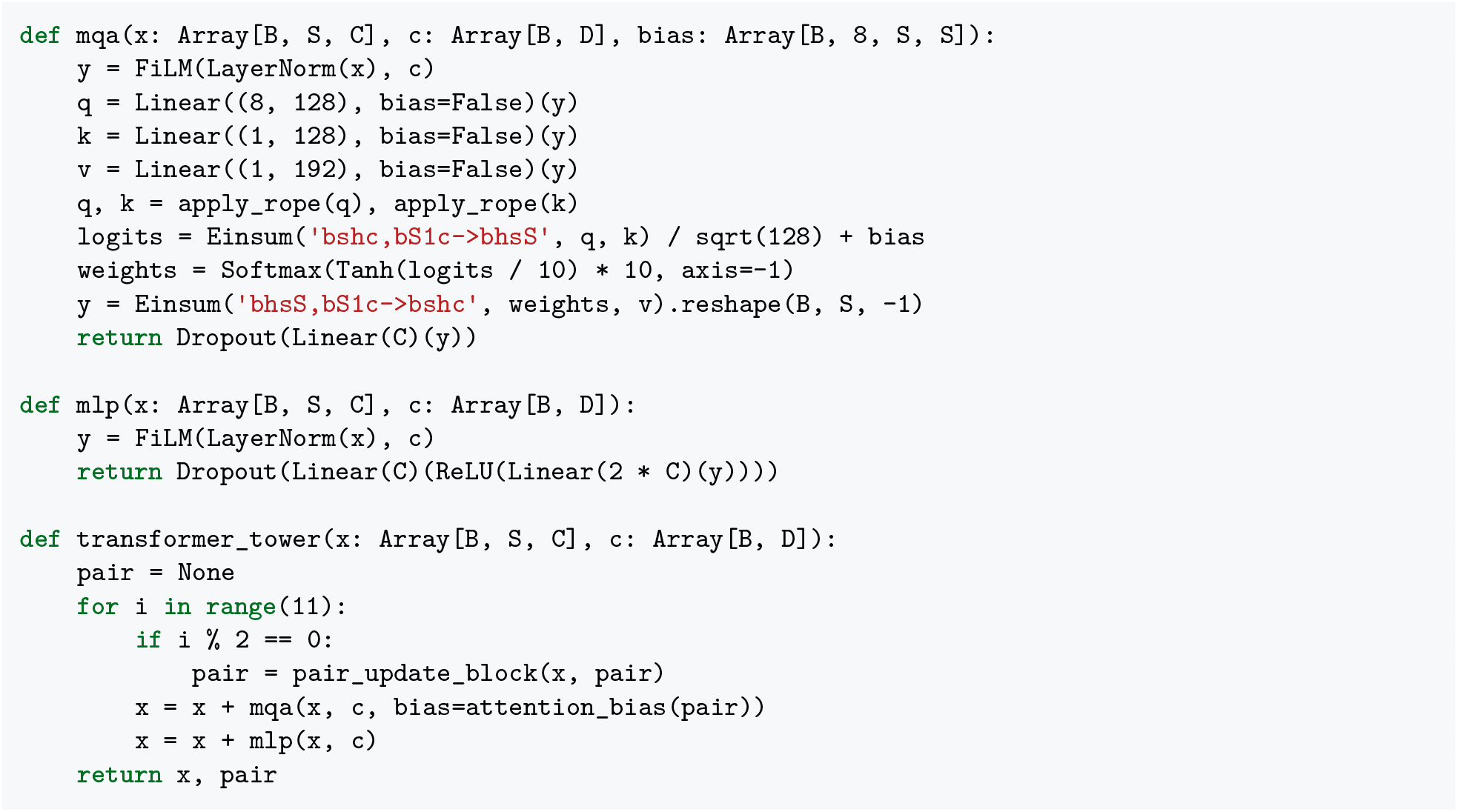

#### Pairwise blocks

A 2D pairwise representation at 2048 bp resolution (*P* = 512, *F* = 128 channels) is updated before every second transformer block and feeds an attention bias to all attention layers.

Each update begins with a sequence-to-pair projection in which 128-bp sequence representations are average-pooled 16x to 2048-bp resolution, normalized with LayerNorm, and combined across two pathways. First, queries and keys across 32 heads (128 dimensions each) are rotated by RoPE, multiplied as *q^⊤^k_j_*, and linearly projected to 128 pair channels. Second, the element-wise absolute difference |*x_i_* - *x_j_*| is linearly projected to 128 channels and added, replacing the outer sum used in AG1 with a symmetric term that empirically stabilizes pair activation magnitudes. The resulting activations are post-normalized with RMSNorm, passed through dropout, and added to the existing pair stack, which is then updated through residual single-head axial row attention and a residual MLP, both using RMS pre-normalization.

To bias transformer attention, the pairwise representation is normalized with LayerNorm, activated with GeLU, linearly projected to 8 channels (one per query head), and repeated 16 along both sequence axes to 128-bp resolution without symmetrization, preserving directional query–key context.

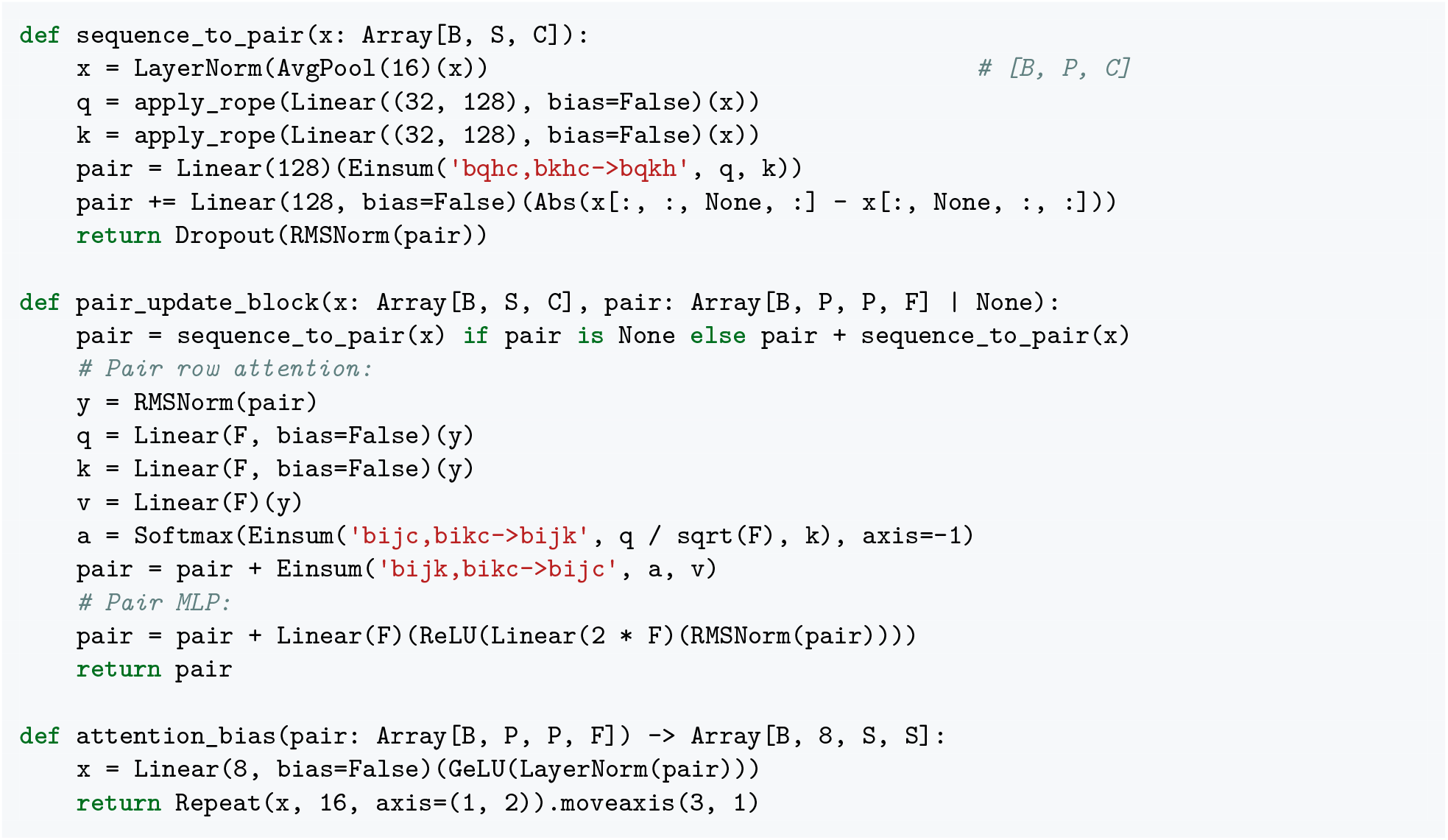

#### Sequence decoder

The sequence decoder mirrors the encoder, progressively upsampling representations from 128 bp resolution (1536 channels) back to 1 bp resolution (768 channels) through seven stages with U-Net skip connections from the corresponding encoder resolutions.

In pseudocode:

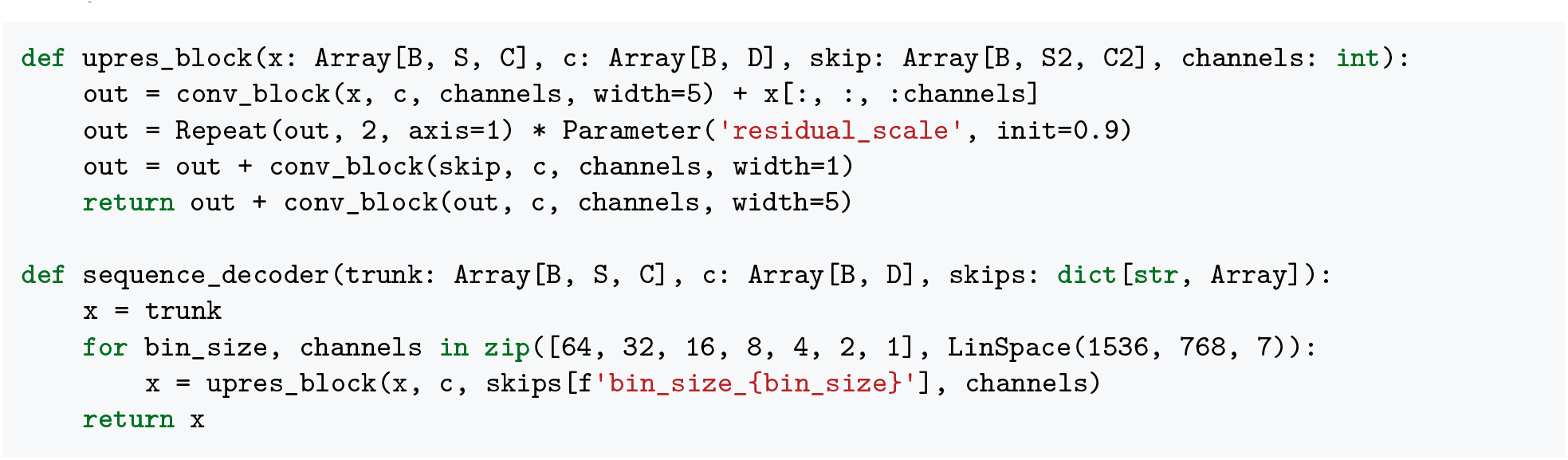

#### Output embedder

A shared pointwise embedder transforms intermediate representations into the final features consumed by the output heads. The block applies RMS pre-normalization, projects channels 2 linearly, optionally incorporates a cross-resolution skip connection from the coarser resolution, and applies a Gated Linear Unit (GLU) activation, which halves the channels back to the original dimension:

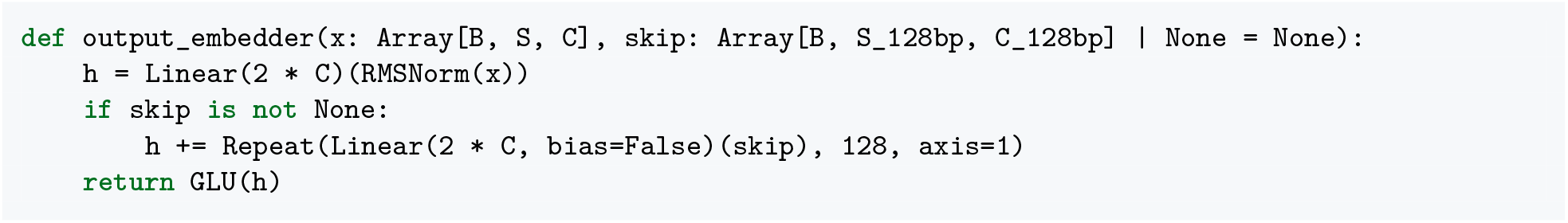

The embedder is applied coarse-to-fine: first to the 128-bp transformer trunk (*C* = 1536) without a skip connection, then to the 1-bp decoder output (*C* = 768) with a skip connection from the 128-bp embedded trunk.

For the 2D pairwise representation at 2048 bp resolution, the pair activations are finalized by symmetrizing across spatial axes, applying RMS normalization, and activating with GeLU before passing to the contact map head:

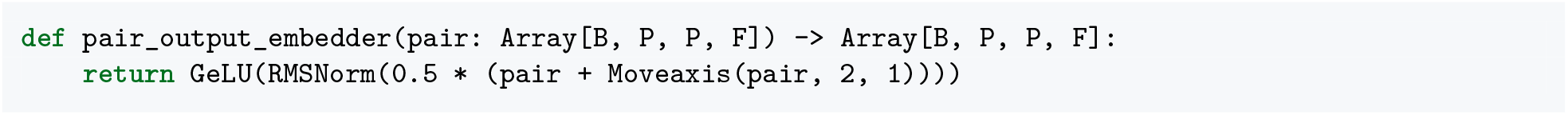

#### Output heads

In AG1^14^, output heads predicted a fixed vector of all consolidated condition–assay tracks simultaneously. In AG2, conditioning decouples the condition from the output head layout: each of the 14 output heads in Stages 1–3 (plus the Stage 4 denoised single-cell gene expression head; 15 heads total in Supplementary Tables 5 and 6) predicts a fixed set of channels reused across all conditions. Within each head, channels correspond to the technical and target grouping factors described in Ontology organization (with replicate experiments matching the same *t*_onto_ and channel pooled into a single track):

- **Bulk coverage, gene expression, and splicing heads** (RNA-seq: 12 channels; bulk gene expression: 12; CAGE: 4; DNase-seq: 2; ATAC-seq: 2; splice site classification: 5; SSU: 6; splice junctions: 6): channels index combinations of data source (ENCODE vs. GTEx), RNA enrichment (poly(A)^+^ vs. total RNA), sequencing endedness (paired-end vs. single-end), and strand orientation. Whereas AG1 pooled paired-end and single-end experiments for a given biosample into a single accessibility track, AG2 assigns paired-end and single-end experiments to separate channels in ATAC-seq (2 channels), DNase-seq (2 channels), and RNA-seq (12 channels).
- **Single-cell heads** (single-cell gene expression: 14 channels; scATAC-seq: 5 channels): channels index combinations of single-cell assay chemistry and processing pipeline, suspension type (whole-cell vs. nuclear), and strand orientation.
- **ChIP-seq heads** (histone ChIP-seq: 43 channels; TF ChIP-seq: 911 channels): channels correspond to the assayed histone modification or transcription factor split by sequencing endedness (paired-end vs. single-end).
- **Methylation and contact map heads** (CpG methylation: 2 channels at 1-bp resolution corresponding to bulk WGBS^28^ and single-nucleus snm3C-seq^29^; contact maps: 1 channel at 2048-bp resolution).

For any given reference condition *t*_onto_, experiments across different protocols or targets populate their respective channels within each head, while unmeasured channels are masked during loss calculation (Stage 1: Pre-training). The distribution of conditions is not uniform across channels: GTEx profiles post-mortem tissues, WGBS and snm3C-seq cover distinct cell-type repertoires, and specific assay protocols are enriched for particular cell lines or tissues. Condition identity is therefore partially confounded with output channel assignment. A complete summary of all output heads, channel counts, spatial resolutions, loss formulations, and per-assay scaling constants (*s*) is provided in Supplementary Tables 5 and 6, and pre-training and distillation loss weights are detailed in Supplementary Table 7. Linear projections in the output heads use weight dropout^121^: during training, an independent dropout mask is sampled once per forward pass and per accelerator chip and applied multiplicatively to the weight matrix before the matrix–vector product. Weight dropout rates are 0.1 for coverage, methylation, and gene expression projections (query, key, and output projections), and 0.2 for contact map, splice site classification, and SSU heads (for splice junctions, standard dropout at a rate of 0.2 is applied to the modulated site embeddings prior to positional encoding).

##### Coverage heads

Coverage tracks (RNA-seq, CAGE, ATAC-seq, DNase-seq, scATAC-seq, and ChIP-seq) are predicted by a linear projection followed by an exponential activation: *ŷ* = exp(Linear(*x*)). For RNA-seq, CAGE, ATAC-seq, DNase-seq, and scATAC-seq, primary predictions are produced at 1-bp resolution alongside an auxiliary prediction and loss at 128-bp resolution from the embedded trunk; histone and TF ChIP-seq heads operate at 128-bp resolution only. The loss is a combination of a Poisson loss on aggregate counts (weight *w*_tot_) and a multinomial negative log-likelihood on positional distributions (weight *w*_pos_), computed over non-overlapping 2^17^-bp windows.

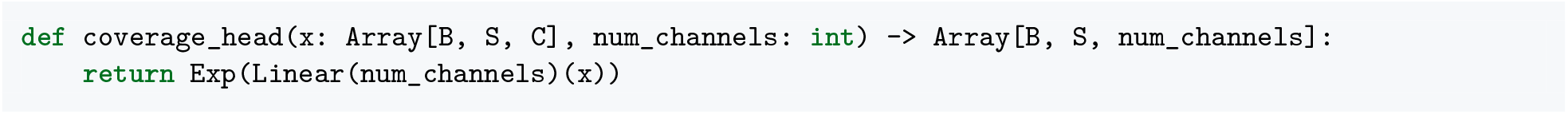

Raw targets *y* are divided by the bin width in base pairs (1 or 128) and by a per-assay scaling constant *s*, raised to the power 0.75 (RNA-seq only), and then soft-clipped via 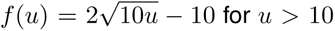 for *u* > 10, giving the scaled targets *ỹ* used in the loss (with the inverse transformation applied to *ŷ* at inference). In AG1, each output track had its own normalization constant (the mean over non-zero values for that track). In AG2, a single scaling constant *s* is shared across all conditions for each assay type. The scaling constants were computed on an early version of the data as the ratio of the total sum to the number of non-zero entries (pooled across all conditions) and rounded to fixed values.

##### Contact map head

Contact map predictions are derived from the finalized 2048-bp pairwise embeddings (Output embedder). A linear layer followed by a softplus activation maps the 128 pair channels to output contact channels, and the head is trained with a Poisson loss against observed-over-expected targets.

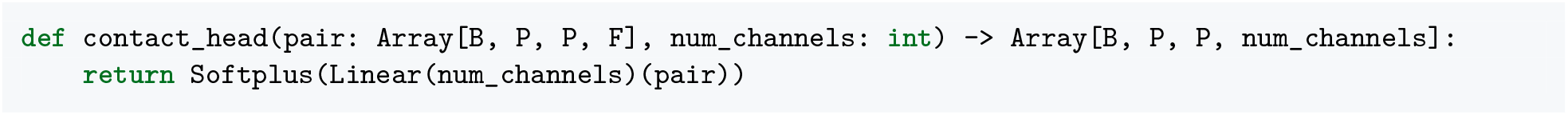

##### Splicing heads

The splicing module predicts three related targets from 1-bp sequence embeddings. First, splice site classification categorizes each position into five classes (non-splice site, forward strand donor, forward strand acceptor, reverse strand donor, and reverse strand acceptor) trained with a categorical cross-entropy loss. Splice sites are defined per condition rather than per assay, as the union over the condition’s RNA-seq assays (Splice site classification labels). Second, SSU predicts usage probabilities per assay and strand for the queried condition via a sigmoid linear projection trained with a binary cross-entropy loss. Third, splice junction counts are predicted from the pairwise interaction between donor and acceptor embeddings. A linear layer projects the trunk into a 256-dimensional splice-site embedding. At each splice site position, learned affine parameters modulate the embedding for each of the three RNA-seq assays across strand and role (donor/acceptor), followed by dropout (rate 0.2) during training and RoPE. An inner product between donor and acceptor embeddings passed through a softplus activation produces predicted scaled junction counts. Target junction counts are divided by *s* (*s* = 2), raised to the power 0.75, and soft-clipped above 10 as for coverage targets (with inverse unscaling applied at inference). The junction loss combines a Poisson loss on total counts per donor and acceptor (weight 0.1) and a cross-entropy loss on donor-to-acceptor and acceptor-to-donor read ratios (weight 0.5). Splice site positions are provided per condition during training (up to *N* = 512 sites per strand and role) from the condition’s dataset, and extracted at inference by thresholding the classification predictions (classification probability ≥ 0.1).

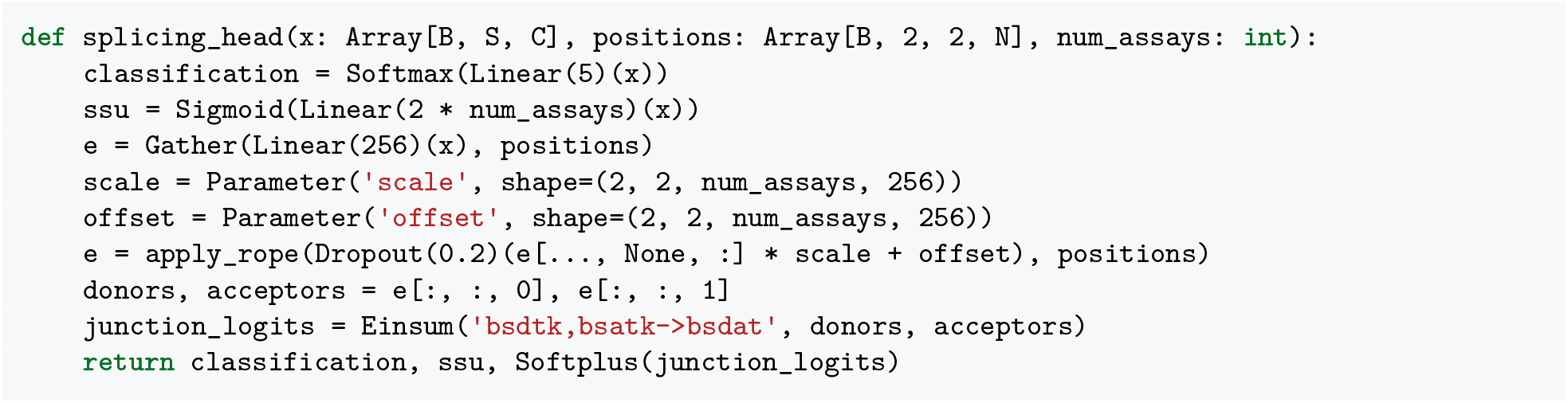

##### Methylation head

The methylation head predicts CpG methylation fractions from 1-bp sequence embeddings via a linear projection and sigmoid activation, producing predicted methylation probabilities *p* ∈[0, 1]. Given depth-normalized methylated and total coverage values *k* and *n* (CpG methylation data), the head is trained by minimizing the coverage-weighted binomial negative log-likelihood −*k* log *p−*(*n* − *k*) log(1 − *p*), masking CpG sites without coverage; at non-CpG positions *n* is set to 1 and *k* to 0, so these bases are trained to predict zero methylation. An auxiliary loss of the same form is computed at 128 bp resolution from the 128-bp embedded trunk against mean-pooled *k* and *n*.

##### Gene expression head

AG2 includes dedicated gene expression heads for both bulk and single-cell RNA-seq. Rather than predicting positional read coverage along the chromosome, these heads predict gene-level aggregated expression values for each annotated gene across the 1-Mb locus. The head takes 1-bp sequence embeddings and a binary exon mask *M*^exon^ {0, 1} *^B×S×^*^2^*^×G^* indicating annotated exonic positions on forward and reverse strands for up to *G* = 256 genes in the window. Because each gene lies on a single strand, *M*^exon^ is collapsed across the strand axis into an unstranded mask `gene_mask` ∈{0, 1} *^B×S×G^*, which the head uses with cross-attention to pool sequence features across exons:

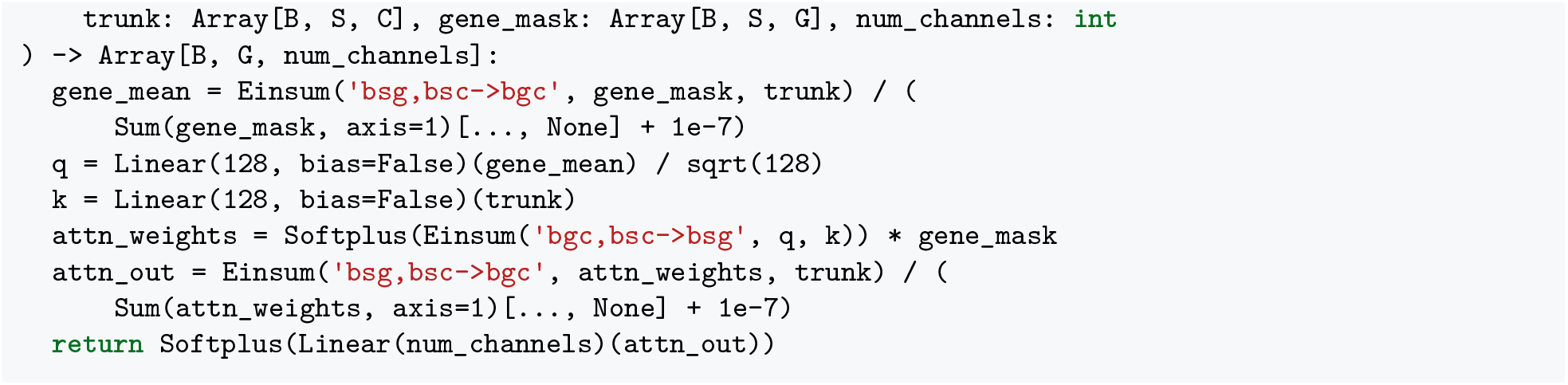

Queries *q* are derived by average-pooling the trunk embedding across each gene’s exonic positions and projecting to dimension 128, while keys *k* are projected from the full sequence. Attention weights aggregate sequence features across exons, followed by a linear projection and softplus activation to predict non-negative expression values.

Because annotated genes are transcribed from a specific strand, predictions and training targets are filtered by strand compatibility. Taking the maximum of *M*^exon^ over sequence positions gives a gene strand-presence mask *M*^gene^ ∈{0, 1} *^B×^*^2^*^×G^*, which identifies whether each gene possesses exonic sequence on the forward or reverse strand within the 1-Mb locus. For a head with *J* output channels, a fixed binary matrix *M*^chan^ ∈{0, 1} ^2^*^×J^* defines the strand assignment of each output channel *j* ∈{1, … , *J*}. The composite gene-channel presence mask is computed as:

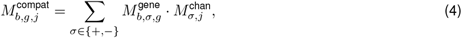

ensuring that gene *g* is active for channel *j* only if the gene is encoded on the compatible strand and has annotated exons in the receptive field. Predictions are masked by 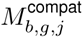. During training, target expression values *y* are transformed into scaled targets *ỹ* prior to evaluating the Poisson negative log-likelihood: bulk (*s* = 42) and Stages 1– 3 single-cell (*s* = 48) targets are divided by *s*, raised to the power 0.75, and soft-clipped above 10, whereas Stage 4 denoised single-cell targets (*s* = 48) are scaled as *ỹ* = log(*y*/*s* + 1); at inference, the corresponding inverse transformation maps *ŷ* back to the original scale. The loss is restricted to observed entries via a combined binary mask:

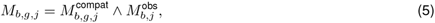

where 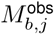 denotes whether assay channel *j* was experimentally profiled for the sampled condition. Targets for unmeasured channels or genes on the opposite strand are masked out, contributing zero loss and zero gradient. The loss is normalized by total tensor capacity:

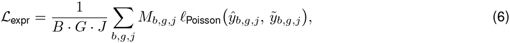

dividing the sum of masked Poisson losses by the fixed capacity *B* · *G* · *J* rather than the variable count of observed entries, thereby stabilizing gradient magnitudes across batches with differing gene densities and channel observation sparsity. An auxiliary gene expression loss is identically computed at 128 bp resolution from the 128-bp embedded trunk, with the exonic mask max-pooled to 128-bp bins.

#### Imputation

Because every output head predicts all of its channels for whichever condition is queried, AG2 can impute unmeasured modalities at inference by reading from the target output channel under either a reference condition or expression conditioning. In this work, we evaluated cross-modality imputation across bulk chromatin accessibility, RNA-seq, and histone and TF ChIP-seq, as well as primary-cell H3K27ac enhancer imputation from scRNA-seq and scATAC-seq and chromatin accessibility imputation under expression conditioning (Imputation split; Supplementary Table 4).

#### Ablations

To control for differences in bulk training data between AG1 and AG2, we trained the unconditioned AG1 multitask architecture from scratch on *fold-1* using the AG2 bulk training datasets (bulk RNA-seq, CAGE, DNase-seq, ATAC-seq, histone and TF ChIP-seq, CpG methylation, and splicing targets, alongside AG1 4D Nucleome contact maps, with PRO-cap, single-cell modalities, and the bulk gene expression head omitted). Training ran on 256 TPU v4 chips (4 × 8 × 8 topology, batch size 64) following the AG1 optimization schedule for 15,000 steps (5000 linear warmup steps to a peak learning rate of 0.004 followed by 10,000 cosine decay steps to 0), with weight decay 0.4 and dropout 0.3.

To evaluate the contribution of conditioning at different network stages (Supplementary Fig. 4), we trained five restricted FiLM configurations from scratch on *fold-1* using the Stage 1 pre-training recipe (100,000 steps, batch size 512, 3 replicate seeds per configuration) without Stage 2 condition refinement or distillation, and compared them against the unconditioned AG1 multitask model (“Multitask”, 3 seeds) and standard pre-trained AG2 (“Full conditioning”, 3 seeds). In standard AG2 (“Full conditioning”), FiLM layers are active in the sequence encoder, transformer tower, and sequence decoder, while the pointwise output embedder uses static normalization without FiLM. The five restricted configurations enable FiLM selectively in: the pointwise output embedder only (“Output only”), the sequence decoder only (“Decoder only”), both the decoder and output embedder (“Decoder & output”), the transformer tower and decoder (“Transformer & decoder”), or the transformer tower, decoder, and output embedder (“Transformer, decoder & output”). In all modules where FiLM is disabled, normalization layers fall back to static per-channel scale and offset parameters. Because the histone and TF ChIP-seq heads read 128-bp representations directly from the transformer trunk via the output embedder without passing through the sequence decoder, “Decoder only” provides no condition-specific signal along the ChIP-seq pathway.

### Training

AG2 was trained in four sequential stages: pre-training, condition refinement, distillation, and post-training for expression conditioning.

#### Stage 1: Pre-training

During pre-training, AG2 was trained across human and mouse reference genomes to jointly predict all output modalities for the set of reference conditions. At each step, a species and a 1-Mb training interval were sampled uniformly, and a reference condition for that species was sampled with probability proportional to the number of coverage, methylation, and gene expression output heads with data available for it (assigning zero weight to the bulk gene expression head so that bulk RNA-seq was not double-counted alongside its coverage head). Sequence augmentation consisted of a random genomic shift (sampled uniformly in [ 1024, +1024] bp) and reverse complementation with 50% probability. Under reverse-complement augmentation, both the input DNA sequence and spatial target tracks were reverse-complemented, and stranded channels were swapped to their complementary strand.

##### Handling channel sparsity

Because experimental profiling is sparse across the corpus, most conditions have data for only a small subset of the model’s total output channels. To handle this sparsity without biasing gradients:

- For each training example, a binary observation mask *M* was constructed alongside the targets, indicating which channels and spatial positions contained valid experimental observations for the sampled condition (unmeasured assays and unsequenced assembly gaps were marked with 0).
- Loss functions across all heads were evaluated exclusively over observed entries by masking both predictions and targets with *M*.
- Coverage, contact map, methylation, and site-level splicing losses were normalized per example by the count of observed entries, whereas gene expression and splice junction losses were normalized by fixed tensor capacity (*B* · *G* · *J* or maximum junction slots). In all cases, unmeasured channels contributed zero loss and zero gradient.

##### Optimization

Training used AdamW^122^ with *β*_1_ = 0.9, *β*_2_ = 0.99, *ε* = 5 × 10*^−^*^7^, and a weight decay of 0.05 applied to kernel weights, scale parameters, and embeddings (biases and offsets were excluded). We used a numerically stable Adam parameterization in which the second-moment state stores 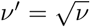 directly and updates 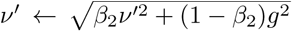 via stable hypotenuse evaluation, avoiding *g*^2^ underflow and overflow in bfloat16 arithmetic. After each update, reference condition embedding vectors whose root mean square exceeded 1.0 were rescaled to a root mean square of 1.0. The learning rate used linear warmup to 0.002 over 1000 steps and then decayed linearly to 0 over the remaining steps. Transformer and sequence-to-pair dropout rates were 0.1, and gradients were clipped to a global norm of 0.2. Training ran for 100,000 steps (125,000 for *all-folds* models) with a global batch size of 512 in bfloat16 mixed precision on 2048 TPU v4 chips (8 × 16 × 16 topology) with 4-way sequence parallelism.

#### Stage 2: Condition refinement

After pre-training, a condition-refinement stage fine-tuned only the conditioning pathway, comprising the FiLM layers, reference condition embedding table, and taxonomy embedding table, while keeping all sequence encoder, transformer tower, pairwise block, sequence decoder, and output head parameters frozen. Dropout was disabled and weight decay was set to zero. The learning rate warmed up linearly to 4 × 10*^−^*^4^ over 1000 steps and remained constant at 4 × 10*^−^*^4^ thereafter. Training ran for 50,000 steps (62,500 for *all-folds* models) with a global batch size of 1024 on 2048 TPU v4 chips using 2-way sequence parallelism, with the same data pipeline, condition sampling strategy, and loss functions as pre-training. This stage allowed the condition embeddings and modulation layers to adapt and specialize without altering the learned sequence representations.

#### Stage 3: Distillation

We previously demonstrated with AG1^14^ that distilling predictions from an ensemble of teacher models into a single student network substantially improves variant effect prediction accuracy and robustness across molecular QTL benchmarks, while eliminating the latency and compute overhead of evaluating large ensembles at inference time. Building on these results, AG2 employed a dedicated distillation stage to train a unified student model on predictions from the eight teacher models (Stages 1–2).

##### Target materialization

Distillation targets were pre-computed offline across human and mouse genomic intervals. For each training example, the input DNA sequence was perturbed with a 4% random nucleotide substitution rate, and the condition was specified via its reference condition embeddings. Each teacher model was evaluated on both the original forward strand and its reverse complement. Predictions were averaged across strands and across all ensemble members to yield strand-symmetric, denoised pseudo-targets.

To minimize storage overhead while preserving high numerical fidelity, predictions were quantized to 8-bit integers (uint8):

- **Log-space quantization** (1-bp and 128-bp coverage tracks, gene expression, splice junction counts, and contact maps): Predictions were log-transformed and linearly scaled to [0, 255] per track. When a track contained exact zeros, value 0 was reserved as a dedicated sentinel for zero while positive values were mapped to [1, 255] (255 levels); otherwise, the full [0, 255] range (256 levels) was used.
- **Linear quantization** (1-bp and 128-bp CpG methylation probabilities, SSU probabilities, and 5-class splice site classification logits): Bounded probabilities in [0, 1] and per-track [min, max] logit ranges were linearly mapped to [0, 255].
- **Exact storage** (top-512 predicted splice site positions, sequence tokens, condition indices): Stored directly without quantization.

Per-track min and max scaling statistics were stored alongside each record to enable exact dequantization at load time. Records were serialized in sharded archives, with gene masks stored sparsely as coordinate index pairs.

##### Distillation training

The student model was initialized from scratch with the same architecture as the teacher models and trained on ensembled teacher targets using the same loss formulations with rebalanced sub-weights (Supplementary Table 7), except that CpG methylation used unweighted binary cross-entropy against continuous teacher methylation probabilities in [0, 1] (with auxiliary weight 1/128 at 128-bp resolution) in place of the coverage-weighted binomial negative log-likelihood.

To increase prediction accuracy at shorter sequence lengths, we performed simulated segment-packing data augmentation during distillation. With 95% probability, the model was trained to process standard 1-Mb sequences; with 5% probability, shorter segments (8 kb to 512 kb in dyadic powers of two) were greedily packed into the 1-Mb buffer separated by 1024-bp zero-padded isolation gaps. The 1024-bp gap exceeds the 513-bp overhang radius of the convolutional encoder and gap positions were explicitly zeroed out across all encoder and decoder stages, while block-diagonal attention masking at the 128-bp transformer bottleneck ensured tokens attended only within their respective segment. Relative positional encodings ensured intra-segment representations remained invariant to absolute buffer positions, and losses were normalized strictly over valid non-gap tokens.

Distillation used the same learning rate schedule shape as pre-training (1000 linear warmup steps to 0.002 followed by linear decay to 0 over 100,000 total steps, or 125,000 for *all-folds* models) with a global batch size of 512 on 2048 TPU v4 chips, with transformer and sequence-to-pair dropout increased to 0.2, weight decay set to 0.06, and AdamW *ε* reduced to 2 × 10*^−^*^7^.

#### Stage 4: Post-training for expression conditioning

AG2 can make predictions across the entire CZ CELLxGENE corpus and generalize to unseen cell states by incorporating cell-level representations from a single-cell expression autoencoder (Single-cell expression autoencoder). Whereas Stages 1–3 conditioned on discrete reference conditions (pseudobulks defined by ontology terms for single-cell data), Stage 4 enabled conditioning on individual cells through a targeted post-training phase that adapted the distilled student model to embeddings from the cell encoder.

Starting from the Stage 3 distilled student model, the sequence encoder, transformer tower, pairwise blocks, sequence decoder, output embedders, FiLM modulation layers, reference condition and taxonomy embeddings, and all standard output heads remained frozen. Consequently, at inference under expression conditioning, predictions for chromatin accessibility and all other non-expression modalities are produced directly by the frozen standard output heads of the distilled model, modulated via FiLM by the adapter-projected cell embedding. Training optimized only two newly initialized components:

1. **Adapter:** A two-layer projection network (LayerNorm → Linear(8192) → GeLU → Linear(256)) that maps cell embeddings from the cell encoder (*z* ∈ ℝ^256^) into the 256-dimensional condition embedding space.
2. **Denoised single-cell gene expression head:** A freshly initialized gene expression head with the same architecture as the other gene expression heads (Output heads) that predicts strand-specific expression (2 channels: forward and reverse strand) for protein-coding and lncRNA genes, pooling sequence features over annotated exons.

Training was conducted at single-cell resolution over human 1-Mb training intervals from the corresponding genome split (*fold-1* for interval and dataset-adaptation evaluations, or *all-folds* for variant and cohort evaluations), with the same [ 1024, +1024]-bp shift and reverse-complement augmentations as Stage 1. Intervals were paired with human scRNA-seq profiles from the combined post-training corpus (Single-cell RNA-seq data processing for cell encoder training and post-training; 584 datasets and 20,426 (composite ontology CURIE, dataset) groups for *all-folds*). In contrast to the pooled pseudobulk profiles used in earlier stages, each training example paired a sampled 1-Mb genomic interval with a single cell embedding pre-computed across the compendium (Single-cell expression autoencoder). To ensure balanced coverage across rare cell states and study sizes, cells were drawn by two-stage hierarchical uniform sampling: first selecting a (composite ontology CURIE, dataset) group uniformly at random across all available groups, and then sampling a cell uniformly at random within the chosen group. Because each step drew one cell per sequence, a training run sampled 25.6 million cells (32 million for *all-folds* models) with replacement. This was fewer than the number of cells in the compendium, so coverage was determined by stratified sampling rather than exhaustive passes over all cells.

Rather than supervising on sparse single-cell UMI counts, Stage 4 used denoised expression profiles reconstructed by the autoencoder (Single-cell expression autoencoder), normalized per cell across the global gene vocabulary to CPM.

The denoised single-cell gene expression head was trained to predict these continuous, stranded CPM targets for annotated protein-coding and lncRNA genes (2 channels: forward and reverse strand) using a Poisson negative log-likelihood loss on targets scaled as log(*y*/48 + 1), together with an auxiliary loss of the same form at 128 bp resolution, as for the other gene expression heads (Output heads). These denoised targets provided dense supervision that improved the signal-to-noise ratio relative to sparse single-cell counts while preserving cell-state-specific transcriptional regulation.

Optimization used AdamW with a global batch size of 512 (one cell per sequence), a peak learning rate of 0.001 (1000 linear warmup steps followed by linear decay to 0 over the remaining steps), weight decay 0.02, *ε* = 5×10*^−^*^7^, and dropout disabled. Training ran for 50,000 steps (62,500 for *all-folds* models) on 2048 TPU v4 chips.

### Model selection

The training recipe (architecture, training data, and hyperparameters) was chosen by comparing runs on a composite validation score computed periodically during training. This score combined (i) track reconstruction metrics on intervals from the validation fold, each expressed as a relative improvement over a fixed reference model with respect to a chance-level baseline, with (ii) reduced-size versions of downstream benchmarks (variant effect prediction, single-cell reconstruction and imputation), weighted 35%, 35%, 20% and 10% for reconstruction, variant effects, single-cell reconstruction and imputation, respectively. The reduced variant benchmarks used fixed variant subsets drawn from validation chromosomes (Variant benchmarks split).

This score was only used to choose the training recipe. We did not use it, or any other validation metric, for early stopping, checkpoint selection or seed selection. Once the recipe was fixed, the models reported in this work were trained from scratch for a fixed number of steps (Training), and the checkpoint at the last training step was used. Test-set performance was only examined after the final models had been chosen. Because validation metrics informed only the choice of recipe, and not the selection of any individual set of model weights, validation and test set metrics are comparable.

### Single-cell expression autoencoder

To condition AG2 on single-cell transcriptomic profiles, we trained a single-cell expression autoencoder based on scVI^51^. The cell encoder maps a cell’s UMI counts across *G* genes to a cell embedding *z* ∈ ℝ^256^, and the decoder maps *z* to gene expression fractions *ρ* Δ*^G−^*^1^ on the probability simplex, which are scaled by the cell’s library size to parameterize a negative binomial (NB) likelihood over the observed counts.

#### Gene vocabulary

Expression is represented over a fixed vocabulary of the protein-coding and lncRNA genes in GENCODE v46 (*G* = 39,323). Because count matrices from CZ CELLxGENE Discover and other public repositories are processed independently by study authors (for example, by applying dataset-specific low-expression filters or aligning to different reference annotations), individual datasets may omit a subset of these genes. Any gene absent from a dataset is set to zero and treated as an observed zero count by the cell encoder and the reconstruction loss.

#### Architecture and loss

Given a cell’s UMI counts 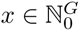 with library size 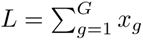, counts are normalized as 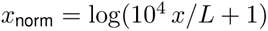, with dropout at rate 0.1 applied during training. The cell encoder projects *x*_norm_ to 2048 channels, applies two residual MLP blocks (LayerNorm → Linear(4096) → GeLU → Linear(2048)) and a final layer normalization, and projects to the cell embedding *z* R^256^. Unlike in a variational autoencoder, *z* is deterministic rather than sampled, and no Kullback–Leibler divergence term is included in the loss.

The decoder applies a single non-residual MLP block of the same form to *z* followed by layer normalization to produce *h*_dec_ ∈ ℝ^2048^, which a linear layer *W* maps to *G* logits. Without gene masking, a softmax over all *G* genes gives expression fractions *ρ* = Softmax(*W* (*h*_dec_)), parameterizing expected counts *μ* = *ρL*. Counts are modeled as *x_g_* ∼ NB(*μ_g_*, *θ_g_*), where 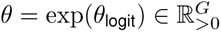 is a learned per-gene inverse dispersion shared across cells, and the unmasked reconstruction loss is the NB negative log-likelihood 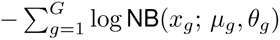. In pseudocode:

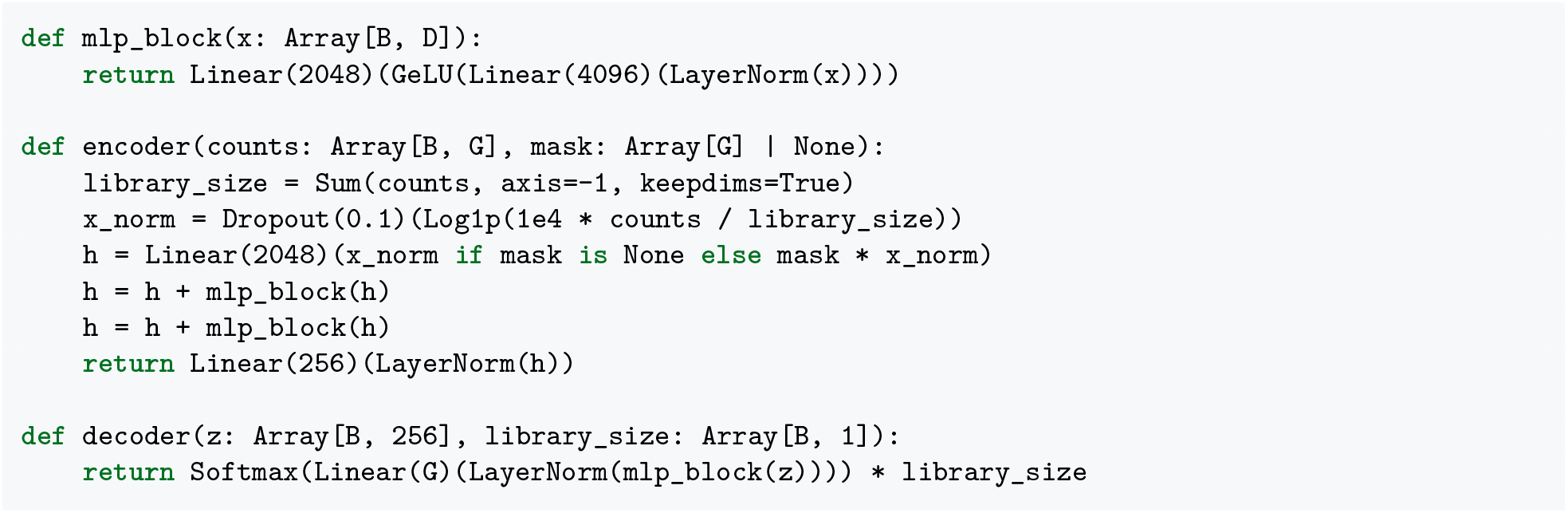

#### Gene masking

AG2 is evaluated on genomic intervals held out from training (Data splits). If the cell encoder received the expression of genes in these intervals, or if the autoencoder were trained to reconstruct them, information about held-out intervals could leak into the cell embedding. For the *fold-1* models, we therefore trained a separate autoencoder with a binary mask *m* ∈ {0, 1} *^G^*, where *m_g_* = 1 if the gene body of gene *g* is fully contained in the 1-Mb input window of at least one *fold-1* training interval and *m_g_* = 0 otherwise. The library size *L* was computed over all *G* genes, after which held-out genes were set to zero in *x*_norm_ both during autoencoder training and when computing cell embeddings for post-training and evaluation. During training, only unmasked genes contributed gradients to the cell encoder and decoder MLP: the logits *W* (*h*_dec_) of held-out genes were set to −∞ before the softmax, the resulting fractions were scaled by the unmasked library size 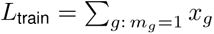 to give *μ*^train^, and the NB loss was summed over unmasked genes. A second NB loss over all *G* genes, with mean *μ*^all^ = Softmax(*W* (stop_gradient(*h*_dec_))) *L*, fit the output layer *W* and inverse dispersions *θ* (including those of held-out genes) without propagating gradients into *h*_dec_ or the cell encoder. At inference, the decoder outputs *μ*^all^, so the reconstructions used as post-training targets covered all *G* genes. The autoencoder used by the *all-folds* models was trained without a mask. In pseudocode:

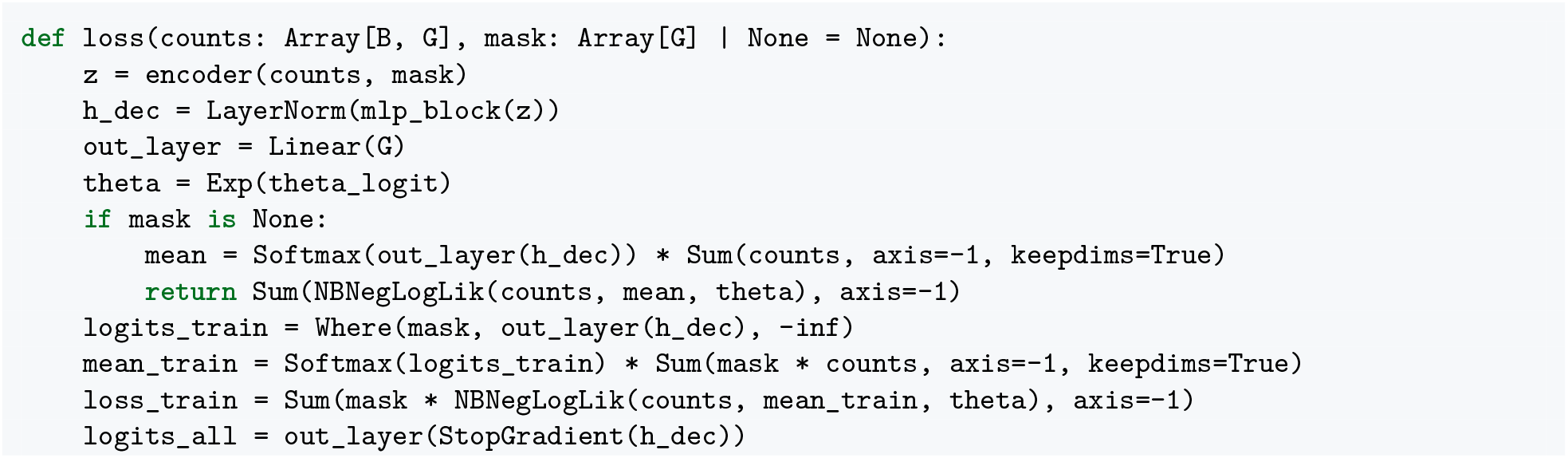
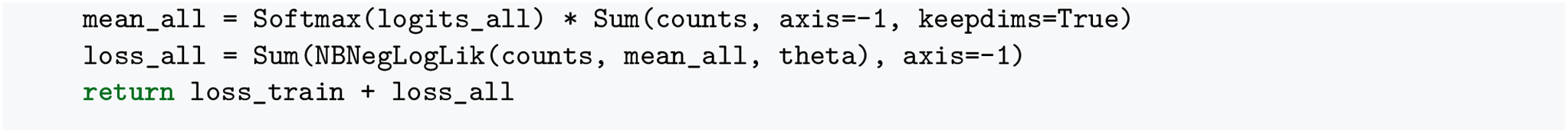

#### Data split

The autoencoder was trained on a human scRNA-seq corpus combining CZ CELLxGENE Discover^2^ and our human single-cell pre-training datasets (Single-cell RNA-seq data processing for cell encoder training and post-training), together with unlabeled human basal ganglia profiles from HMBA^116^ (excluded from Stage 4 post-training).

For the *fold-1* models, CELLxGENE datasets published after 1 October 2025 formed the temporal test set, and eight datasets across six CELLxGENE collections (healthy skin, tonsil, COVID-19 lung from three donors, developing cerebellum, developing bone marrow, and healthy kidney) formed the validation set. All other non-test datasets from these six collections, together with an immune aging atlas collection, were excluded from training, and all remaining datasets formed the training set. For the *all-folds* models, the autoencoder was trained on all datasets in the corpus, including the validation, test, and held-out collection datasets.

#### Optimization

Training used AdamW (*β*_1_ = 0.9, *β*_2_ = 0.99, *ε* = 5 × 10*^−^*^7^) with a weight decay of 1.0 applied to linear kernel weights only, and gradients were clipped to a global norm of 1000. The learning rate warmed up linearly to 0.001 over 10,000 steps and followed a cosine decay to 0 over the remaining steps. Training ran for 250,000 steps on 16 TPU v4 chips with a global batch size of 1024 cells sampled uniformly across training cells. Cell embeddings and denoised reconstructions were pre-computed once and stored for Stage 4 post-training.

### Evaluated model configurations

Downstream evaluations used four AG2 model configurations matched to each task (Data splits):

- ***fold-1* teacher and pre-trained models (Stages 1–2; Stage 1 for ablations):**Trained on the *fold-1* genomic interval split with all reference tracks observed. A teacher model (Stages 1–2) was used for held-out *fold-1* test-interval evaluations across reference conditions (Fig. 2a,b, Supplementary Fig. 3), whereas Stage 1 pre-trained *fold-1* models (*n* = 3 replicate seeds per configuration, without Stage 2 condition refinement) were used for the conditioning pathway ablations (Supplementary Fig. 4) alongside an unconditioned AG1 multitask baseline retrained on matched bulk datasets.
- ***fold-1* post-trained distilled student (Stages 1–4, temporal split):** A *fold-1* distilled student model (Stages 1–3) extended in Stage 4 with the gene-masked, temporal-split single-cell expression autoencoder (Single-cell expression autoencoder; standalone denoising evaluated in Supplementary Fig. 9c). Used to evaluate generalization to seen and chronologically held-out CELLxGENE scRNA-seq datasets on held-out *fold-1* intervals (Fig. 3b,c, Supplementary Fig. 9d,e), compared against per-group fine-tuning of the same *fold-1* distilled student and against Stage 4 post-training of a single Stage 1 pre-trained *fold-1* model.
- ***all-folds* imputation-split distilled student (Stages 1–3):** Trained across all eight genomic folds with validation, test, and withheld bulk PBMC and astrocyte tracks masked out (Imputation split). Used for bulk cross-modality imputation on *fold-5* intervals (Supplementary Fig. 7a–c, Supplementary Fig. 8), primary PBMC H3K27ac enhancer classification (Supplementary Fig. 7d,e), TenK10K single-cell QTL scoring without bulk PBMC data (“AG2 (no bulk PBMC)”; Supplementary Fig. 5e), and astrocyte CRISPRi E2G linking without bulk astrocyte data (“AG2 (no bulk astrocytes)”; Fig. 4c).
- ***all-folds* post-trained distilled student (Stages 1–4, full corpus):** The primary model trained across all eight genomic folds and all reference tracks (Stages 1–3) and extended in Stage 4 with the unmasked all-datasets single-cell expression autoencoder (Stage 4: Post-training for expression conditioning), enabling the same frozen sequence backbone to be queried via either reference conditions or expression conditioning. Used for all remaining evaluations: variant effect benchmarks on held-out test chromosomes and locus vignettes (Fig. 2c–h, Supplementary Fig. 2, Supplementary Fig. 5), SARS-CoV-2 vaccination and Parkinson’s disease cohort analyses (Fig. 3d–j, Supplementary Fig. 10), CRISPRi E2G benchmarks and the 640-condition E2G atlas (Fig. 4b–k, Supplementary Figs. 11 to 13), and TraitGym and the 510-condition genome-wide association study (GWAS) variant effect atlas and metacell dissections (Fig. 5, Supplementary Fig. 14, Supplementary Fig. 15).

### Interval evaluations

A central goal of AG2 is to predict how regulatory signals vary across conditions, that is, which genes or regulatory elements are differentially active between tissues, cell lines, or cell types. To measure this, we evaluated how accurately the model predicts condition-specific variation on held-out genomic intervals (Data splits); all results in this section are reported on held-out *fold-1* test intervals (Fig. 2a,b, Supplementary Fig. 3, Supplementary Fig. 4).

#### Interval scoring

For each evaluation, the model was conditioned on every constituent reference condition (tissue, cell line, or cell type) in a comparison group. For each 1-Mb (2^20^ = 1,048,576 bp) input sequence, spatial track predictions were cropped to the central 196,608-bp (3×2^16^ bp) target window (discarding 425,984 bp from each flank) to avoid sequence-edge effects, and per-feature values were extracted for annotated features completely contained within this central window:

- **Bulk RNA-seq.** For each protein-coding gene, base-pair-resolution predictions were summed across annotated exons of its protein-coding transcripts, scaled from coverage to reads per million (dividing by the 100-bp read-length factor used in track normalization), and divided by exonic length in kilobases to yield a reads-per-kilobase-per-million-like activity value.
- **Single-cell RNA-seq.** Per-gene expression predictions (in CPM) were read directly from the single-cell gene expression output head.
- **Single-cell ATAC-seq.** Per-peak accessibility was obtained by averaging base-pair-resolution scATAC-seq predictions across each consensus peak region.
- **ChIP-seq (TF and histone).** Per-peak signal was obtained by averaging 128-bp ChIP-seq predictions across each consensus peak region for the target TF or histone modification.

#### Evaluation design

We defined the following evaluations across modalities:

- **Bulk RNA-seq.** We compared all GTEx tissues (Fig. 2a, Supplementary Fig. 4b) or 18 paired-end ENCODE total RNA-seq cell lines (Supplementary Fig. 4b). Each 1-Mb input sequence was centered on the midpoint of a protein-coding gene (excluding chromosomes Y and M), omitting intervals whose central 196,608-bp window overlapped *fold-1* training or validation intervals; protein-coding genes completely contained within each central 196,608-bp window were evaluated, deduplicating genes across overlapping windows by retaining the window centered closest to the gene midpoint.
- **Single-cell RNA-seq.** We compared groups of cell types drawn from the same tissue, lineage, assay, and source dataset (Fig. 2b, Supplementary Fig. 3), such as lung epithelial cell types assayed with 10x 3’ single-nucleus RNA-seq (snRNA-seq) or peripheral blood immune cell types assayed with 10x 5’ scRNA-seq, using the same gene-centered intervals and deduplication procedure as for bulk RNA-seq.
- **Single-cell ATAC-seq.** We compared groups of cell types drawn from the same tissue, lineage, assay, and source dataset (Fig. 2b, Supplementary Fig. 3), analogous to scRNA-seq. Within each dataset, SnapATAC2^106^ (MACS3^123^) peak calling was run on raw fragments grouped by cell type (excluding cell types with fewer than 50 cells), and called peaks were merged across cell types within that dataset into a unified per-study consensus peak set. Consensus peaks completely contained within the central 196,608-bp windows of held-out *fold-1* test intervals were evaluated.
- **ChIP-seq (TF and histone).** Each comparison group comprised all ENCODE EFO conditions profiled for a given TF or histone modification (Supplementary Fig. 4b). We evaluated 10 TFs (CEBPB, CTCF, MAX, POLR2A, POLR2AphosphoS5, RAD21, REST, SIN3A, TAF1, and YY1) and 11 histone modifications (H3K4me3, H3K4me1, H3K36me3, H3K27ac, H3K9ac, H3K27me3, H3K4me2, H3K79me2, H2AFZ, H3K9me3, and H4K20me1), each profiled across more than 10 EFO conditions. When both single-end and paired-end tracks were available for a given EFO condition and target TF or histone modification, the paired-end track was selected. For each target, consensus peak regions were defined by merging called ENCODE BED peaks across all profiled EFO conditions using SnapATAC2^106^. To focus evaluation on condition-specific regulatory elements, 250 active consensus peaks completely contained within the central 196,608-bp windows of held-out *fold-1* test intervals were selected per profiled condition without replacement across conditions (ranking peaks by the fewest active conditions, with ties broken randomly), and predictions were extracted across all *fold-1* test intervals containing these selected peaks.

Conditioning the model on each of the *T* reference conditions in a comparison group across *N* evaluated features (genes or consensus peaks) yielded a predicted activity matrix 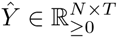 with entries *Ŷ_i,t_*, which was compared against the corresponding observed activity matrix 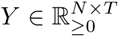 with entries *Y_i_*_,*t*_. The complete list of single-cell comparison groups and their constituent cell types is provided in Supplementary Table 8.

##### Gene expression and chromatin accessibility log fold change

For each feature *i* (gene or peak) and condition *t*, relative activity was computed as the log fold change (LFC) by centering log-transformed activity against the mean log-transformed activity of that feature across all *T* conditions in the comparison group:

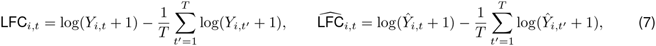

where *Y_i,t_* and *Ŷ_i,t_* are the entries of the observed and predicted activity matrices *Y* and *Ŷ* defined above. For stranded RNA-seq, the plus- and minus-strand tracks of each sample were treated as a single track, with each gene’s activity taken from the strand matching the gene’s orientation; unstranded tracks were used for genes on either strand. For each condition *t* (one track per condition), Pearson *r* was computed between predicted and observed LFCs across all *N* features, measuring how accurately the model captured differential up- and downregulation across conditions. The metric reported for each comparison group is the mean of these per-track Pearson *r* values across all conditions in the group (including in Supplementary Fig. 4b); in Fig. 2a, where the distribution across GTEx tissues is shown, we report the median per-tissue Pearson *r*.

##### Cross-condition variation explained

To summarize how much of the observed variation between conditions within a comparison group the model captured, we computed the fraction of variance in observed LFCs explained by predicted LFCs, pooled over all features and conditions in the group:

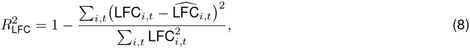

reported as a percentage (Fig. 2a,b), with performance across single-cell comparison groups summarized by the median (Fig. 2b). Because LFCs are centered per feature, the denominator measures the variation in observed log-activity remaining after removing each feature’s mean across conditions. This measure can be negative when predictions deviate from observations more than a constant (zero-LFC) prediction would.

##### ChIP-seq peak AUPRC

For ChIP-seq TF and histone modification tracks, condition specificity was evaluated using the area under the precision–recall curve (AUPRC) for peak detection over consensus peak sets. For each evaluated condition *t*, consensus peaks were ranked by their mean predicted signal across the region and labeled against ground truth as:

- **Positive peaks (***y* **= 1):** Regions containing an experimentally called peak in the evaluated condition.
- **Negative peaks (***y* **= 0):** Regions called as active peaks in other conditions within the consensus set, but lacking a called peak in the evaluated condition.

Evaluating classification against active peaks from other conditions, rather than unannotated genomic background, tests whether the model captures condition-specific regulatory activation and silencing beyond general sequence motifs. For the *fold-1* ChIP-seq evaluation (Supplementary Fig. 4b), AUPRC was computed across the selected consensus peaks for each condition and averaged across conditions within each target TF or histone modification; the reported score for each modality is the mean across its evaluated targets.

#### AG1 single-cell fine-tuned baseline

To construct a comparable single-cell baseline (AG1-sc), we fine-tuned AG1^14^ on the same human and mouse single-cell pre-training tracks as AG2 (Single-cell RNA-seq and ATAC-seq data), warm-starting from the pre-trained *fold-1* checkpoint for held-out interval evaluations (Fig. 2b, Supplementary Fig. 3) and from the distilled *all-folds* model for variant effect evaluations (Fig. 2e, Supplementary Figs. 5 and 6). We attached species-specific unconditioned multitask heads (as in AG1) predicting all single-cell tracks simultaneously and applied a stop-gradient operator at the inputs to all pre-trained bulk heads so that only single-cell losses updated the sequence trunk. Single-cell gene expression used the AG2 exon-pooled cross-attention head, target scaling (*s* = 48), and loss (Output heads; head loss weight 1.0, without weight dropout), while single-cell chromatin accessibility used an AG1 coverage head and loss^14^ (per-track target normalization, head loss weight 4.0).

Both models were fine-tuned on 256 TPU v4 chips (4×8×8 topology, batch size 64) for 10,000 steps using AdamW with weight decay 0.1 on kernel weights, global gradient norm clipping at 0.5, and trunk dropout 0.3. Single-cell heads warmed up linearly to a learning rate of 0.002 over steps 0–1000 and cosine-decayed to 0 over steps 1000–10,000, whereas pre-trained parameters were held at learning rate 0 for the first 1000 steps, warmed up linearly to 0.002 over steps 1000–2000, and cosine-decayed to 0 over steps 2000–10,000.

### Variant evaluations

#### Variant scoring

Across all variant evaluations, we scored genetic variants by centering a 1-Mb (2^20^ = 1,048,576 bp) input sequence on each variant and contrasting model predictions for sequences carrying the alternate (ALT) versus reference (REF) allele.

For existing bulk modalities (bulk RNA-seq, bulk ATAC-seq, DNase-seq, ChIP-seq, contact maps, and splicing; Supplementary Fig. 2), variant scoring was performed identically to AG1^14^. AG2 introduces scoring strategies for two new single-cell modalities:

##### Single-cell chromatin accessibility

Variants were scored using a center-mask scorer over a window *W*_1201_ of width 1201 bp centered on the variant (unless noted otherwise), summing predicted base-resolution signal across *W*_1201_ prior to log_2_(*x* + 1) transformation: 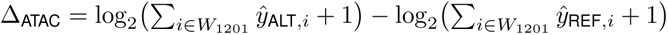.

##### Single-cell gene expressio

Unlike bulk RNA-seq coverage tracks, which aggregate predictions across annotated exons, the single-cell gene expression heads directly output scalar gene-level expression values (in CPM) for each annotated gene in the 1-Mb interval. For a target gene, the variant score is the predicted natural-log fold change Δ_expr_ = log(*ŷ*_ALT_ + 0.001) − log(*ŷ*_REF_ + 0.001).

##### *In silico* mutagenesis (ISM) for contribution scores

To construct ISM attribution logos, we followed the same procedure as in AG1^14^.

#### Bulk and single-cell eQTL evaluations

To evaluate AG2’s ability to predict variant effects on gene expression across tissues and cell types, we developed an eQTL benchmark suite from three resources: (1) bulk tissue eQTLs from GTEx v11^34^, (2) single-cell immune eQTLs from the TenK10K study^7^, and (3) single-cell intestinal eQTLs from IBDverse^38^ (Fig. 2c–e, Supplementary Figs. 5 and 6). Across these datasets, evaluations were developed from three predictive tasks: causality classification, effect direction (sign) prediction, and effect magnitude (coefficient) correlation. All evaluations are reported on variants on held-out test chromosomes (Variant benchmarks split) except for IBDverse sc-eQTL evaluations (Single-cell eQTLs).

##### Bulk tissue eQTLs (GTEx v11)

For bulk tissue evaluations, we used SuSiE fine-mapped eQTLs from the GTEx Analysis v11 release^34,71^. Variants were restricted to single-nucleotide variants (SNVs). When a variant–gene pair appeared multiple times within the same tissue in the SuSiE fine-mapping tables, we deduplicated entries by retaining the association with the highest posterior inclusion probability (PIP).

AG2 was conditioned on the target tissue ontology term, and variant effects were scored using the RNA-seq gene variant scorer on the output channel corresponding to the recount3^97^ processing pipeline.

##### Single-cell eQTLs

We evaluated single-cell *cis*-eQTL (sc-eQTL) predictions using SuSiE fine-mapped associations from the TenK10K^7^ and IBDverse^38^ cohorts. AG2 was not trained on data from either study; scRNA-seq reference conditions in the training corpus were derived from independent atlases (Kock *et al.*^108^ for most of blood conditions; Elmentaite *et al.*^124^ for ileum and rectum conditions; and Elmentaite *et al.*^124^ and ENCODE^46^ for colon conditions). We excluded eGenes with no detectable expression in any of the mapped reference conditions, ensuring that variant effect predictions were evaluated only for genes the model can meaningfully score in a given tissue (blood or intestinal). For evaluation, each cell type was mapped to a training reference condition (Supplementary Tables 9 and 10), and variant effects were scored using the single-cell gene expression scorer on the channel corresponding to that reference dataset during training.

*TenK10K.* We evaluated sc-eQTLs across 28 PBMC cell types^7^, aligning cell-type labels to the Kock *et al.* PBMC reference conditions (Supplementary Table 9), except for hematopoietic stem and progenitor cells (HSPCs), which were conditioned on the matching bone marrow reference condition^125^. To assess the contribution of bulk data from purified blood cell types, we also evaluated the separately trained imputation-split model (Imputation split; “AG2 (no bulk PBMC)”), which excluded all bulk PBMC tracks from training together with the imputation validation and test tracks, conditioned and scored identically (Supplementary Fig. 5e).

*IBDverse.* sc-eQTLs were derived from 72 cell types from the terminal ileum, rectum, and peripheral blood of individuals with and without inflammatory bowel disease^38^, as processed by the eQTL Catalogue (release r8; study QTS000045)^39^. Because each eQTL Catalogue dataset pooled cells of a given cell type across the sampled tissues of each donor, we assigned each cell type to its donor-majority site in the IBDverse single-cell atlas: for each cell type, we retained donors with at least 20 cells of that cell type pooled across tissues, computed the fraction of each donor’s cells originating from each tissue, and selected the tissue with the highest mean fraction across retained donors. As peripheral blood cell types were already covered by the larger, better-powered TenK10K cohort, we restricted this benchmark to the ileum and rectum and excluded cell types whose donor-majority site was blood. Cell-type mappings aligned IBDverse labels to reference conditions in the donor-majority tissue, or to the closest cell type from a related tissue where no match was available (Supplementary Table 10).

We evaluated IBDverse across all chromosomes rather than restricting to held-out test chromosomes (Variant benchmarks split) because its smaller donor cohort yielded fewer high-confidence fine-mapped variants than GTEx or TenK10K. To prevent overlap with variants used during model and scorer selection from other studies, we first excluded all variants (both positive and negative) appearing on non-test chromosomes in the GTEx eQTL, GTEx splicing QTL (sQTL), TenK10K sc-eQTL, and TenK10K single-cell chromatin accessibility QTL (sc-caQTL) causality benchmarks. We then removed cell types with fewer than 10 high-confidence causal variants (PIP > 0.9). Together, these filtering steps retained 3524 causal sc-eQTLs across 47 intestinal cell types: 34 with the terminal ileum as donor-majority site (15 epithelial and 19 immune) and 13 epithelial cell types with the rectum as donor-majority site (Supplementary Table 10). IBDverse was not used for model or scorer selection (Model selection).

##### Causality evaluation

We evaluated AG2’s ability to discriminate causal regulatory variant–gene associations (PIP > 0.9) from non-causal pairs (PIP < 0.01) within each tissue or cell type.

Causal eQTLs are strongly enriched near TSSs and within active regulatory elements. To control for genomic location, variants within each GTEx tissue, TenK10K cell type and IBDverse cell type were partitioned by overlap with active regulatory elements, defined using ENCODE v4 SCREEN candidate *cis*-regulatory elements (cCREs)^40^ for GTEx, TenK10K MACS3 scATAC-seq peaks^37^ for TenK10K, and the union of TenK10K peaks and ENCODE human colon^46,126^ MACS3 scATAC-seq peaks for IBDverse. For IBDverse, because our scATAC-seq training corpus contained colon profiles but no terminal ileum or rectum datasets, we used 21 ENCODE colon tracks (10x Chromium and sciATAC-seq) as the intestinal chromatin accessibility reference, calling MACS3 peaks per experiment and merging them into 501-bp summit-centered intervals with SnapATAC2^106^ before taking the union with TenK10K blood peaks. Each causal SNV was matched without replacement to the non-causal control in the same stratum with the closest distance to its own target eGene TSS at a 1:1 ratio (GTEx: *n* = 67,674 total across 33,837 positives and 33,837 controls; TenK10K: *n* = 12,692 across 6346 positives and 6346 controls, IBDverse: *n* = 7048 across 3524 positives and 3524 controls). Here we defined the eGene TSS as the median TSS across all transcripts of that gene in the GENCODE V46 GTF, as done in AlphaGenome1^14^.

We computed a combined precision–recall curve by pooling across tissues in GTEx and cell types in TenK10K or IBDverse. We additionally report tissue/cell-type-weighted AUPRC as done in AG1.

##### Effect direction prediction

For high-confidence causal variants (PIP > 0.9), we assessed classification accuracy in predicting whether a variant increases or decreases expression of its target eGene (GTEx: *n* = 33,829, excluding eight associations lacking allelic fold-change estimates; TenK10K: *n* = 6346, IBDverse: *n* = 3524). We evaluated sign prediction performance across confidence thresholds by pooling instances across tissues or cell types, ranking variants according to their predicted effect magnitude, and computing sign accuracy as a function of recall (the fraction of top-ranked variants retained), summarized by the area under the sign accuracy–recall curve (AUC). Additionally, we computed the tissue/cell-type-weighted sign-prediction area under the receiver operating characteristic curve (AUROC) as done in AG1.

##### Effect magnitude correlation

For high-confidence causal variants (PIP > 0.9), we evaluated whether predicted effect scores correlate with observed effect sizes, measured as allelic fold change in GTEx and single-cell regression slope coefficients in TenK10K and IBDverse (GTEx: *n* = 33,829, TenK10K: *n* = 6346, IBDverse: *n* = 3524). We computed the tissue/cell-type-weighted Spearman *ρ* as in AG1.

##### Scooby baseline

For TenK10K, we compared against Scooby^21^, using the published model trained on OneK1K PBMC scRNA-seq. Scooby predicts single-cell RNA-seq coverage from a 524,288-bp input conditioned on a cell embedding. We obtained a pseudobulk prediction for each cell type by summing predictions over all OneK1K cells with that annotation, averaging forward and reverse-complement predictions. Each TenK10K cell type was mapped to the closest of the 21 OneK1K annotations (Supplementary Table 9); eight had only approximate matches (naive and intermediate B cells to B cells; innate lymphoid cells and proliferating natural killer (NK) cells to NK cells; AXL^+^ dendritic cells (ASDCs) and type 1 conventional dendritic cells (cDC1s) to dendritic cells; proliferating CD4^+^ and CD8^+^ T cells to the corresponding effector memory cells). Inputs were centered on the variant, and variant effects were scored as the log fold change, with a pseudocount of 1, of predicted counts summed over the eGene’s exons.

Because Scooby predicts only the central 196,608 bp of its input, eGenes with no exon in this window received a score of zero; this affected 3.4% of causality and 3.6% of sign/coefficient pairs, all with the variant more than 35 kb from the TSS, so the 0–35 kb strata (Supplementary Fig. 6a) compare all models on fully scored pairs.

##### Tissue-specificity stratification

To stratify tissue-specific versus nonspecific regulatory effects (Fig. 2d), GTEx tissues were grouped by hierarchical average-linkage clustering on log-transformed median gene expression profiles (cosine distance cutoff 0.1). A causal variant–gene pair (PIP > 0.9) was labeled *tissue-specific* (*n* = 7169) if it showed PIP > 0.01 in only a single tissue cluster and *nonspecific* (*n* = 26,660) otherwise.

##### Single-cell eQTLs with expression conditioning

In addition, we evaluated the post-trained model (Stage 4: Post-training for expression conditioning) on both the sign prediction and causality benchmarks for the Tenk10k and IBDverse sc-QTLs. For Tenk10k, we supplied cell embeddings derived from an independent PBMC scRNA-seq dataset unseen during pre-training (Gong *et al.*^53^) in place of learned reference condition embeddings (Supplementary Fig. 10a,b). We subsampled this dataset to 1,000,000 cells, passed each cell through the cell encoder, and averaged the resulting cell embeddings across all cells assigned to the same cell type to form the mean cell embedding. Because this secondary PBMC dataset only contained cell-type matches for 25 of the 28 TenK10K cell types, the evaluation was restricted to this 25-cell-type subset. To assess sensitivity to the number of cells averaged into the mean cell embedding, we also repeated both evaluations using random subsamples of 1, 16, and 128 cells per cell type (10 independent trials per sample size). For IBDverse, we supplied the scRNA-seq data derived from the IBDverse study^38^, averaged embeddings across cells from each cell type’s donor-majority intestinal tissue, and evaluated the same 47 cell types described above (Single-cell eQTLs, Supplementary Fig. 10c,d).

#### Single-cell chromatin accessibility QTL (caQTL) evaluations

We used SuSiE fine-mapped *cis*-caQTLs from the TenK10K study^37^ (Supplementary Fig. 5b–e), where the target regulatory feature is an ATAC-seq peak rather than a gene. As with single-cell eQTLs, AG2 was not trained on TenK10K data; PBMC single-nucleus ATAC-seq profiles in our reference training corpus came from Benaglio *et al.*^110^. For evaluation, models were conditioned on the target cell-type ontology term, and variants were scored using the single-cell chromatin accessibility center-mask scorer on the channel corresponding to the reference dataset. Cell-type mappings represent a best-effort alignment between TenK10K labels and the reference condition set (Supplementary Table 9); as with sc-eQTLs, HSPCs were conditioned on the equivalent bone marrow cell type in the reference corpus. Cell types with no fine-mapped positive variants (PIP > 0.9) residing in their target peak in either the validation or test split were excluded from evaluation, leaving 24 evaluated PBMC cell types.

Two evaluation tasks were defined on held-out test chromosomes:

1. **Causality classification:** Causal SNVs (PIP > 0.9) residing within their target peak (*n* = 5116) were contrasted against non-causal in-peak SNVs (allele frequency >0.1, nominal *P* > 0.1, and absent from SuSiE credible sets) drawn from peaks in the same cell type that contained at least one nominally associated variant (*P* < 0.01). Within each cell type, controls were matched 4:1 without replacement on distance to their respective peak midpoint (*n* = 25,580 variant-cell-type pairs total).
2. **Effect-size correlation:** Cell-type-weighted Spearman rank correlation (*ρ*) between predicted scores and observed caQTL slopes across all causal SNVs (PIP > 0.9, SNP inside its target peak; *n* = 5,116).

#### Variant effect benchmarks shared with AG1

For the remaining bulk variant effect benchmarks (GTEx splicing QTLs (sQTLs), rare splicing outliers, chromatin accessibility and DNase sensitivity QTLs, and GTEx polyadenylation QTLs; Supplementary Fig. 2), datasets, variant scorers, and metrics followed AG1^14^, evaluated on held-out test chromosomes (Variant benchmarks split) with AG2 conditioned on the corresponding tissue or cell-type reference condition. For polyadenylation QTL (paQTL) causality on genes with multiple polyadenylation sites (PASs; 400-bp PAS mask), positive fine-mapped variants within cumulative distance thresholds of 50 bp to 10,000 bp from the target PAS were matched 1:1 to distance-matched negative controls across *n* = 100 random negative-sampling permutations to obtain per-threshold AUPRC distributions (Supplementary Fig. 2e).

#### Locus-level variant vignettes

Locus-level variant analyses in Fig. 2f–h used the *all-folds* distilled AG2 model:

- ***IL20RA* (rs62420789, `chr6:137026715:G>A`):** Bulk tissue variant scores were computed across seven GTEx tissues as in the GTEx eQTL benchmark (Bulk and single-cell eQTL evaluations), alongside single-cell gene expression variant scores and observed single-cell log(CPM + 1) expression of *GRHL2* and *ZEB1* across constituent reference conditions. Saturated ISM of predicted *IL20RA* expression (10x 3’ nuclear channel) was computed over a 64-bp window in lung goblet and lung blood vessel endothelial cells and displayed across `chr6:137026698-137026732` (Fig. 2f). The list of reference conditions used to construct the heatmap is given in Supplementary Table 11.
- ***FBXO31* (rs8052244, `chr16:87368655:G>C`):** Conditioned on peripheral blood CD14^+^ monocytes (`UBERON:0000178_CL:0001054`), we predicted reference and alternate local scATAC-seq (`chr16: 87368155-87369155`) and bulk RNA-seq (`chr16:87326986-87389142`) coverage, gene-body H3K27me3 log fold change across *FBXO31*, and saturated ISM over a 64-bp window (displayed across `chr16: 87368643-87368687`) using the 1201-bp scATAC-seq center-mask scorer and the single-cell gene expression scorer (10x 5’ whole-cell channel) for *FBXO31* (Fig. 2g).
- ***HK1* congenital hyperinsulinism:** Six disease-associated *de novo* non-coding SNVs within the *HK1* intronic regulatory region^42^ were compared against 80 common intronic variants from gnomAD v4.1.1 (allele frequency >0.01, `intron_variant`) within 5 kb of `chr10:69348900` (`chr10:69343901-69353899`), scoring predicted *HK1* single-cell gene expression log fold changes averaged across observed 10x 3’ single-cell expression channels in seven pancreatic and neuroendocrine primary reference conditions (Fig. 2h). The list of reference conditions used is given in Supplementary Table 12.

### Single-cell expression autoencoder and post-trained model evaluations

#### Comparison of the post-trained model versus fine-tuning

To evaluate the post-trained model’s ability to reconstruct cell-type specificity of gene expression on held-out genomic intervals (Fig. 3b,c, Supplementary Fig. 9d,e), we compared expression conditioning (Stage 4: Post-training for expression conditioning) against a baseline model fine-tuned directly on each dataset. We evaluated both approaches on a benchmark of single-cell RNA-seq datasets from CELLxGENE that were not part of the AG2 pre-training corpus, spanning datasets both seen and unseen during cell encoder training and AG2 post-training.

##### Benchmark construction

Starting from all human CELLxGENE datasets consisting entirely of primary data, we retained those containing data with an assay and suspension type that matched a combination present in the AG2 single-cell gene expression channels. Within each dataset, we selected the dominant (tissue, assay, suspension type) combination and did a best-effort manual grouping of its cell types into five broad lineages: immune, neural, epithelial, endothelial, and stromal. Any lineage containing at least five cell types with at least 50 cells each formed an evaluation group; if a lineage had more than 10 cell types, we kept only the 10 most abundant. Each evaluation group therefore compares 5–10 related cell types that share the same dataset, tissue, assay, suspension type, and lineage. Because datasets within a CELLxGENE collection often come from the same study and samples, we kept only one evaluation group per lineage and collection: the one with the most cell types, with ties broken by the total number of cells. A single dataset can still contribute evaluation groups for several lineages. This procedure yielded 167 evaluation groups in total: 147 (from 120 datasets in 118 collections) in the cell encoder training split and 20 (from 16 datasets in 16 collections) in a held-out split of datasets published after 1 October 2025 (Supplementary Table 13). We note that the biased distribution of tissues in CELLxGENE is reflected in this evaluation.

##### Post-trained model

For each group of cells within a given cell type, tissue, assay, and suspension type, we passed all constituent cells (or, for datasets exceeding one million cells, a cell-type-stratified subsample of up to one million cells) through the cell encoder and averaged the resulting cell embeddings. We then conditioned the *fold-1* post-trained model on these mean cell embeddings to generate predictions.

##### Fine-tuning baseline

For each of the 167 evaluation groups, we fine-tuned the *fold-1* distilled AG2 model independently on the single-cell gene expression data of its source dataset. The reference condition embedding table was re-initialized from scratch with entries for only the cell types in that group. The reference condition embedding table and the FiLM layers were unfrozen while all remaining parameters were held fixed. This configuration outperformed unfreezing only the reference condition embeddings on the LFC prediction task across the eight CELLxGENE validation datasets (Single-cell expression autoencoder). Training used a batch size of 64 for 5000 steps at a constant learning rate of 10*^−^*^4^ without warmup, with transformer and sequence-to-pair dropout disabled (retaining 0.1 weight dropout in the single-cell gene expression head) and only the single-cell gene expression head active, restricted to *fold-1* training intervals.

##### Base model comparison

We additionally post-trained a single Stage 1 pre-trained *fold-1* model (prior to Stage 2 condition refinement; labeled “AG2 Post-trained from teacher”) using the identical Stage 4 procedure and compared it against post-training the distilled student (Supplementary Fig. 9d,e).

##### Evaluation

Both approaches were scored using the cross-gene LFC Pearson correlation (Gene expression and chromatin accessibility log fold change), computed across the cell types within each evaluation group. Metrics are reported on held-out *fold-1* validation intervals, which were not used for fine-tuning or model selection in these comparisons.

#### Parkinson’s disease cohort sc-eQTL and caQTL evaluations

To evaluate variant effect prediction in an unseen disease cohort, we benchmarked AG2 on caQTLs and eQTLs from the substantia nigra and cingulate cortex in a Parkinson’s disease single-nucleus multiome study (Sigalova *et al.*^49^). Variants were evaluated natively on the Telomere-to-Telomere (T2T-CHM13v2.0) assembly and gene annotation without liftover to GRCh38.

##### Conditioning

Gene identifiers in the cohort’s 10x Chromium snRNA-seq count matrices were mapped to the cell encoder’s GENCODE v46 Ensembl vocabulary (protein-coding and lncRNA genes), and cell embeddings from the cell encoder (unmasked *all-folds* model) were averaged within each (brain region, cell type) pair to form the mean cell embedding (Stage 4: Post-training for expression conditioning).

##### Baselines

We compared against: (i) the published sequence-model predictions of Sigalova *et al.*^49^ (CREsted for caQTLs and Scooby for eQTLs), and (ii) the *all-folds* distilled AG2 model conditioned on the closest brain reference condition in our training corpus (Supplementary Table 14: dorsolateral prefrontal cortex, UBERON:0009834, across 23 cingulate cortex cell types mapping to 21 distinct reference conditions; midbrain, UBERON:0001891, for seven substantia nigra cell types; and primary motor cortex, UBERON:0001384, for substantia nigra astrocytes).

##### Chromatin accessibility QTLs (caQTL-ASCA)

We evaluated the high-confidence caQTL-ASCA callset of Sigalova *et al.*, comprising variants with concordant effects (false discovery rate, FDR < 0.1) in both across-individual *cis*-caQTL mapping and within-individual allele-specific chromatin accessibility (ASCA). Sign prediction was evaluated against the reported allelic imbalance direction (*n*_pos_ = 43,526, *n*_neg_ = 43,070) using the 1201-bp single-cell chromatin accessibility center-mask scorer (Variant scoring), averaged across channels.

##### Expression QTLs

We evaluated *cis*-eQTLs (FDR < 0.05 across substantia nigra and cingulate cortex cell types) that overlapped the caQTL-ASCA callset in the same (variant, cell type, brain region) tuple, retaining the minimum *P* -value variant per (gene, cell type, brain region) triplet. Restricting to target eGenes within the 1-Mb scoring window yielded 1620 eQTLs (of 1739 total; *n*_pos_ = 846, *n*_neg_ = 774). Variants were scored with the single-cell gene expression scorer (Variant scoring), averaged across channels for reference conditioning or read from the target-strand channel of the denoised single-cell gene expression head for expression conditioning.

##### Scooby comparison

Sigalova *et al.* scored each caQTL-ASCA variant against all genes whose 524-kb Scooby input window (centered on the gene body) contained the variant, reporting only predictions with LFC > 0.05 (the fine-tuned Scooby model weights were not publicly available). We harmonized cell-type labels between the Scooby predictions and the eQTL set, collapsed duplicate entries that differed only in peak identifier and had identical LFC, and matched predictions to eQTLs on (variant, gene, cell type, brain region), yielding 276 Scooby-scored eQTLs. eQTLs without a reported Scooby prediction were treated as abstentions, so Scooby was represented by a single operating point: its recall was the fraction of eQTLs with a reported prediction, and its sign accuracy was computed over those eQTLs.

##### Evaluation

For both tasks, sign accuracy was computed by pooling all (variant, cell type, brain region) instances, ranking by predicted effect magnitude, and evaluating cumulative sign accuracy as a function of recall. Published Scooby predictions were filtered at LFC > 0.05; the fraction of eQTLs retained by this filter defined Scooby’s recall, and AG2’s recall was read off its sign accuracy–recall curve at Scooby’s sign accuracy.

#### Imputing accessibility and predicting variant effects in vaccine-induced CD8^+^ T cells

To evaluate whether post-trained AG2 can impute unseen epigenetic modalities and resolve cell-state-specific non-coding variant effects purely from single-cell transcriptomic profiles, we analyzed a held-out multimodal PBMC SARS-CoV-2 mRNA vaccination atlas comprising paired CITE-seq (scRNA-seq and surface protein) and ASAP-seq (scATAC-seq and surface protein) profiles^48^. All evaluations used post-vaccination PBMC samples pooled across days 11 and 28 across annotated level-2 T-cell states, including vaccine-induced group A CD8^+^ T cells (VI-A CD8^+^ T), effector memory CD8^+^ T cells (CD8 TEM), and naive CD8^+^ T cells (CD8 Naive).

##### Conditioning

As in the Parkinson’s disease cohort, we mapped CITE-seq scRNA-seq gene symbols to the cell encoder’s GENCODE v46 Ensembl vocabulary (protein-coding and lncRNA genes), passed all post-vaccination cells through the cell encoder, and averaged the resulting cell embeddings within each level-2 T-cell state to form its mean cell embedding (Stage 4: Post-training for expression conditioning).

##### Consensus peak calling and differential accessibility evaluation

Observed ASAP-seq fragment files were processed with SnapATAC2^106^ to call per-population peaks across level-2 PBMC cell states and merge them into 236,854 501-bp consensus peaks; observed counts were summed per T-cell state and normalized to CPM. We conditioned the post-trained AG2 model on the mean cell embedding of each T-cell state and imputed base-resolution scATAC-seq tracks across all genome-wide evaluation intervals using the 10x nuclear scATAC-seq channel. Predicted accessibility for each peak was obtained by averaging the base-pair-resolution predictions over the consensus peak region, and was then normalized to CPM across peaks. We restricted the evaluation to peaks predicted to be accessible (CPM ≥5 in VI-A CD8^+^ T or CD8 TEM; *n* = 37,035) and computed the log fold change between VI-A CD8^+^ T and CD8 TEM as LFC = log(CPM_VI-A_ + 1) log(CPM_TEM_ + 1), separately for observed and imputed accessibility. We report the Pearson correlation between observed and imputed LFCs across these peaks (Fig. 3e).

##### Locus analysis at rs13066062

To dissect the COVID-19 severity-associated variant rs13066062 (`chr3: 45976852:G>A`), we imputed base-resolution scATAC-seq tracks and 128-bp-resolution histone ChIP-seq tracks across a 1-Mb interval centered on the variant conditioned on the post-vaccination mean cell embeddings for CD8 Naive, CD8 TEM, and VI-A CD8^+^ T cells. Observed ASAP-seq CPM bigWig tracks and imputed scATAC-seq tracks were extracted across a 10-kb window (`chr3:45971852-45981852`), smoothed with a 128-bp uniform moving average, and min–max scaled to [0, 1] across the three CD8^+^ T-cell states independently for observed and imputed tracks (Fig. 3f). Imputed H3K27ac tracks (averaged across H3K27ac channels) were extracted unsmoothed over the same 10-kb window on a shared raw signal axis without cross-state normalization (Supplementary Fig. 10k). Saturated ISM was performed across the 41-bp window surrounding rs13066062 (`chr3:45976832-45976873`) conditioned on VI-A CD8^+^ T cells using the 1201-bp single-cell chromatin accessibility center-mask scorer (Variant scoring). Cell-state-resolved variant scores (ALT A REF G) across CD8 Naive, CD8 TEM, and VI-A CD8^+^ T cells were computed with the denoised single-cell gene expression scorer for *CXCR6* and *FYCO1* (Supplementary Fig. 10i) and the 2001-bp histone ChIP-seq center-mask scorer (averaged across channels per modification) across H3K27ac, H3K4me1, H3K4me3, H3K36me3, H3K27me3, and H3K9me3 (Variant scoring; Supplementary Fig. 10k).

##### Metacell-resolved variant effects at rs13066062

To resolve variant effects across all post-vaccination PBMCs, we embedded day 11 and day 28 CITE-seq cells with the cell encoder after randomly subsampling each level-2 cell type to at most 10,000 cells (affecting only CD14^+^ monocytes). Within each level-2 cell type, cells were partitioned into *k* = max(1, round(*n*/64)) metacells by *k*-means clustering of their cell embeddings (scikit-learn^127^), yielding 881 cell-type-pure metacells (including two VI-A CD8^+^ T metacells spanning 115 cells). Conditioning post-trained AG2 on each metacell’s mean cell embedding, we scored rs13066062 over a 1-Mb interval centered on the variant using the 1201-bp single-cell chromatin accessibility center-mask scorer (10x nuclear scATAC-seq channel) and the denoised single-cell gene expression scorer for *CXCR6* (Variant scoring; Fig. 3g,h, Supplementary Fig. 10j). For visualization, a uniform manifold approximation and projection (UMAP; umap-learn^50^, `min_dist` = 0.5, other parameters default) was fitted on the per-cell embeddings. Each cell was colored by its metacell’s predicted effect, and each metacell was placed at the UMAP projection of its mean embedding. Metacells were ranked by predicted effect on *CXCR6* expression, with rank 1 the largest predicted decrease.

#### Single-cell autoencoder evaluation

We compared 256-dimensional cell embeddings from the cell encoder (unmasked *all-folds* model) against an scVI model^51^ trained directly on the CITE-seq RNA component of the SARS-CoV-2 vaccination PBMC atlas of Zhang *et al.*^48^ across all four timepoints (*n* = 113,897 cells across six donors and 30 level-2 cell types; Supplementary Fig. 9a,b), using level-2 cell-type annotations as biological labels and donor identity as the batch variable.

##### Representations

For AG2, each cell was passed directly through the cell encoder without dataset-specific training or batch covariates. For scVI (scvi-tools^112^), we selected the top 2000 donor-batch-aware highly variable genes directly from raw UMI counts in Scanpy^114^ (Seurat v3 flavor^115^) and trained a two-layer, 30-dimensional model with a negative binomial likelihood and donor batch covariate under default hyperparameters. Both latent spaces were projected into two dimensions with UMAP in Scanpy^114^ (Supplementary Fig. 9b).

##### Benchmarking

We evaluated biological conservation (*k*-means normalized mutual information and adjusted Rand index, cell-type local inverse Simpson’s index (cLISI), and cell-type silhouette width) and donor batch mixing (batch-removal-adapted silhouette score^128^) using scib-metrics^52^ without cross-method min–max rescaling (`min_max_scale=False`), reporting the unscaled biological conservation, batch correction, and overall (0.6 × biological + 0.4 × batch) scores (Supplementary Fig. 9a).

##### Cell encoder denoising across cell counts

To evaluate how many cells are required for the single-cell expression autoencoder to recover cell-type-specific expression profiles, we evaluated its denoised gene expression reconstructions across datasets from the held-out CELLxGENE temporal split (Comparison of the post-trained model versus fine-tuning, *n* = 16 datasets, Supplementary Fig. 9c). For each cell type and each subsample size **k** ∈ {1, 2, 4, 8, 16, 32, 64}, we performed 20 independent random subsampling trials without replacement, averaging the embedding produced by the cell encoder across the *k* sampled cells in each trial and then decoding into a single denoised profile; we also evaluated averaging over all available cells (All). Across all cell counts, these denoised profiles derived from averaged cell embeddings were evaluated against the full observed pseudobulk expression profile of that cell type using the cross-gene LFC Pearson correlation across cell types (Comparison of the post-trained model versus fine-tuning). For each held-out dataset, we averaged the correlation across the 20 subsampling trials at each *k*, reporting both individual dataset trajectories and the median across datasets.

### Imputation evaluations

#### Imputation metrics

Following the ENCODE Imputation Challenge^43^, evaluation metrics were chosen for (1) scale and offset invariance, (2) sensitivity to condition-specific peaks and expression differences, (3) stratification by assay channel and sequencing endedness, and (4) complementary assessment of discrete peak detection and continuous profile shape.

Scale-dependent regression metrics such as mean squared error (MSE) were omitted during final evaluation because technical scale and background differences penalize well-shaped profiles and because narrow histone marks (such as H3K4me3) exhibit higher native variance than broad marks (such as H3K9me3)^43^. Instead, we report genome-wide Pearson *r* for continuous profile shape alongside consensus peak AUPRC and cross-condition gene LFC Pearson *r* for condition specificity.

##### Channel stratification and sequencing endedness

Imputation metrics and baselines were stratified by AG2 output channel (Supplementary Table 6), where each channel is defined by experimental modality, data source, strand, ChIP target (transcription factor or histone modification), sequencing endedness (single-end vs. paired-end), and suspension type for single-cell data.

Consequently, for DNase-seq, ATAC-seq, and ChIP-seq, we treated single-end and paired-end data as distinct channels. Conflating single-end and paired-end tracks posed a major issue in the ENCODE Imputation Challenge^43^, where the distributional shift between single-end training data and paired-end test data required post-hoc reprocessing and quantile normalization. Modeling endedness as a channel attribute avoids cross-protocol distribution shifts and prevents technical artifacts from confounding evaluation.

##### Consensus peak definitions and peak AUPRC

For each chromatin channel (defined by assay, ChIP target where applicable, and sequencing endedness), we constructed a unified set of non-overlapping consensus peaks with SnapATAC2^106^ by merging ENCODE narrowPeak calls across all profiled conditions in that channel (spanning training and held-out splits). For each of the 176 held-out chromatin test tracks (spanning 25 of the 30 bulk test conditions, as the remaining five held out RNA-seq only; Supplementary Fig. 7a, Supplementary Fig. 8a), we scored each consensus peak by the mean of the predicted track across the 32-bp genomic bins spanning that peak (Metric computation) and computed the AUPRC for classifying peaks called in the held-out condition (positives) versus consensus peaks called only in other conditions of that channel (negatives), as in ChIP-seq peak AUPRC. Within each modality (ATAC-seq, DNase-seq, histone ChIP-seq, and TF ChIP-seq), per-track AUPRCs were aggregated via a weighted mean using each track’s positive peak fraction (the proportion of the channel’s consensus peaks active in that condition) as its weight, which we refer to as support-weighted peak AUPRC (Supplementary Fig. 7b).

##### Condition specificity and peak frequency stratification

To assess model accuracy across regulatory elements with varying degrees of biological selectivity (Supplementary Fig. 8b–e), consensus peak regions were stratified by empirical peak frequency, defined as the proportion of conditions in which a consensus peak is active across all profiled conditions in the assay channel. Peaks were partitioned into eight frequency bins of width 12.5%, ranging from rare, condition-specific elements (active in 0 < *f* ≤ 12.5% of conditions) to ubiquitous, constitutively open elements (active in >87.5%). Peak AUPRC was evaluated independently within each frequency bin using support-weighted aggregation across tracks, accompanied by an empirical random baseline reflecting the bin’s support-weighted positive peak prevalence. This stratification decouples performance on invariant regulatory landmarks from the model’s ability to capture condition-specific chromatin signal.

##### Gene LFC Pearson *r*

For bulk RNA-seq, genome-wide track correlations are dominated by housekeeping genes expressed across conditions. For each protein-coding gene (GENCODE v46) completely contained within the central 196,608-bp target window of a *fold-5*test interval, we summed 32-bp bin coverage across the full gene body, divided by the 100-bp read-length factor and gene length in kilobases, and clamped negative baseline predictions at zero. Merging plus- and minus-strand tracks per biosample as in Gene expression and chromatin accessibility log fold change yielded five evaluated RNA-seq tracks from the eight held-out tracks (181 evaluated tracks in total across all bulk modalities). Given that the imputation evaluation spans tracks from heterogeneous RNA-seq assays, we applied per-track uniform-quantile normalization across genes to the log(*x* + 1)-transformed expression values prior to per-gene centering across the held-out tracks, and then computed the mean per-track Pearson *r* as in Gene expression and chromatin accessibility log fold change.

##### Continuous signal correlations

For each held-out chromatin track, continuous Pearson *r* was computed between observed and predicted profiles across all evaluated 32-bp genomic bins and averaged across tracks within each modality (Supplementary Fig. 8f).

##### Metric computation

Evaluation metrics were computed per track at 32 bp resolution across the 1749 human *fold-5* test intervals (Imputation split). For each 1-Mb input sequence, spatial predictions were first cropped to the central 196,608-bp target window (discarding 425,984 bp from each flank, as in Interval evaluations). Predictions and observed targets at 1 bp resolution were downsampled to 32 bp via sum-pooling (with binary peak and gene-body masks max-pooled), while 128-bp ChIP-seq tracks were upsampled to 32 bp by bin repetition and dividing by the number of repeats to preserve total signal. Continuous 32-bp predictions and targets were concatenated across all central 196,608-bp windows to compute genome-wide Pearson *r*. For discrete elements (consensus peaks and protein-coding gene bodies), only features completely contained within each central 196,608-bp window were retained and concatenated across windows to compute peak AUPRC, frequency-stratified peak AUPRC, and gene LFC Pearson *r*.

#### Imputation baselines

To benchmark AG2 on bulk track imputation, we compared it against a retrained external baseline, Avocado, and two training-track baselines: the cross-condition channel mean and an oracle that averages the most correlated training tracks.

##### Mean training track baseline

For each test track, the mean baseline predicts the average signal across all training tracks available for that channel. This baseline captures assay-specific signal profiles and invariant genomic landmarks, but cannot capture condition-specific variation; for RNA-seq, because held-out tracks span distinct channels (such as total and poly(A)^+^ protocols), channel-specific training means yield non-zero LFC correlation without capturing condition-specific regulation.

##### Closest training tracks baseline

To establish an empirical upper bound for nearest-neighbor transfer, we evaluated a closest training track oracle. For each evaluated interval, we identified the training tracks of the same channel that had the highest Pearson correlation with the ground-truth test track, and averaged their signal. Because selecting the closest tracks uses the held-out target track itself, this method serves as an oracle upper bound for nearest-neighbor transfer. We averaged the five closest training tracks of the same channel (or all available tracks if fewer than five existed); for the single test channel lacking same-endedness training tracks (paired-end H3K79me2), the mean and closest baselines used single-end tracks.

##### Avocado multi-resolution tensor factorization

We trained the Avocado tensor factorization model^45^ across the six bulk modalities in the imputation split (ATAC-seq, DNase-seq, CAGE, RNA-seq, histone ChIP-seq, and TF ChIP-seq). Avocado models epigenomic tracks by learning latent representations for cell types, assays, and multi-scale genomic positions, and then passing concatenated embeddings through an MLP to predict continuous signal. Training followed the official two-stage protocol:

- **Hyperparameter sweep on pilot regions:** We conducted a 100-run random hyperparameter search on the ENCODE pilot regions (∼1% of the human genome) for 800 epochs, sweeping cell-type and assay latent dimensions (16 to 256), multi-scale genomic factors at 32 bp (5 to 25), 320 bp (10 to 50), and 6.4 kb (15 to 75), and MLP architectures (depth 0 to 4, width 128 to 2048).
- **Genome-wide fine-tuning and model selection:** The top 10 candidate models ranked by validation MSE were fine-tuned across all human autosomes (chromosomes 1–22) and sex chromosomes (X and Y). Genomic position embeddings were fine-tuned separately per chromosome for an extended 800 epochs (compared to the default 400 epochs). The final model was selected based on validation set performance across our primary biological evaluation metrics (support-weighted peak AUPRC).
- **Resolution and preprocessing:** To maintain native compatibility with our evaluation pipeline and track resolutions without lossy downsampling, we adapted Avocado from its original 25 bp resolution to 32 bp resolution. Consequently, the multi-scale genomic factors originally defined at 25 bp, 250 bp, and 5 kb were shifted to 32 bp, 320 bp, and 6.4 kb, respectively. High-resolution 1-bp tracks (ATAC-seq, DNase-seq, CAGE, RNA-seq) were downsampled to 32 bp via sum-pooling, and 128-bp tracks (histone ChIP-seq and TF ChIP-seq) were upsampled to 32 bp by bin repetition and dividing by the number of repeats. Input tracks were normalized using an arcsinh transformation prior to training and inverted with sinh at inference.

#### Primary PBMC enhancer annotation

We evaluated primary-cell H3K27ac imputation using the *all-folds* distilled imputation-split model (Imputation split), in which all bulk PBMC tracks were withheld from training so that the nine peripheral blood immune reference conditions (`UBERON:0000178` paired with `CL:0001054` CD14^+^ monocyte, `CL:0000236` B cell, `CL:0000787` memory B cell, `CL:0000788` naive B cell, `CL:0000895` naive CD4^+^ T cell, `CL:0000900` naive CD8^+^ T cell, `CL: 0000907` central memory CD8^+^ T cell, `CL:0000913` effector memory CD8^+^ T cell, and `CL:0000623` natural killer cell) were trained exclusively on scRNA-seq and scATAC-seq pseudobulks. Candidate elements were ENCODE GRCh38 representative DNase I hypersensitive sites (rDHSs; `ENCFF708KEK`) fully contained within genome-wide evaluation intervals. Following the ENCODE cCRE convention to capture flanking nucleosomes, each rDHS was expanded by 500 bp on each side when scoring histone ChIP-seq and scored unexpanded for scATAC-seq. Mean signal per element was log(*x* + 1)-transformed and *z*-scored across all genome-wide rDHSs independently for each cell type and assay.

To classify elements with enhancer-like signatures (Supplementary Fig. 7d), we restricted within each cell type to accessible (*z*_scATAC_ ≥ 1.67), non-promoter rDHSs (midpoint ≥200 bp from the nearest GENCODE v46 protein-coding or lncRNA TSS) and labeled elements positive if held-out ENCODE primary PBMC Mint-ChIP-seq H3K27ac fold-change-over-control signal^46^ satisfied *z*_H3K27ac_ ≥ 1.67. Within each cell type, we computed AUPRC (as average precision) comparing AG2-imputed H3K27ac (paired-end ENCODE histone ChIP-seq channel) against observed primary scATAC-seq (10x nuclear channel) and observed H3K27ac ChIP-seq from lineage-matched ENCODE cell lines (HL-60 for CD14^+^ monocytes; GM12878 for B, memory B, and naive B cells; and DND-41 for CD4^+^ T, CD8^+^ T, and NK subsets). To evaluate cell-type specificity across all genome-wide rDHSs (Supplementary Fig. 7e), each site’s mean *z*-score across the nine PBMC cell types was subtracted independently for imputed and observed Mint-ChIP H3K27ac, and Pearson *r* was computed across all pooled (rDHS, cell type) pairs.

### Enhancer–gene linking evaluations

#### Enhancer–gene scoring

We scored E2G pairs using input gradients of predicted gene expression, using the same procedure for the E2G benchmarks reported here and for genome-wide inference. For each gene, we centered a 1-Mb input interval on its median TSS and conditioned the model on the reference condition of interest. We then defined a scalar predicted expression readout *ŷ_g_* for the target gene *g* from one of two heads:

- **Bulk RNA-seq:** predicted base-resolution coverage averaged across observed channels on the gene’s strand, summed over annotated exonic positions of *g* (GENCODE v46) to give *ê_g_*, and transformed as *ŷ_g_* = log(*ê_g_* + 10*^−^*^6^).
- **Single-cell gene expression:** predicted scalar gene-level expression for *g* averaged across observed channels on the gene’s strand (*ê_g_*) and transformed as *ŷ_g_* = log(*ê_g_* + 10*^−^*^6^).

For both heads, we used only channels where the reference condition was observed during training (i.e., imputed channels were omitted).

We computed the gradient of *ŷ_g_* with respect to the one-hot input sequence and took, at each position, the gradient for the reference base minus the mean gradient across the four bases. The score for an enhancer was the mean of the absolute attribution over a 2400-bp window centered on the enhancer midpoint, weighted by a Gaussian kernel (*σ* = 300 bp) centered on the midpoint. Enhancers whose midpoint fell outside the input interval (more than 524,288 bp from the TSS) received a score of zero.

In contrast to previous work^14,16^, which normalized enhancer gradient scores by the mean absolute attribution across the entire input window, we omitted this window-wide normalization for all evaluated models, as we found that using unnormalized scores improved performance across all models (Supplementary Fig. 11a–c).

##### Expression-conditioned enhancer–gene scoring

To evaluate E2G linking without relying on predefined reference conditions, we also scored the astrocyte^56^ and ENCODE-rE2G K562^55^ CRISPRi benchmarks using expression conditioning (Stage 4: Post-training for expression conditioning). For each benchmark, we passed the single-cell RNA-seq profiles from the corresponding dataset through the cell encoder (Single-cell expression autoencoder) and averaged the resulting cell embeddings to form the mean cell embedding. Conditioning the post-trained AG2 model on this embedding, we defined the scalar readout *ŷ_g_* from the denoised single-cell gene expression head on the target gene’s strand and computed Gaussian-weighted input-gradient attribution scores identically to the reference-conditioned setting.

#### Enhancer–gene linking benchmarks

##### Held-out CRISPRi benchmark

We evaluated E2G linking on the held-out CRISPRi benchmark from ENCODE-rE2G^55^, which comprises element–gene pairs tested in K562, WTC11, Jurkat, GM12878 and HCT116 cells that do not overlap the ENCODE-rE2G training data. After updating outdated gene symbols to match GENCODE v46, all 4378 pairs (190 positives) were retained. By distance from the element to the gene TSS, there were 58 pairs (30 positives) within 10 kb, 670 (87) at 10–100 kb, 3600 (73) at 100–2500 kb and 50 (0) beyond 2.5 Mb. By cell line: K562, 1918 (118); WTC11, 1921 (15); HCT116, 396 (34); Jurkat, 75 (7); GM12878, 68 (16). Following the authors, each pair was weighted by the probability that it reflects a direct rather than indirect effect, as provided by the benchmark^55^.

##### K562 CRISPRi benchmark

We also evaluated E2G linking on the K562 CRISPRi benchmark assembled in the ENCODE-rE2G study^55^. Following the authors, we retained element–gene pairs with a defined regulatory label whose target gene belonged to the ENCODE-rE2G gene universe (10,356 pairs, 471 positives), and excluded three negative pairs whose target gene Ensembl identifiers were absent from GENCODE v46, yielding 10,353 element–gene pairs (471 positives). Using the element-to-TSS distances provided with the benchmark, there were 221 pairs (131 positives) within 10 kb of the TSS, 1597 pairs (257 positives) at 10–100 kb, and 8535 pairs (83 positives) at 100–2500 kb.

##### Astrocyte CRISPRi benchmark

We used the CRISPRi screen of candidate enhancers in cultured human astrocytes from ref.^56^. Following the authors’ criteria, positives were defined as element–gene pairs with a significant decrease in expression (FDR < 0.1 and a negative log fold change, termed “permissive hits” by the authors) whose element overlaps an ENCODE candidate regulatory element. Negatives were all other tested pairs whose element overlaps an ENCODE candidate regulatory element. This procedure yielded 1902 pairs, of which 122 were positives.

##### Scoring with AG2

All E2G benchmarks were scored using the *all-folds* distilled AG2 model. For each element– gene pair, AG2 was conditioned on the cell line or cell type in which the pair was tested. For the held-out CRISPRi benchmark, the model was conditioned on the corresponding cell line (K562, WTC11, Jurkat, GM12878 or HCT116), and *ŷ_g_* was computed from bulk RNA-seq predictions (between one track per gene for Jurkat and six for GM12878). For the astrocyte benchmark, the model was conditioned on the brain astrocyte reference condition (`CL:0000127`, `UBERON:0000955`), and *ŷ_g_* was computed from single-cell gene expression predictions. The single-cell RNA-seq training data for this condition was a pseudobulk of all 47,577 cells profiled in the screen itself^56^, so this output corresponded to the cells in which the element–gene pairs were tested.

To test whether AG2 can link enhancers to genes without any observed bulk data for the cell type, we also evaluated the model trained with the imputation split enabled (Imputation split; “AG2 (no bulk astrocytes)”). In this split, all human astrocyte bulk tracks (RNA-seq, CAGE, splice junctions, DNase-seq, and histone and TF ChIP-seq) were held out from training, while the brain astrocyte single-cell RNA-seq tracks from the screen were retained. The screen pseudobulk was therefore the only training data for this condition. This model was conditioned and scored identically to AG2 on the astrocyte benchmark.

##### Baselines

For AG1^14^ and Borzoi^16^, which are not conditioned, we applied the same scoring procedure to their bulk RNA-seq tracks whose ontology term matched the tested cell line or cell type, using the same track criteria and averaging as for AG2. For the held-out CRISPRi benchmark, this selected the same assays as for AG2: one to three tracks per gene for AG1, and one to 14 for Borzoi, which provides one track per ENCODE experiment.

Neither model has single-cell outputs, so for the astrocyte benchmark we used their ENCODE astrocyte RNA-seq tracks (`CL:0000127`). For Borzoi, we averaged the predictions of the four released replicate models and computed gradients of this ensemble prediction. Borzoi uses a 524,288-bp input window, so elements more than 262 kb from the TSS lie outside its input and were assigned a score of zero.

We also compared against three predictors that do not use a sequence model. For the held-out CRISPRi benchmark, we used the published ENCODE-rE2G predictions for the same pairs^55^. For the astrocyte benchmark, we used the EGrf predictions released with the dataset^56^. For both benchmarks, we also scored each pair by the inverse of its distance to the TSS. All methods were evaluated on the same set of pairs.

##### Metrics

We report AUPRC, computed as average precision. For the held-out CRISPRi benchmark, pairs were weighted by their direct-effect probability as described above; astrocyte pairs were unweighted. We computed AUPRC across all pairs and within TSS-distance bins (0–10 kb, 10–100 kb and 100–2500 kb), skipping bins without positives.

#### Expression-conditioned enhancer–gene linking in the Parkinson’s disease cohort

In the cingulate cortex of the Parkinson’s disease single-nucleus multiome cohort^49^, we compared E2G links predicted via expression conditioning against those predicted from matched brain reference conditions (Supplementary Fig. 11g). Predictions were generated natively on the T2T-CHM13v2.0 assembly and gene annotation without liftover to GRCh38 across protein-coding genes (*n* = 19,748). We evaluated the 21 cingulate cortex cell types with distinct dorsolateral prefrontal cortex (`UBERON:0009834`) reference condition proxies (Supplementary Table 14; L5 IT A and Lamp5 Exc were excluded because they shared reference proxies with L5 IT B and Lamp5, respectively, and immune and unclassified neuron clusters lacked reference proxies). Each cell type was scored under (i) expression conditioning on the mean 10x snRNA-seq cell embedding using the denoised single-cell gene expression head, and (ii) reference conditioning on its matched proxy using the standard single-cell gene expression head.

We evaluated candidate E2G links between the 19,898 caQTL-ASCA chromatin accessibility peaks (FDR ≤ 0.1) identified in the Parkinson’s disease study^49^ and protein-coding genes within the 1-Mb scoring window. Mapping each peak to all 500-bp tiles on our genome-wide scoring grid overlapping it by at least 250 (Genome-wide candidate tiling and scoring), we called a (peak, gene) pair active if it passed all three atlas thresholds on predicted target gene expression, local chromatin accessibility, and input-gradient attribution (Threshold selection). To quantify cell-type specificity and agreement across conditioning modes, we computed the pairwise Jaccard similarity between the sets of called (peak, gene) links for each of the 21 expression-conditioned cell types and each cell type’s reference condition proxy (Supplementary Fig. 11g, left), alongside the pairwise Jaccard similarity among the reference condition proxies themselves across the same 21 × 21 cell-type layout as a baseline of cross-cell-type similarity (Supplementary Fig. 11g, right).

### Complex trait evaluations

#### TraitGym evaluation

To evaluate zero-shot prioritization of non-coding variants implicated in complex human traits, we benchmarked AG2 on the TraitGym complex traits dataset^73^. This benchmark comprises 11,400 variants, contrasting fine-mapped GWAS causal variants (PIP > 0.9) against negative control variants matched for distance to the nearest TSS, minor allele frequency, and linkage disequilibrium (LD) score at a 1:9 positive-to-negative ratio.

##### AG2 variant scoring

We used precomputed variant scores across 500 prioritized primary single-cell reference conditions (a subset of the 504 reference conditions in GWAS panel), retaining only experimentally observed (non-imputed) tracks. For each of the eight profile modalities (1-bp bulk ATAC-seq, DNase-seq, CAGE, bulk RNA-seq, CpG methylation, and scATAC-seq; 128-bp histone and TF ChIP-seq), each observed track was scored as the ℓ_2_ norm of log(*ŷ*_ALT_ + 1) log(*ŷ*_REF_ + 1) across the 1-Mb window centered on the variant. For single-cell gene expression, each observed channel was scored as the ℓ_2_ norm of predicted gene-level variant scores Δ_expr_ across all annotated genes in the window. The final zero-shot score per variant was the maximum score across all observed tracks and all nine modalities.

##### Baselines and ensemble

We compared AG2 against three baselines: (i) AG1^14^, using its zero-shot maximum score across all output tracks spanning its six corresponding modalities (bulk ATAC-seq, CAGE, histone modification ChIP-seq, transcription factor ChIP-seq, DNase-seq, and bulk RNA-seq); (ii) CADD^129^ whole-genome PHRED-scaled pathogenicity scores; and (iii) GPN-Star^74^ scores (negated so that higher values indicate greater predicted causality). We also evaluated an equal-weight ensemble combining AG2 and GPN-Star (“AG2 + GPN-Star”), computed by min–max scaling each method’s scores across the benchmark variants to [0, 1] and averaging the two normalized scores.

We evaluated all methods across the unified variant set using AUPRC (computed as average precision).

### Precomputed variant and enhancer–gene linking scores

#### GWAS panel

We used the *all-folds* distilled AG2 model to generate an atlas of scores for 6,825,839 variants across 510 conditions (504 reference conditions and 6 conditions evaluated via Stage 4 expression conditioning).

##### Variant set

We scored the union of three sources of common and trait-associated SNVs. First, all gnomAD^72^ variants with a genome allele frequency strictly between 0.05 and 0.95 (6,127,336 variants). Second, all SNVs in Open Targets Platform^27^ release 26.03 GWAS credible sets that were fine-mapped with SuSiE (Open Targets confidence ≥3, i.e. SuSiE with in-sample or out-of-sample LD; loci fine-mapped with Probabilistic Identification of Causal SNPs (PICS) only were excluded), giving 1,879,681 variants. Third, all variants of the TraitGym complex-traits benchmark^73^ (11,400 variants; mitochondrial variants excluded). Indels were excluded, and every variant was verified to have a reference allele that matches the GRCh38 (hg38) reference sequence before scoring; no mismatches were found. The final deduplicated set comprises 6,825,839 unique SNVs on chromosomes 1–22 and X, each scored across 510 conditions. Of these 3,709 SNVs are unique to TraitGym (3,675 matched controls, 34 positives). The remaining 6,822,130 SNVs were used for GWAS analysis.

##### Scored conditions

We scored 510 primary human conditions: 504 reference conditions selected to maximize tissue and cell-type coverage across our single-cell corpus, plus six bone-marrow and endometrial cell states scored via Stage 4 expression conditioning using mean cell embeddings from the cell encoder (osteoblasts, preosteoblasts, osteoclasts, and megakaryocytes from CZ CELLxGENE Discover dataset `1a2e3350-28a8-4f49-b33c-5b67ceb001f6`, and decidualized and non-decidualized endometrial stromal cells from dataset `9ddea8d9-cc4c-420a-90f6-880996f808d4`^130^). The conditions are listed in Supplementary Table 15.

#### E2G atlas

We used the *all-folds* distilled AG2 model to generate an atlas of E2G linking predictions across 640 primary single-cell reference conditions, annotating all human protein-coding and lncRNA genes in GENCODE v46 (*G* = 39,323).

##### Genome-wide candidate tiling and scoring

For each biological condition and target gene, a 1-Mb input sequence window was centered on the median TSS, and contiguous 500-bp sliding-window tiles (with 250-bp step size) were scored across the ±524,288 bp interval. Candidate tiles were evaluated using a paired backward and forward pass over the 1-Mb reference sequence. First, following Enhancer–gene scoring, we computed the input gradient of the scalar log-transformed predicted expression readout *ŷ_g_* with respect to the one-hot reference sequence, using the single-cell gene expression head for reference conditions or the denoised single-cell gene expression head under expression conditioning. Each candidate tile was assigned its Gaussian-weighted mean absolute gradient score, stored alongside the model’s scalar reference gene expression prediction for gene *g*. Second, a complementary forward pass extracted predicted base-resolution scATAC-seq profiles across the same 1-Mb interval, scoring each tile by the maximum predicted base-resolution signal within the tile per channel, averaged across channels.

##### Threshold selection

Candidate E2G links were filtered through a three-stage gating pipeline requiring thresholds on (i) predicted target gene expression, (ii) predicted local chromatin accessibility, and (iii) input-gradient attribution, calibrated against CRISPRi perturbation benchmarks in primary human astrocytes (UBERON:0000955_CL: 0000127) and K562 cells (EFO:0002067)^55,56^:

- **Target gene expression gate (***ŷ_g_* ≥**0.5):** To ensure that links were called only for transcriptionally active target genes in a given condition, we required a threshold on the scalar predicted log-transformed gene expression *ŷ_g_* (averaged across observed channels on the gene’s strand before log transformation; Enhancer– gene scoring). A threshold of *ŷ_g_* ≥0.5 retained nearly all CRISPRi-validated target genes in both astrocytes (97.7%) and K562 (96.7%), while excluding 69.7% and 68.3% of all GENCODE v46 genes in astrocytes and K562, respectively, consistent with the expectation that only a subset of genes is expressed in any given cell type.
- **Local chromatin accessibility gate (scATAC ≥1.25):** To restrict candidate tiles to accessible regions in the target condition, we required a predicted local scATAC-seq score ≥1.25, retaining 100% of CRISPRi-validated positive enhancers in astrocytes and 98.9% in K562 while excluding 98.4% and 93.4% of distal inaccessible tiles (>2 kb from both the target gene TSS and any ENCODE ATAC-seq narrowPeak), respectively.
- **Gradient attribution threshold (score ≥0.0012):** To call active E2G links among accessible candidate tiles of expressed genes, we required a threshold on the Gaussian-weighted input-gradient attribution score. A threshold of 0.0012 maximized the *F*_1_ score in the astrocytes benchmark (*F*_1_ = 0.65, precision = 0.69, recall = 0.61), while being within the near-optimal plateau for K562 (*F*_1_ = 0.74, precision = 0.71, recall = 0.77).

##### Summit-based tile merging and summary statistics

To define active E2G links and compare them across biological conditions, we applied a three-step consolidation procedure:

1. **Per-gene summit tile extraction:** Candidate tiles passing all three thresholds were designated as active tiles. Within each biological condition and target gene, we grouped strictly overlapping active tiles and only retained the tiles with the maximum gradient attribution score within each group (summit tiles); the remaining tiles were discarded.
2. **Within-condition enhancer merging:** Within each biological condition, overlapping 500-bp per-gene summit tiles were merged across all regulated target genes into mutually disjoint enhancer intervals, and every target gene whose summit tile fell within a merged interval was linked to that shared enhancer locus.
3. **Cross-condition consensus reconciliation:** To compute statistics across the 640 biological conditions, per-condition enhancer intervals linked to the same target gene were reconciled into cross-condition consensus enhancer intervals via greedy clustering at a 50% mutual reciprocal overlap threshold (overlap (*A*, *B*) ≥ 0.5 × max(|*A*|, |*B*|). Intervals were processed in decreasing order of the number of conditions in which that E2G link was observed. Each interval was either retained as a new cluster representative or assigned to the previously retained interval with which it had the largest reciprocal overlap; the consensus interval of each cluster was the coordinates of its representative interval rather than the union of its members.

Promoter-proximal and distal tiles were processed jointly in all three steps. Whereas the 1-Mb scoring window was centered on each gene’s median TSS across transcripts, TSS distances and promoter-proximal (≤500 bp) versus distal (>500 bp) labels were measured from the enhancer interval midpoint to the gene’s representative TSS in GENCODE v46 (the start of its MANE Select transcript^131^, or its Ensembl canonical transcript if absent). Because classification was performed per link, a shared enhancer interval can be promoter-proximal to one gene and distal to another. Unless stated otherwise, reported summary statistics refer to distal E2G links.

Per-condition yields (Supplementary Fig. 13d–f; Supplementary Table 16) and pooled link distributions (Supplementary Fig. 13a–c,g) were both computed directly on the within-condition merged enhancer intervals from step 2 (prior to step 3 cross-condition reconciliation). For each condition, we report the number of predicted active distal enhancer intervals (Supplementary Fig. 13d), distal E2G links (Supplementary Fig. 13e), and target genes with at least one distal link (Supplementary Fig. 13f). Pooling step 2 intervals across all 640 conditions, we also report distributions of (i) enhancer-to-TSS distance, measured from the enhancer interval midpoint to the linked gene’s representative TSS (Supplementary Fig. 13a); (ii) distinct target genes linked per (condition, enhancer interval) (Supplementary Fig. 13b); (iii) distinct distal enhancer intervals linked per regulated (condition, gene) pair (Supplementary Fig. 13c); and (iv) within-condition merged enhancer interval length (Supplementary Fig. 13g). The cross-condition consensus intervals from step 3 were used only for consensus enhancer interval lengths (Supplementary Fig. 13h) and cross-condition specificity analyses, defined by the number of conditions in which each consensus E2G link was active (Fig. 4f,g).

##### Locus analysis at *BCL11A*

For *BCL11A* (`ENSG00000119866`; MANE Select TSS at `chr2:60553654`), we retrieved within-condition merged active E2G links (promoter-proximal and distal) across 13 reference conditions (Supplementary Table 16): dorsolateral prefrontal cortex medial ganglionic eminence interneuron, bone marrow (hematopoietic stem cell, megakaryocyte–erythroid progenitor, proerythroblast, erythroblast, orthochromatic erythroblast, and granulocyte–monocyte progenitor), interventricular septum mast cell, liver neutrophil, lung monocyte, mucosa dendritic cell, thymus pro-B cell, and blood naive B cell. Links were plotted in 500-bp midpoint bins (maximum gradient score per bin; Fig. 4h) alongside GRCh38 reference intervals (converted from GRCh37 via UCSC liftOver where reported on hg19) for the erythroid +62 kb (`chr2:60490400-60491800`), +58 kb core (`chr2:60495049-60495539`, intersection of Bauer *et al.* and Javierre *et al.*), and +55 kb (`chr2:60497800-60499200`) enhancers^57,58,60^, and promoter capture Hi-C HindIII fragments in naive and total B cells (+200 kb, `chr2:60352101-60353675`^58^) and dorsolateral prefrontal cortex (+479 kb, `chr2:60071284-60077096`^59^). For Fig. 4i, bone marrow erythroblast base-resolution scATAC-seq predictions (10x nuclear channel, 128-bp centered moving average) and active E2G links were paired with wild-type and `sg1617`-edited HUDEP-2 data^60^: unsmoothed 20-bp ATAC-seq bigWig tracks (`GSE292669`; mean of two coverage-normalized replicates) and enhancer-probe Capture-Pore-C contacts (`GSE292694`), quantified per baited enhancer (+55, +58, and +62 kb) as uniquely mapped paired-end tags (PETs) joining distinct NlaIII fragments overlapping the enhancer and promoter (TSS 1 kb) per million genome-wide PETs (mean of two replicates, unthresholded). In a 1-Mb window centered on `chr2:60495266` (first base of the GATAA motif in the `sg1617` gRNA target, `chr2:60495263-60495284`^61,66^), we computed saturated ISM across `chr2:60495262-60495285` in bone marrow erythroblasts using the 1201-bp single-cell chromatin accessibility center-mask scorer (Variant scoring) on the reference and `chr2:60495266:G> A` sequences (Fig. 4j), and plotted unsmoothed base-resolution reference and alternate scATAC-seq predictions across the +58 kb core enhancer in bone marrow erythroblasts and blood naive B cells (Fig. 4k).

### GWAS analysis

#### GWAS fine-mapping data and trait definitions

We obtained fine-mapped GWAS credible sets and study metadata from the Open Targets Platform (release 26.03)^27^. We used credible sets fine-mapped with SuSiE (with an in-sample or out-of-sample LD reference), excluding credible sets derived with PICS. We grouped GWAS studies into 28 traits by curating their reported trait names (Supplementary Table 17). Two of these traits pooled related diseases: autoimmune diseases (inflammatory bowel disease, rheumatoid arthritis, psoriasis and psoriatic arthritis, systemic lupus erythematosus, type 1 diabetes, autoimmune thyroid disease, ankylosing spondylitis, multiple sclerosis, celiac disease, myasthenia gravis, Sjögren’s syndrome and sarcoidosis) and asthma/allergy (asthma, eczema, urticaria and allergic rhinitis). For each trait, we pooled the credible sets of all its studies and assigned each variant its maximum PIP across them. All analyses used SNVs only. As a background set, we used common SNVs (gnomAD^72^ v4.1 genome allele frequency between 0.05 and 0.95) that were not in any of the SuSiE GWAS credible sets. The combined set of variants corresponds to the GWAS Panel, excluding the variants unique to Traitgym.

#### Definition of called variants

For the GWAS analysis presented in this paper, we restricted our attention to AG2 single-cell gene expression variant effect predictions, as these can be directly used to nominate genes and cell types. Specifically, for each variant, nearby protein-coding gene (GENCODE v46^132^) and condition, we recorded the predicted log fold change in single-cell gene expression (Variant scoring), as well as the gene’s predicted expression level for whichever allele was more highly expressed. We say AG2 *called* a variant–gene pair in a condition if, in at least one of the condition’s single-cell gene expression output channels, the absolute predicted log fold change Δ_expr_ exceeded 0.1 and the gene was predicted to be expressed (log(predicted CPM + 1) > 1.5). When several channels qualified, we kept the one with the largest absolute predicted log fold change. For the six conditions scored using expression conditioning, a variant–gene pair was also called if the predicted variant effect from AG2’s denoised expression output passed the same thresholds.

Of the 6,822,130 variants analyzed, 134,813 were called for at least one gene in at least one condition at the selected thresholds.

#### TenK10K threshold sweep

We calibrated the two call thresholds (Definition of called variants) using sc-eQTLs from the TenK10K study^7^. We used its SuSiE credible sets for protein-coding genes (GENCODE v46^132^), restricted to SNVs, and its nominal association *P* values for all variant–gene pairs tested in each cell type. We matched each TenK10K cell type to the AG2 reference condition of the same cell type (Supplementary Table 9). We defined the following variants and (variant, gene, condition) triplets:

- **Fine-mapped eQTLs:** SNVs with PIP > 0.9 in at least one credible set (6362 SNVs). Each such credible set contributed an on-target triplet: the SNV, the gene of the credible set and the reference condition matched to its cell type (14,010 distinct triplets).
- **Off-target triplets:** a fine-mapped eQTL paired with a protein-coding gene and cell type for which it was tested but not associated (nominal *P* > 0.05) and not in a credible set. Triplets that were also on-target, which can occur when two cell types share a reference condition, were excluded.
- **Background variants:** tested SNVs with a minor allele frequency >0.01 that had nominal *P* > 0.05 for every gene and cell type tested and were not in any credible set (1,235,667 SNVs).

All other SNVs, including credible-set SNVs with PIP ≤ 0.9, were excluded.

We scored these variants using the AG2 single-cell gene expression output and performed a sweep over the thresholds we used to define a call. Specifically, we varied the log fold change threshold from 0.05 to 0.20 in steps of 0.01, and the expression threshold over 0 (no expression requirement), 0.5, 1, 1.5, 1.75, 2 and 2.5. For each of the 112 threshold combinations, we computed three quantities:

- **Recall:** the fraction of fine-mapped eQTLs called in at least one of their on-target triplets.
- **Enrichment for fine-mapped eQTLs:** the recall divided by the fraction of background variants called for any gene in any of the 26 reference conditions. This comparison is conservative, because fine-mapped eQTLs counted only if called on target.
- **On-target enrichment:** the fraction of on-target triplets called divided by the fraction of off-target triplets called.

#### Cell-type specificity of calls

To relate TSS distance to cell-type specificity without relying on tissue or dataset labels, we grouped reference conditions by transcriptional identity. Starting from the CPM matrix (averaged over channels where relevant) of the 1943 reference conditions in the AG2 single-cell expression pre-training corpus, we retained protein-coding genes with mean expression 1 in at least 20 conditions, applied a log(CPM + 1) transform, and selected the 2000 genes with the highest standard deviation across conditions. Each gene was scaled to unit variance without centering, flooring standard deviations at their median.

We factorized this matrix using non-negative matrix factorization (NMF; scikit-learn^127^; 40 programs, NNDSVDa initialization, Frobenius loss), weighting each condition’s program usages by the ℓ_2_ norm of the program’s gene loadings and normalizing them to sum to one. We excluded eight programs manually judged to reflect technical factors rather than cell-type identity and renormalized over the remaining 32 programs.

We clustered the 504 GWAS-panel reference conditions by Ward’s hierarchical clustering on the square root of their program usages, cutting the tree into 60 cell-type clusters (grouping the same cell type across tissues and datasets; median size, 8 conditions) nested within 12 lineage clusters. Each condition scored with expression conditioning formed a singleton cell-type cluster within the lineage cluster of a related reference condition: decidual stromal cells for both endometrial stromal cell states, tracheal chondrocytes for osteoblasts and preosteoblasts, splenic macrophages for osteoclasts, and bone-marrow erythroblasts for megakaryocytes.

A call in a given condition was considered corroborated if the condition was scored using expression conditioning or if another condition in its cell-type cluster showed a same-sign effect at relaxed thresholds (|log fold change| > 0.05 and log(predicted CPM + 1) > 1, in any track). Call breadth was defined as the number of lineage clusters containing at least one corroborated called condition (or any called condition, if none were corroborated), yielding four specificity categories:

- **Single condition:** called in 1 condition.
- **Lineage-restricted:** called in 2 conditions, breadth 1.
- **Multi-lineage:** breadth 2–3.
- **Broad:** breadth ≥ 4.

Call-level analyses (variant-to-gene TSS distance) used each call’s own category, whereas variant-level analyses (e.g., ENCODE cCRE overlap) assigned each called variant the broadest category among its calls. Here and below, a gene’s TSS was taken from its MANE Select transcript^131^ in GENCODE v46, or from its Ensembl canonical transcript for the 839 protein-coding genes lacking one.

#### Conservation of called variants

We annotated each variant with its distance to the nearest protein-coding TSS, in four bins: <1 kb, 1–10 kb, 10–100 kb and >100 kb. Baseline conservation falls steeply with distance to the TSS, so we compared called with uncalled variants within each distance bin. We extracted phyloP scores at each variant’s position from the Zoonomia 241-mammal alignment^133^. A position was conserved if phyloP ≥ 2.27, the threshold for evolutionarily constrained bases at a 5% false discovery rate in the Zoonomia alignment^133^. Variants without a phyloP score were excluded.

#### Overlap of called variants with ENCODE cCREs

We intersected variants with the ENCODE registry of cCREs, version 4^40^. A variant was in a cCRE if its position lay within any cCRE. We stratified called variants by their variant-level specificity category. For each category and distance bin, we computed the fraction of called variants in a cCRE, with 95% CIs from the block bootstrap, and compared it with the fraction of uncalled variants in a cCRE in the same distance bin.

#### Cell-type enrichment of GWAS credible sets

For each trait and condition, we tested whether the trait’s credible-set SNVs were called in the condition more often than background variants. Let *a* be the number of the trait’s credible-set SNVs called (for any gene) in the condition, and *b* the corresponding number of background variants. Let *N*_CS_ and *N*_BG_ be the total numbers of credible-set SNVs of the trait and of background variants. We applied a one-sided Fisher’s exact test for enrichment to the 2 × 2 table [[*a*, *b*], [*N*CS − *a*, *N*BG − *b*]] and corrected for the number of conditions tested per trait with the Benjamini–Hochberg procedure. We refer to the corrected *P* value as the enrichment FDR. For each trait, the condition with the lowest enrichment FDR was its trait-matched condition.

Seventeen traits had an enrichment FDR ≤10*^−^*^7^ in at least one condition: Alzheimer’s disease, asthma/allergy, atrial fibrillation, autoimmune diseases, blood pressure, cholesterol, coronary artery disease, glomerular filtration rate, height, lung capacity, platelet count/volume, prostate cancer, red blood cell count/volume, serum globulin, type 2 diabetes, waist-to-hip ratio and white blood cell count. These traits and their trait-matched condition were used in all subsequent analyses. The ontologies of the trait-matched conditions are given in (Supplementary Table 18).

To construct the trait-by-cell-type heatmap, we transformed the enrichment FDRs of these 17 traits to -log_10_(FDR) and *z*-scored each trait across all conditions. For each trait, we took the five conditions with the lowest enrichment FDR (among those with FDR < 0.05) and displayed the union of these conditions across traits. Traits and conditions were ordered by hierarchical clustering of the displayed *z*-scores (average linkage, Euclidean distance). Column annotations show broad lineages, assigned by matching condition names against curated lists of cell-type keywords.

#### PIP enrichment

For each of the 17 traits, we considered all of its credible-set SNVs, with one entry per trait–SNV pair carrying the SNV’s maximum PIP across the trait’s credible sets. We binned the pairs by PIP: 0–0.01, 0.01–0.05, 0.05–0.1, 0.1–0.25, 0.25–0.5, 0.5–0.75 and 0.75–1 (intervals closed on the right). We used two definitions of a called trait–SNV pair. Under the first, the SNV was called (for any gene) in the trait’s trait-matched condition. Under the second, used for comparison, the SNV was called in any of the 510 conditions. For each definition, we pooled pairs across traits and computed the call rate (the fraction of called pairs) in each PIP bin relative to the call rate in the lowest bin (PIP ≤ 0.01), with 95% CIs from the block bootstrap.

### PIP enrichment stratified by eQTL colocalization

This analysis used the same trait–SNV pairs, trait-matched conditions, PIP bins and fold definition as the PIP enrichment analysis. We split the trait–SNV pairs into two groups according to whether any of the trait’s credible sets containing the SNV colocalized with an eQTL or sc-eQTL credible set, using the colocalization results of the Open Targets Platform (release 26.03, the same release as the GWAS data)^27^. Colocalization was defined as a COLOC^134^ posterior probability of a shared causal variant (*H*_4_)≥0.8, for any gene, tissue or cell type. Folds were computed separately within each group, relative to the call rate at PIP 0.01 in the same group, with 95% Cis from the block bootstrap.

### Direction-of-effect concordance

If a gene affects a trait through its expression level, the ALT allele’s effect on the trait should equal the direction of the predicted expression change multiplied by the effect of increased gene dosage. We approximated the effect of increased dosage as the opposite of the effect of loss of function (LoF), as estimated by rare-variant gene burden tests. We obtained the burden test results from the Open Targets Platform (release 26.03, the same release as the GWAS data)^27^.

We tested this prediction for every trait–gene pair in which the gene was called at one of the trait’s credible-set SNVs in the trait-matched condition, counting each pair once. Each pair was represented by the highest-PIP SNV among those at which the gene was called (PIP: the maximum across the trait’s credible sets; ties broken by the largest absolute predicted log fold change) and by the sign of that SNV’s AG2 score. Directions were then defined as follows:

- **GWAS direction:** the sign of the SNV’s ALT-allele effect size in the trait’s credible sets that contained it. Each credible set contributed under every ontology term of its study, and for each term we took the majority sign across credible sets (ties were discarded).
- **LoF direction:** the sign of the effect estimate (or the log odds ratio for binary traits) of the most significant Open Targets gene burden result for the same gene and term. Burden results for synonymous-variant masks were excluded, because these masks serve as negative controls. A positive sign means that LoF increases the trait value or disease risk.

Terms were matched by name, so that the same trait under different ontologies (EFO and the Ontology of Biological Attributes) matched. Medication-use, sex-interaction and Gene Ontology terms were excluded. A term was concordant if sign(GWAS effect) = − sign(AG2 score) × sign(LoF effect).

Each trait–gene pair was classified as concordant or discordant by majority over its matched terms, that is, terms with both a GWAS and a LoF direction. Pairs without a matched term, as well as ties, were excluded. We computed the fraction of concordant pairs among concordant and discordant pairs and tested it against 0.5 with a two-sided exact binomial test.

We repeated the test on the pairs whose gene was not the nearest gene to the pair’s SNV, that is, pairs in which another protein-coding gene had a TSS strictly closer to the SNV than the gene’s own TSS. Ties, which occur for example when a readthrough gene shares the TSS of its upstream gene, were counted as nearest.

#### Locus-level and metacell variant interpretation

Individual GWAS loci (Fig. 5a,b,l; Supplementary Fig. 15a,b) were scored with the *all-folds* distilled AG2 model using the single-cell gene expression variant scorer (Variant scoring; observed channel with the largest absolute predicted log fold change retained when multiple channels applied):

- ***ETS2* (21q22):** All seven SNVs in the inflammatory bowel disease credible set of Stankey *et al.*^67^ containing rs2836882 (`chr21:39094644:G>A`) were scored on nearby protein-coding genes across seven primary ileum immune reference conditions (Fig. 5a,b, Supplementary Table 19).
- ***SCUBE3* (6p21.31):** The two SNVs in the standing-height credible set containing rs2038740 (`chr6: 35146765:C>T`) were scored in adult bronchus chondrocytes (`UBERON:0002185`, `CL:0000138`). Saturated ISM of predicted *SCUBE3* expression (10x 5’ whole-cell channel) was computed on the alternate (rs2038740-T) sequence across a 64-bp window and displayed across the central 42 bp (`chr6:35146733-35146775`; Fig. 5l).
- ***HERC2*–*OCA2* (15q13.1):** In primary skin melanocytes (`UBERON:0002097`, `CL:0000148`), we scored rs12913832 (`chr15:28120472:A>G`), rs6497271 (`chr15:28120285:A>G`), the 26 remaining SNVs in the adjacent melanoma credible set^135^, and the double-alternate haplotype carrying both rs12913832-G and rs6497271-G relative to the reference sequence (Supplementary Fig. 15a).
- ***SMIM1* (1p36.32):** The two reticulocyte-count credible-set variants rs6673829 (`chr1:3774926:G>A`) and rs1175550 (`chr1:3774964:A>G`)^27,136^ were scored on *SMIM1* across 21 primary bone-marrow hematopoi-etic reference conditions (Supplementary Fig. 15b).

To resolve variant effects across developmental and tissue single-cell atlases (Fig. 5m, Supplementary Fig. 14d, Supplementary Fig. 15c,d), raw UMI count matrices from CZ CELLxGENE Discover were randomly subsampled without replacement within each cell type to at most 10,000 cells (seed 0, excluding zero-count cells) and mapped to the cell encoder’s GENCODE v46 Ensembl vocabulary (explicitly stored zeros removed). All retained cells were passed through the cell encoder (unmasked *all-folds* model, *z* R^256^) and partitioned within each cell type into *k* = max(1, round(*n*/64)) cell-type-pure metacells by *k*-means clustering (`random_state` = 1, `n_` `init` = 10). Each metacell was scored with the *all-folds* post-trained AG2 denoised single-cell gene expression scorer (Stage 4: Post-training for expression conditioning), conditioned on the mean cell embedding of its member cells (for metacells exceeding 256 cells, the 256 cells nearest the metacell mean cell embedding in Euclidean distance were averaged). Two-dimensional UMAP coordinates (umap-learn^50^, `min_dist` = 0.5, `random_state` = 42) were fitted on single-cell embeddings, with metacell mean cell embeddings projected via `transform`. This protocol was applied to: (i) human embryonic limb lateral plate mesoderm (5–9 post-conception weeks; CELLxGENE `f9685456-c8a8-43ad-a326-476273ef36a0`; 100,935 cells across 34 cell types, 1575 metacells)^68^ for rs2038740 on *SCUBE3*, alongside metacell pseudobulk log(CPM + 1) expression of *SCUBE3* and *SOX9* (Fig. 5m, Supplementary Fig. 14d); (ii) normal-donor human distal airways (CELLxGENE `b61a921b-7fa3-4b42-b455-aaaf32447920`; 69,830 cells across 48 author clusters grouped into 19 cell types, 1088 metacells)^137^ for rs35705950 (`chr11:1219991:G>T`) on *MUC5B* (Supplementary Fig. 15c); and (iii) human endometrium (CELLxGENE `9ddea8d9-cc4c-420a-90f6-880996f808d4`; 75,844 cells across 16 cell types, 1183 metacells)^130^ for rs3820282 (`chr1:22141722:C>T`) on *WNT4* (Supplementary Fig. 15d).

#### Statistics

Unless stated otherwise, 95% CIs of rates and fold enrichments came from a genomic block bootstrap: the genome was divided into 1-Mb blocks, and each of the 2000 replicates resampled the blocks with replacement and kept all entries (variants or trait–SNV pairs) of each drawn block, across groups and traits. CIs are the 2.5th and 97.5th percentiles of the replicate statistic. This approach accounts for correlation between nearby variants (for example, through LD or shared regulatory elements) and for loci shared by related traits. Fold enrichments are ratios of rates pooled across traits. Fisher’s exact and binomial tests were computed with SciPy^138^.

## Acknowledgements

We thank Gabriella Stanton and Charlie Taylor for project management; Raina Thomas and Lucas Tenório for engineering contributions; Antonio García (Bio-Graphics) for figure design and scientific illustrations; and Elucidata for preparing and reprocessing datasets. We are grateful to Joseph Powell and Angli Xue for help with the TenK10K dataset, Trygve Bakken and Nelson Johansen for guidance on brain atlas data, Junyue Cao and Ziyu Lu for providing fragment files for mouse atlases, and Jacob Hepkema and Leopold Parts for providing processed HEK293T ATAC data. We also thank Irina Voineagu for discussions on astrocyte enhancer–gene data, Souradeep Basu and Michal Zielinski for early discussions on conditioning AlphaGenome, Jeff Cao and Ioannis Sarropoulos for feedback on model performance, Sarah Teichmann for pointing us to the *ETS2* example, and Amanda Stafford and Eliseo Papa for feedback on the manuscript. Finally, we acknowledge the study participants, tissue donors, and their families, and thank the contributing laboratories and public data initiatives for generating and sharing the datasets that made this work possible, including CZ CELLxGENE, the Human Cell Atlas, ENCODE, GTEx, 4D Nucleome, FANTOM5, TenK10K, EMBL-EBI, and the Open Targets Platform. This work was funded by Google DeepMind.

## Author contributions

A.G. and Ž.A. conceptualized the study with input from A.M., A. Karollus, J.C., P.K., and D. Hassabis. A.G. led project research and A.M. led engineering, with G.N. and A. Kosiorek leading model development, P.D. leading development of the model server, A. Karollus leading model evaluations, L.M.-P. and E.A. leading enhancer–gene linking, and J.R. leading large-scale inference. A.G., A.M., and Ž.A. conceived the conditional model architecture. G.N., S.D., A. Kosiorek, and V.D. developed and improved the model with input from A.G., A. Karollus, L.M.-P., and A.M. V.D. developed the gene expression head with contributions from G.N., and A.G. developed the methylation head with contributions from G.N. and input from N.L. J.C. advised on splicing and annotations. A. Kosiorek conceptualized and led development of sequence packing with input from G.N. A.G. conceived expression conditioning with input from P.D., A. Kosiorek, A.M., and Ž.A. P.D. and G.N. developed the single-cell expression autoencoder with contributions from A.M., L.M.-P., E.A., A.G., and A. Kosiorek; G.N. developed distillation with contributions from A. Kosiorek and S.D.; and A.G., A.M., and P.D. developed the post-training procedure with input from G.N., E.A., S.D., and A. Kosiorek. L.M.-P., A.G., A.M., and A. Karollus curated, processed, and prepared the training data with contributions from N.L. P.D., A.M., K.T., and V.D. designed and implemented the data pipelines for model training and evaluation with contributions from G.N. and L.N. A. Kosiorek developed the evaluation framework for model selection. P.D. and T.W. developed the model server with contributions from J.R., K.T., A.M., and G.N. P.D. developed the model API with contributions from A.M. and T.W. and input from D. Hariharan. J.R. led the development of infrastructure for large-scale inference with contributions from E.A. and A.M. and input from T.W. and K.T. T.W. advised on engineering pipelines and model serving. G.N. and V.D. developed the AlphaGenome 1 baseline with contributions from A. Karollus and L.M.-P. A. Karollus, J.R., A.M., and L.M.-P. developed the variant scoring methodology. A. Karollus led the development of model evaluations and biological case studies with contributions from L.M.-P., A.G., and A.M. Specific evaluation areas were covered as follows: A. Karollus covered condition-specific interval, ablation, and QTL evaluations and loci with contributions from L.M.-P., A.G., and A.M.; P.D. covered cross-modality imputation evaluations with input from L.M.-P., A.M., A. Karollus, and A.G.; A.M., A.G., and L.M.-P. covered expression-conditioning generalization, SARS-CoV-2 vaccination, and Parkinson’s disease cohort analyses with contributions from E.A.; L.M.-P. and E.A. covered enhancer–gene linking methodology and evaluations with contributions from J.R.; M.P. covered enhancer–gene linking locus analyses with contributions from J.P. and input from A.G.; and A. Karollus developed the GWAS analysis methodology and covered TraitGym and complex-trait variant analyses with contributions from A.M. and J.P. and input from C.B. A.G., A. Karollus, L.M.-P., E.A., M.P., A.M., P.D., and J.P. analyzed data and prepared figures. A.G., A. Karollus, J.P., Ž.A., L.M.-P., E.A., M.P., A.M., P.D., and N.L. wrote and edited the manuscript. L.H.W. managed and coordinated the project. A.G., P.K., and Ž.A. supervised the project. All authors reviewed the manuscript.

## Data availability

No new wet-lab experiments were conducted for this study. Bulk and single-cell functional genomics datasets used to pre-train, post-train, and evaluate AlphaGenome 2 were retrieved from public repositories and consortium portals, including ENCODE, the GTEx Portal, the 4D Nucleome Data Portal, FANTOM5, recount3, CZ CELLxGENE Discover (16 March 2026 snapshot for single-cell expression autoencoder training and post-training), the Gene Expression Omnibus (GEO), ArrayExpress, Zenodo, TenK10K, the eQTL Catalogue, TraitGym, the Open Targets Platform (release 26.03), and gnomAD (v4.1). Per-dataset study citations, accession identifiers, collection URLs, processing provenance, and ontology CURIE mappings are cataloged in Supplementary Tables 1, 2, and 3 (with benchmark cell-type and trait mappings in Supplementary Table 8, Supplementary Table 9 and Supplementary Table 17). Human analyses used the GRCh38 reference assembly with GENCODE v46 annotations (and T2T-CHM13v2.0 where noted in Methods), and mouse analyses used GRCm38 with GENCODE vM23 annotations. A filtered version of the precomputed AlphaGenome 2 predictions generated in this work, comprising the enhancer– gene linking atlas across 640 primary human conditions (Supplementary Table 16) and single-cell expression variant effect scores for trait-associated and common variants across 510 human conditions (Supplementary Table 15), is available at https://deepmind.google.com/science/alphagenome/downloads. Filtering was done on the basis of effect sizes for variant scores and for enhancer-gene links we used the same cutoffs described in the Methods.

## Code availability

AlphaGenome 2 is available for non-commercial use via an API at https://github.com/google-deepmind/alphagenome, which includes Python software to interact with the model. AlphaGenome2 model weights, code, and precomputed predictions are intended for research purposes only. They contain predictions with varying levels of confidence and caution should be exercised in their use. The information provided is not intended to be a substitute for professional medical advice, diagnosis, or treatment, and does not constitute medical or other professional advice. AlphaGenome2 and its outputs have not been validated for, and are not approved for, any clinical or diagnostic use. The model source code and weights will be provided upon final publication.

## AI use declaration

Large language models including Gemini 4 Argon, accessed with Google Antigravity (an agentic development environment) were used throughout this project to brainstorm, critique and implement model analyses and evaluation designs. Gemini was also used to draft and edit manuscript prose and visualization code. In each of these steps, the authors steered the agents and ran several rounds of adversarial review. Every AI-proposed hypothesis, analysis script, quantitative result, figure, and manuscript passage was inspected, verified against primary data, and revised by the authors, and all bibliographic citations were checked against original sources. The authors take full responsibility for all analyses, claims, and content in this manuscript.

## Supplementary Information

### Supplementary Figures

**Supplementary Figure 1.**
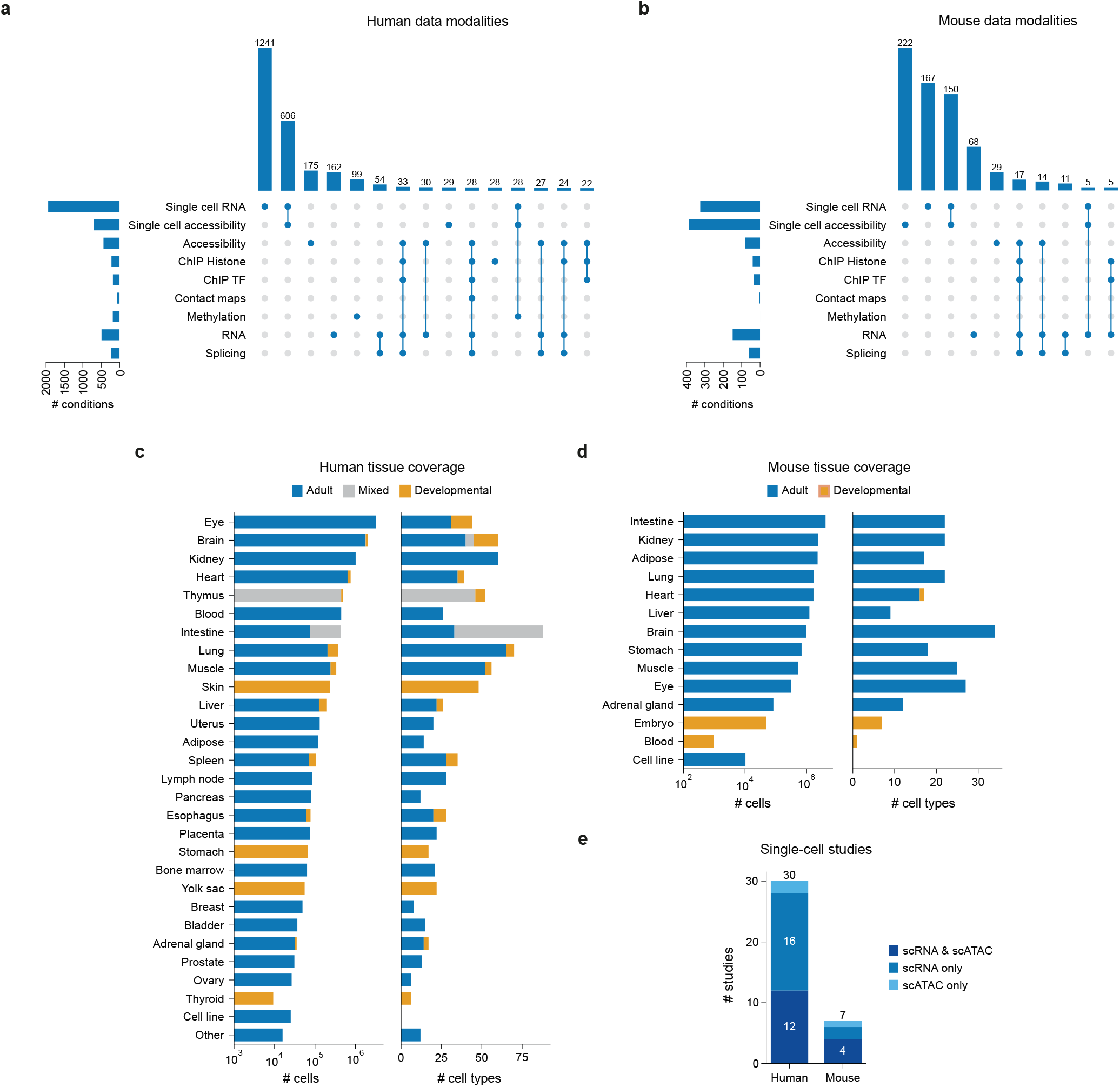
Overview of the AlphaGenome 2 training data corpus. **(a,b)** Modality coverage across conditions for human (**a**) and mouse (**b**). Horizontal bars (left) give the total number of conditions with observed data for each modality; vertical bars (top) give the number of conditions observed with each combination of modalities, indicated by connected dots below, for the most frequent combinations. **(c,d)** Tissue coverage of the single-cell data for human (**c**) and mouse (**d**). Left, number of cells per tissue (log scale); right, number of annotated cell types per tissue. Bars are colored by developmental stage of the source samples (adult, developmental or mixed). **(e)** Number of single-cell studies per organism, split by whether a study contributed both scRNA-seq and scATAC-seq, scRNA-seq only or scATAC-seq only.

**Supplementary Figure 2.**
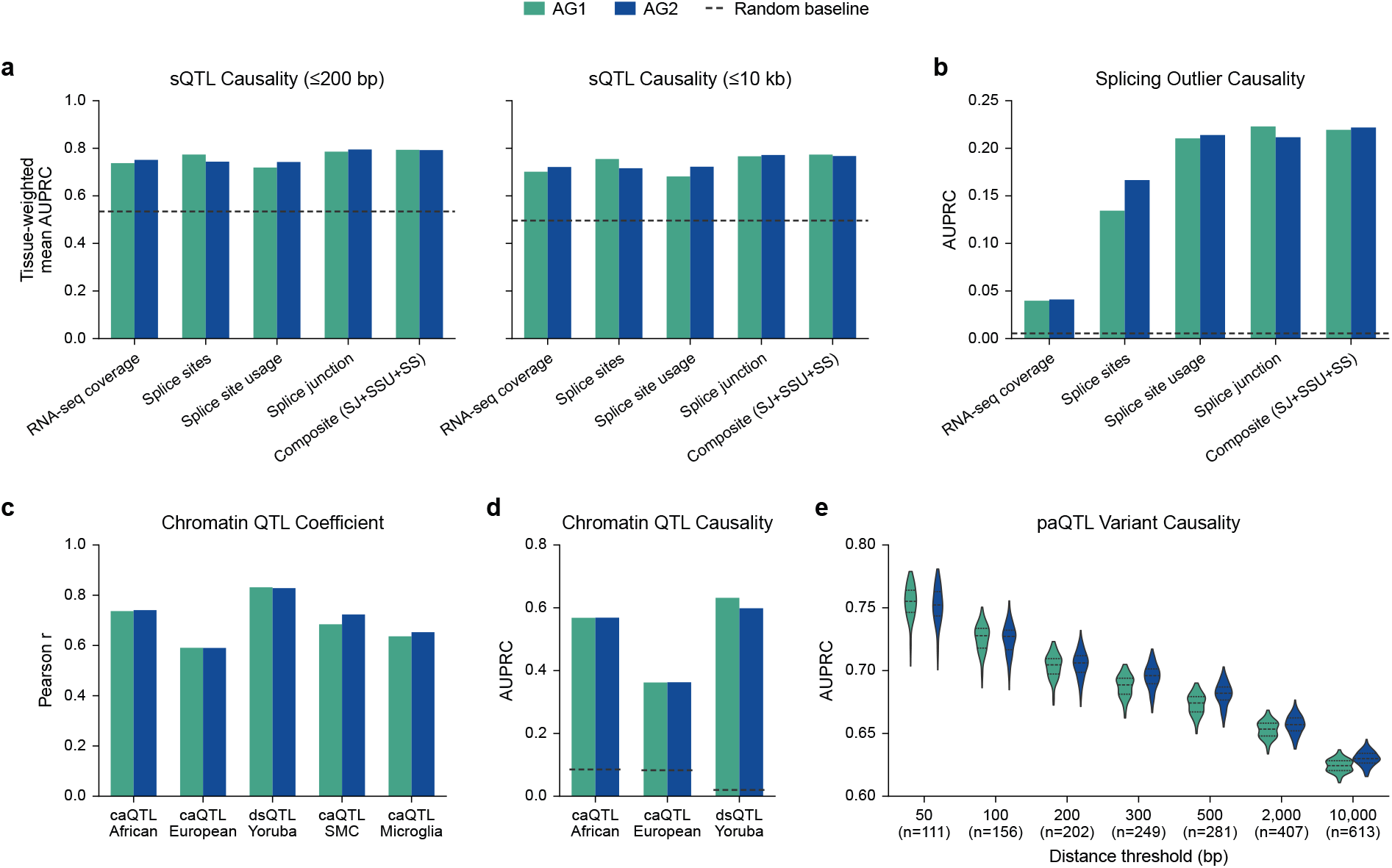
Benchmark comparison of AlphaGenome 2 versus AlphaGenome 1 across core splicing, chromatin, and polyadenylation variant tasks. **(a)** Splicing QTL (sQTL) causality prediction. Tissue-weighted mean area under the precision–recall curve (AUPRC) comparing AlphaGenome 1 (AG1) and AlphaGenome 2 (AG2) across five variant scoring heads (RNA-seq coverage, splice sites, splice site usage, splice junction, and composite “SJ+SSU+SS”) for proximal (*≤*200 bp, left) and distal (*≤*10 kb, right) fine-mapped GTEx sQTLs. Dashed horizontal lines denote random baseline performance. **(b)** Splicing outlier causality prediction. Comparison of AUPRC between AG1 and AG2 across splicing and RNA coverage variant scorers for distinguishing rare GTEx splicing-outlier-associated variants from matched controls. **(c)** Chromatin QTL effect size prediction. Bar plot comparing Pearson correlation (*r*) between predicted and observed variant effect sizes across chromatin accessibility and DNase hypersensitivity QTL benchmarks (“caQTL African”, “caQTL European”, “dsQTL Yoruba”, “caQTL SMC” in smooth muscle cells, and “caQTL Microglia”) for AG1 and AG2. **(d)** Chromatin QTL causality prediction. Comparison of AUPRC between AG1 and AG2 for distinguishing putatively causal chromatin QTLs from matched negative controls across the “caQTL African”, “caQTL European”, and “dsQTL Yoruba” callsets. Dashed horizontal lines indicate random baseline AUPRC. **(e)** Polyadenylation QTL (paQTL) causality prediction. Violin plots showing AUPRC distributions across *n* = 100 random negative-matching permutations (Methods) for AG1 and AG2 on fine-mapped GTEx paQTLs stratified by maximum distance threshold to the target polyadenylation site (from 50 bp to 10,000 bp; x-axis labels give the number of positive paQTLs per threshold). Violins span the observed data range; dashed horizontal lines inside each violin indicate the median and the 25th and 75th percentiles.

**Supplementary Figure 3.**
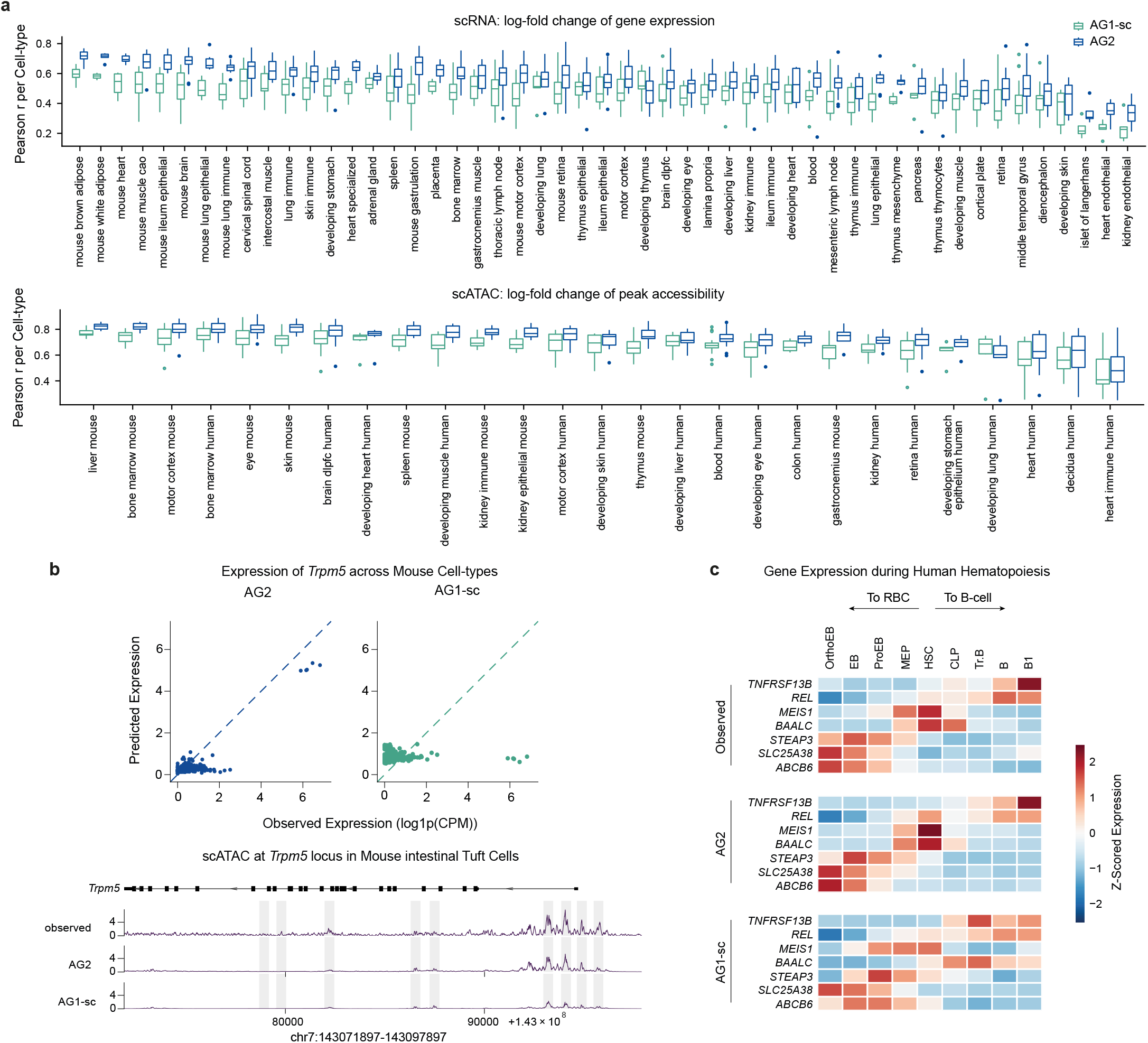
Single-cell gene expression and chromatin accessibility log fold change predictions across human and mouse tissues. **(a)** Cross-cell-type log fold change prediction across human and mouse tissues. Box plots showing the distribution of Pearson correlation (*r*) per cell type between observed and predicted cross-cell-type log fold changes (LFCs) on held-out genomic intervals for single-cell gene expression (top row; 51 comparison groups, *n* = 4–30 cell types per group) and single-cell chromatin accessibility peaks (bottom row; 27 comparison groups, *n* = 5–23 cell types per group; per-group counts in Supplementary Table 8) across diverse mouse and human primary tissues and developmental atlases, comparing AG1 fine-tuned on single-cell data (AG1-sc) and AG2. Box plots display the median (center line), the 25th and 75th percentiles (box bounds), and the whiskers extend to 1.5 times the interquartile range from the box bounds; points beyond whiskers indicate outlier cell types. **(b)** Cell-type-specific expression and chromatin accessibility of *Trpm5* in mouse intestinal tuft cells. Top: Scatterplots of predicted versus observed gene expression (log(CPM + 1)) for *Trpm5* across all profiled mouse primary cell types for AG2 (left) and AG1-sc (right), highlighting selective high expression in chemosensory tuft cells. Bottom: Observed versus predicted single-cell ATAC-seq (scATAC-seq) coverage tracks across the *Trpm5* locus in mouse intestinal tuft cells for observed, AG2, and AG1-sc, with gray vertical bars highlighting cell-type-specific accessible regulatory peaks accurately recovered by AG2. **(c)** Fidelity of predicted gene expression trajectories during human hematopoiesis. Heatmaps displaying row-normalized (Z-scored) observed (top), AG2-predicted (middle), and AG1-sc-predicted (bottom) gene expression across primary hematopoietic differentiation stages from hematopoietic stem cells (HSC) toward erythroid (megakaryocyte–erythroid progenitor (MEP), proerythroblast (ProEB), erythroblast (EB), orthochromatic erythroblast (OrthoEB) *→* red blood cell (RBC)) and lymphoid (common lymphoid progenitor (CLP), transitional B (Tr.B), mature B (B), B1 *→* B-cell) lineages for key lineage-defining marker genes (*TNFRSF13B*, *REL*, *MEIS1*, *BAALC*, *STEAP3*, *SLC25A38*, and *ABCB6*).

**Supplementary Figure 4.**
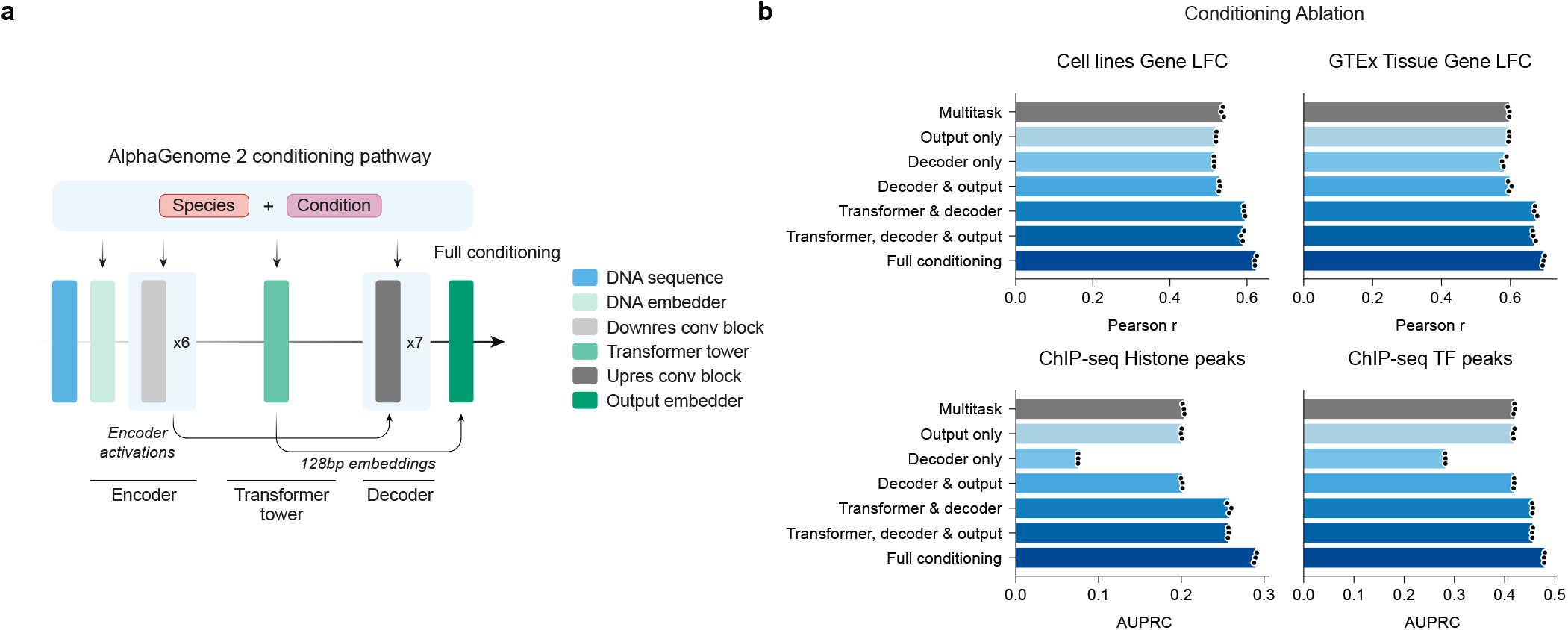
Architectural ablation of biological conditioning pathways in AlphaGenome 2. **(a)** Schematic of the conditioning pathways evaluated across network stages. Species and biological condition embeddings are summed and injected via FiLM across network modules: standard AG2 (“Full conditioning”) modulates the sequence encoder (stem and 6 downsampling convolutional block stages), central transformer tower (128 bp resolution), and sequence decoder (7 upsampling convolutional block stages), whereas restricted ablation configurations selectively enable FiLM in subsets of these stages or in the pointwise output embedder. **(b)** Impact of conditioning placement on held-out genome prediction accuracy. Bar plots comparing model configurations where biological conditioning is restricted to specific architectural stages (“Multitask” baseline without conditioning, “Output only”, “Decoder only”, “Decoder & output”, “Transformer & decoder”, “Transformer, decoder & output”, and “Full conditioning”). Note that “Decoder only” does not condition assays like ChIP-seq defined only at 128-bp resolution. Top row: Cross-condition gene expression log fold change (LFC) Pearson correlation (*r*) across ENCODE cell lines (left) and GTEx tissues (right). Bottom row: Consensus peak classification area under the precision–recall curve (AUPRC) across ChIP-seq histone modifications (left) and ChIP-seq transcription factors (right). Dots indicate individual replicate training runs (*n* = 3 seeds per configuration).

**Supplementary Figure 5.**
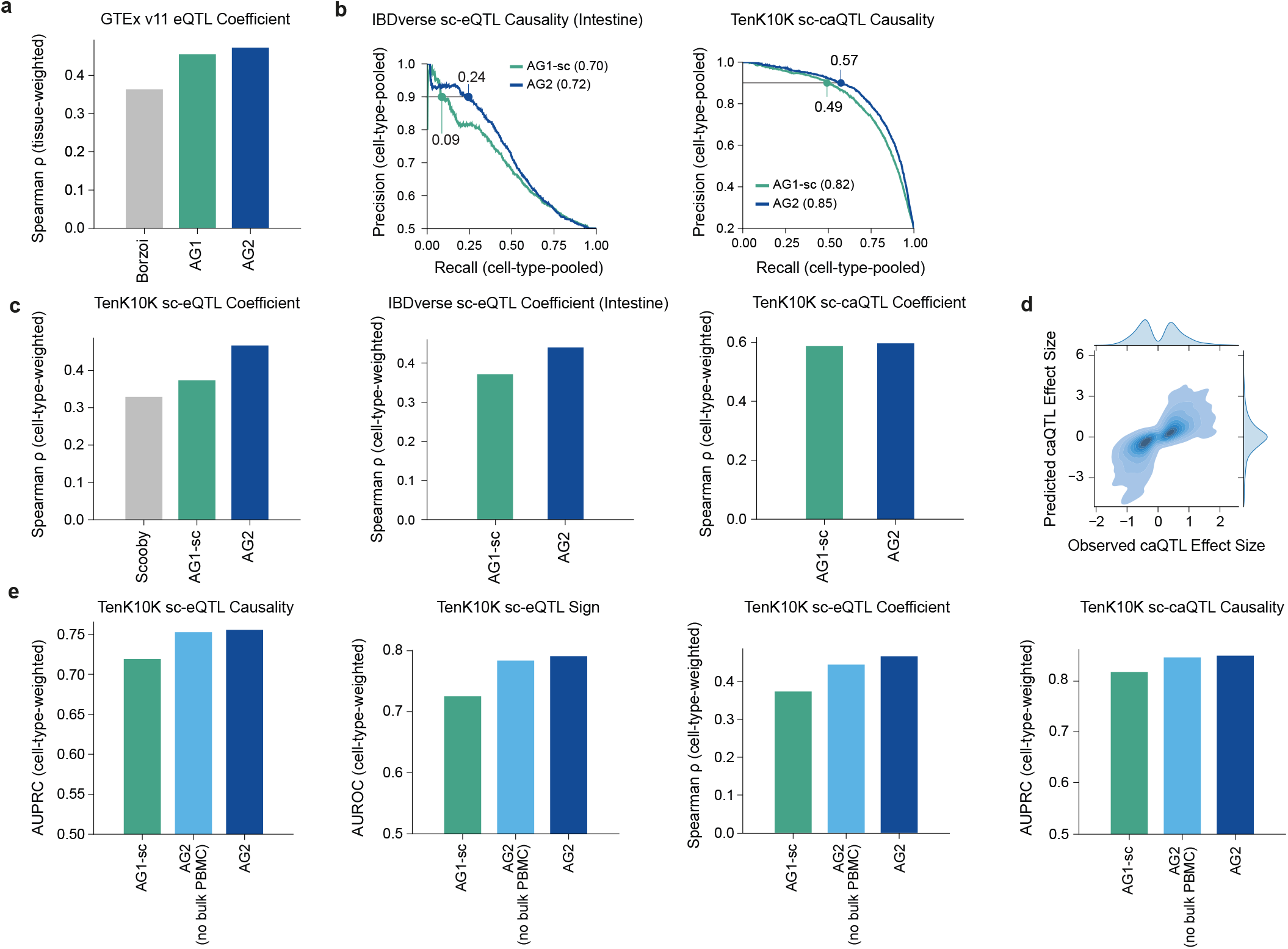
Extended evaluation of bulk tissue and single-cell expression and chromatin accessibility QTL predictions. **(a)** GTEx v11 eQTL effect size prediction. Tissue-weighted mean Spearman rank correlation (*ρ*) between predicted and observed fine-mapped eQTL effect sizes across 49 GTEx tissues for Borzoi, AG1 and AG2. **(b)** Supplementary sc-eQTL and sc-caQTL evaluations. Left: Cell-type-pooled precision–recall (1:1 negative-to-positive ratio) curves for 3524 causal sc-eQTLs across 47 cell types (IBDverse). Right: Precision–recall curves for 5116 causal sc-caQTLs (4:1 negative-to-positive ratio) across 24 cell types (TenK10K). Point callouts indicate recall at 90% precision. **(c)** Single-cell QTL effect size prediction. Cell-type-weighted mean Spearman rank correlation (*ρ*) for single-cell eQTLs across 28 primary immune cell types (left; Scooby, AG1-sc and AG2), single-cell eQTLs across 47 intestinal cell types (middle; AG1-sc and AG2) and single-cell chromatin accessibility QTLs across 24 primary immune cell types (right; AG1-sc and AG2). **(d)** Predicted versus observed single-cell caQTL effect sizes. Contour density plot and marginal distributions comparing AG2-predicted versus observed allelic effect sizes across TenK10K fine-mapped sc-caQTLs. **(e)** Effect of holding out bulk PBMC tracks on TenK10K single-cell QTL performance. Cell-type-weighted mean sc-eQTL causality (AUPRC), sc-eQTL sign (AUROC), sc-eQTL effect size (Spearman *ρ*) and sc-caQTL causality (AUPRC) for AG1-sc, AG2 trained with all bulk PBMC epigenomic and transcriptional tracks held out (AG2 (no bulk PBMC)), and AG2.

**Supplementary Figure 6.**
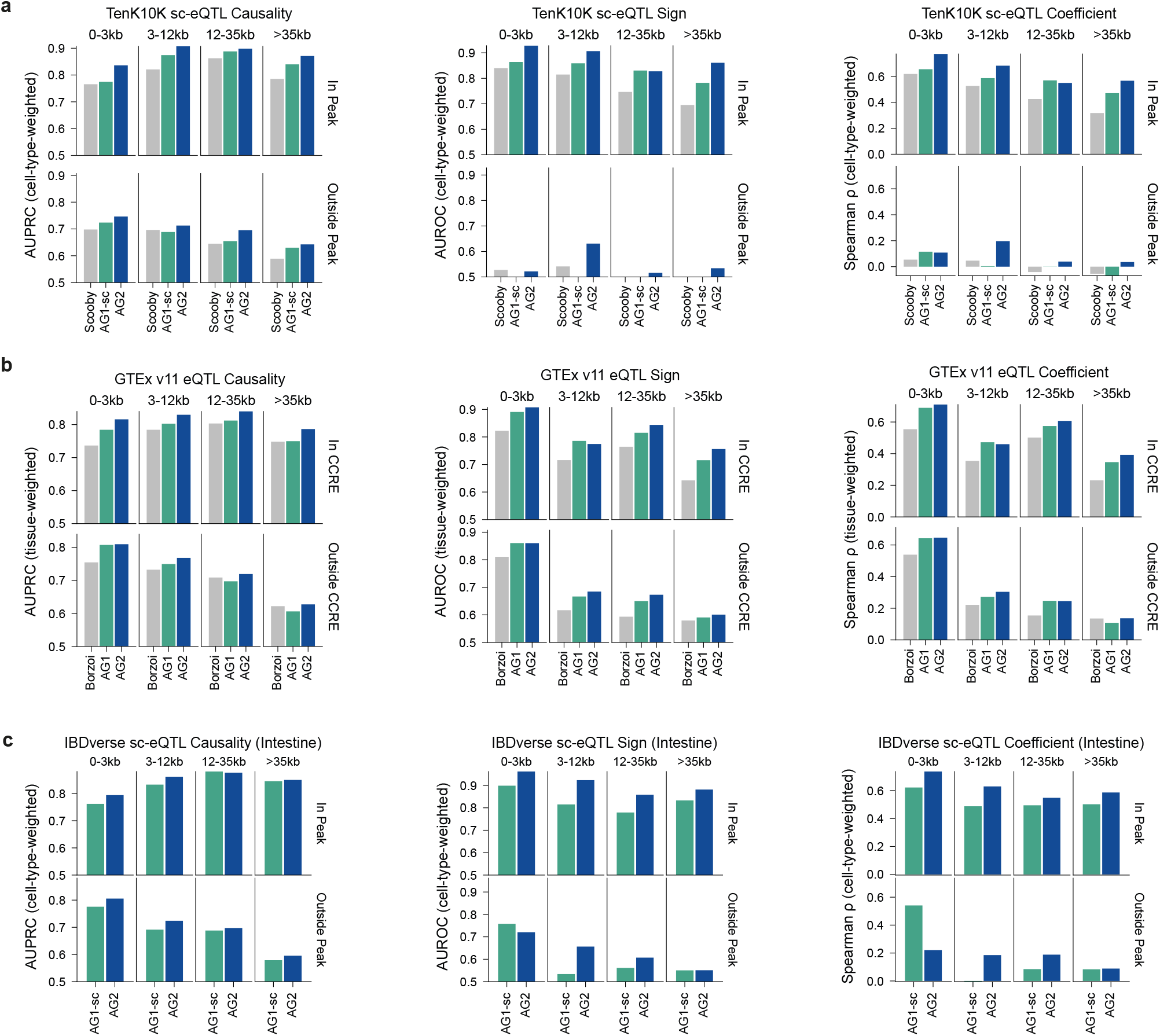
Extended evaluation of bulk tissue and single-cell expression and chromatin accessibility QTL predictions. **(a)** TenK10K sc-eQTL performance stratified by distance to the target gene TSS (0–3, 3–12, 12–35 and >35 kb) and by whether the variant lies inside (top row) or outside (bottom row) an active primary blood scATAC-seq peak. Cell-type-weighted mean causality (AUPRC, left), sign (AUROC, middle) and effect size (Spearman *ρ*, right) for Scooby, AG1-sc and AG2. AUROC and AUPRC y-axes begin at random performance (0.5), small deviations below this are omitted from the plot for simplicity. **(b)** GTEx v11 eQTL performance stratified by distance to TSS (as in **a**) and by overlap with ENCODE SCREEN candidate *cis*-regulatory elements (in cCRE, top row; outside cCRE, bottom row). Tissue-weighted mean causality (AUPRC, left), sign (AUROC, middle) and effect size (Spearman *ρ*, right) for Borzoi, AG1 and AG2. **(c)** IBDverse sc-eQTL performance stratified by distance to TSS (as in **a,b**) and by overlap with active primary blood and colon scATAC-seq peaks (in peaks, top row; outside peaks, bottom row). Tissue-weighted mean causality (AUPRC, left), sign (AUROC, middle) and effect size (Spearman *ρ*, right) for AG1-sc and AG2.

**Supplementary Figure 7.**
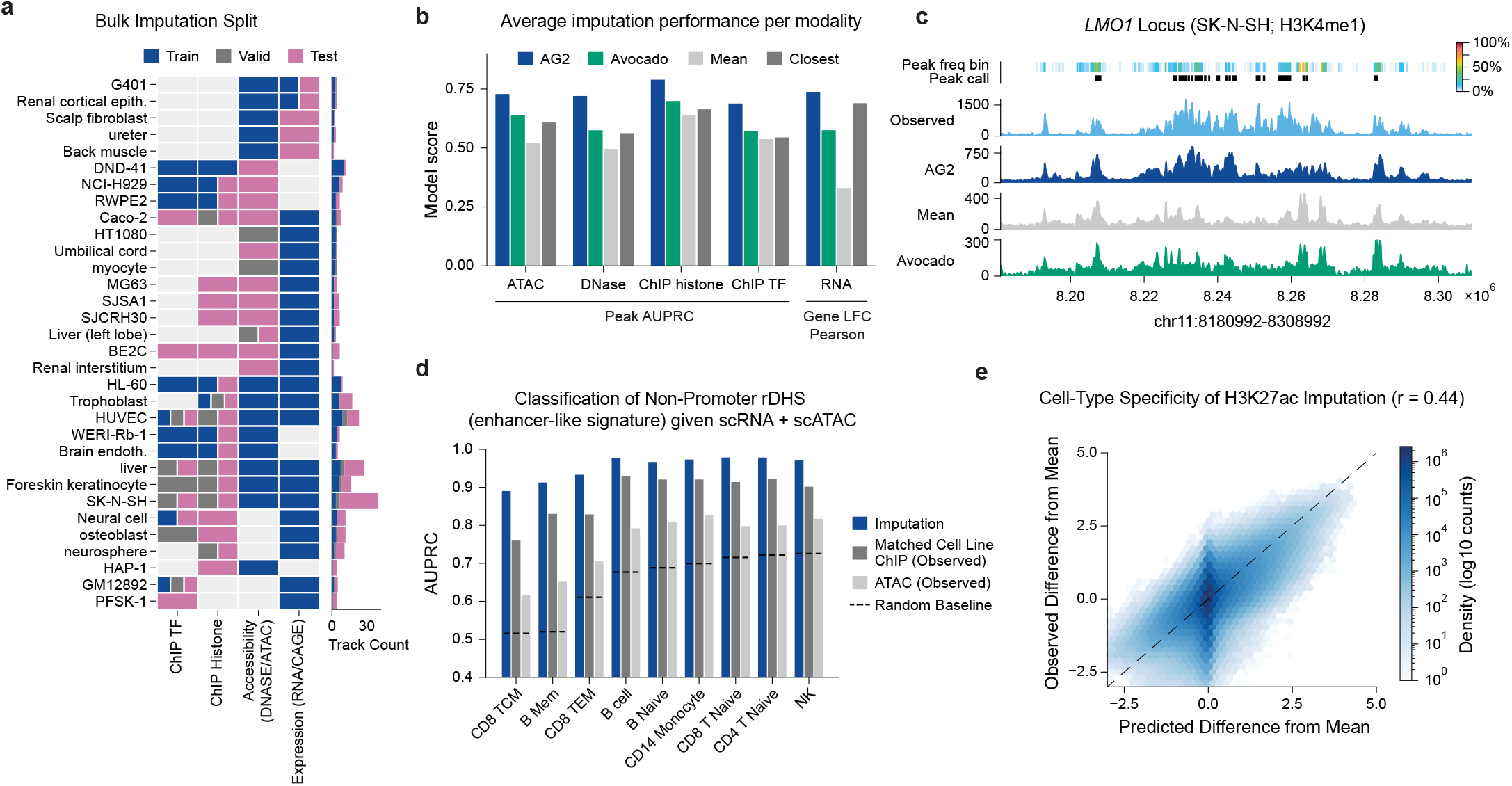
Cross-modality bulk epigenomic imputation and primary-cell enhancer annotation with AlphaGenome 2. **(a)** Bulk epigenomic imputation dataset split. Overview of training, validation, and held-out test tracks across 32 human conditions with held-out tracks (tissues, cell lines, and primary cell types) spanning four bulk assay categories: transcription factor (TF) ChIP-seq (“ChIP TF”), histone modification ChIP-seq (“ChIP Histone”), chromatin accessibility (“Accessibility (DNASE/ATAC)”), and gene expression (“Expression (RNA/CAGE)”). Right bar plot shows track counts per condition, stacked by split. **(b)** Average imputation performance across modalities comparing AG2, retrained Avocado tensor factorization, cross-condition channel mean (“Mean”), and five-closest training-track oracle (“Closest”). Left four groups: support-weighted consensus peak area under the precision–recall curve (AUPRC) for ATAC-seq, DNase-seq, histone ChIP-seq, and TF ChIP-seq. Far right: cross-condition gene log fold change Pearson *r* (“Gene LFC Pearson”) for bulk RNA-seq. **(c)** Example imputed H3K4me1 profile (held-out track) at the *LMO1* locus in SK-N-SH neuroblastoma cells. Tracks show consensus peak empirical frequency bins (0% to 100%, top color bar), experimentally called peaks (“Peak call”, ticks), and observed versus predicted H3K4me1 coverage for AG2, “Mean”, and Avocado. **(d)** Classification of open non-promoter representative DNase I hypersensitive sites (rDHSs) with enhancer-like signatures. Bar plot comparing AUPRC across nine primary peripheral blood mononuclear cell (PBMC) types (CD8 TCM, B Mem, CD8 TEM, B cell, B Naive, CD14 Monocyte, CD8 T Naive, CD4 T Naive, and NK) using AG2-imputed H3K27ac (“Imputation”), observed H3K27ac ChIP-seq from a lineage-matched cell line (“Matched Cell Line ChIP (Observed)”), and observed scATAC-seq alone (“ATAC (Observed)”). Dashed horizontal segments indicate random baseline prevalence per cell type. **(e)** Cell-type specificity of H3K27ac imputation. Hexbin density scatterplot comparing predicted versus observed differences from the cross-cell-type mean H3K27ac *z*-score across genome-wide rDHSs in the nine PBMC types (Pearson *r* = 0.44; Methods).

**Supplementary Figure 8.**
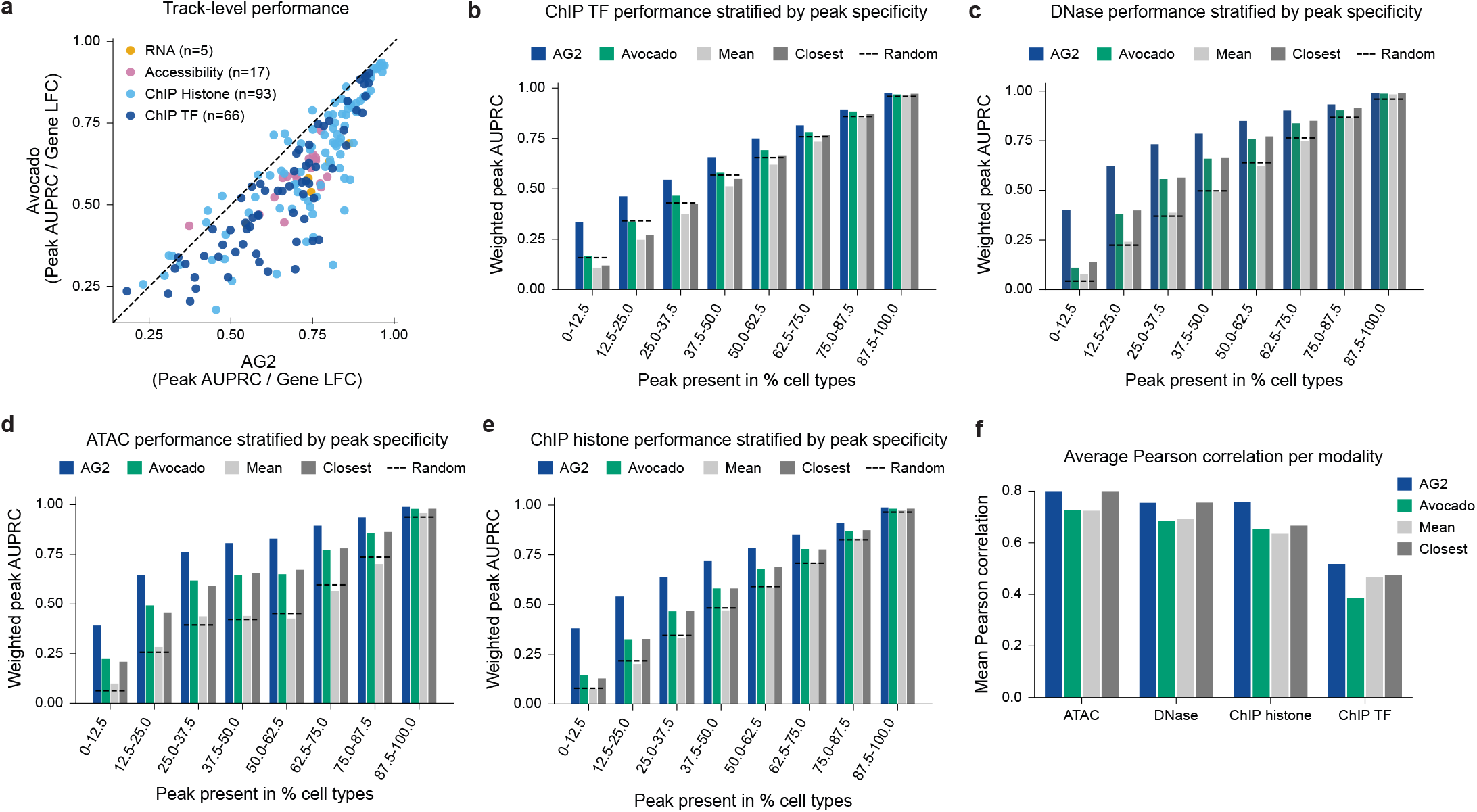
Track-level and frequency-stratified bulk epigenomic imputation benchmarks with AlphaGenome 2. **(a)** Track-level imputation comparison. Scatterplot comparing per-track imputation accuracy (consensus peak area under the precision–recall curve (AUPRC) for chromatin assays; cross-condition gene log fold change Pearson *r* for RNA-seq) between AG2 (x-axis) and retrained Avocado (y-axis) across all 181 evaluated test tracks (RNA-seq, *n* = 5; chromatin accessibility, *n* = 17; histone ChIP-seq, *n* = 93; transcription factor (TF) ChIP-seq, *n* = 66; Methods). Points below the dashed diagonal (*y* = *x*) indicate tracks where AG2 outperforms Avocado (166 of 181 tracks). **(b–e)** Consensus peak classification AUPRC stratified by condition specificity. Support-weighted peak AUPRC across eight empirical peak frequency bins (proportion of profiled conditions in which a consensus peak is called active, in 12.5% increments from rare condition-specific elements, 0–12.5%, to ubiquitous elements, 87.5–100%) comparing AG2, Avocado, cross-condition channel mean (“Mean”), and five-closest training-track oracle (“Closest”) across **(b)** TF ChIP-seq, **(c)** DNase-seq, **(d)** ATAC-seq, and **(e)** histone ChIP-seq. Dashed horizontal segments indicate empirical random baseline prevalence within each frequency bin. **(f)** Continuous profile correlation across chromatin modalities. Bar plot comparing mean genome-wide continuous Pearson *r* at 32 bp resolution across held-out test tracks for ATAC-seq, DNase-seq, histone ChIP-seq, and TF ChIP-seq across AG2, Avocado, “Mean”, and “Closest”.

**Supplementary Figure 9.**
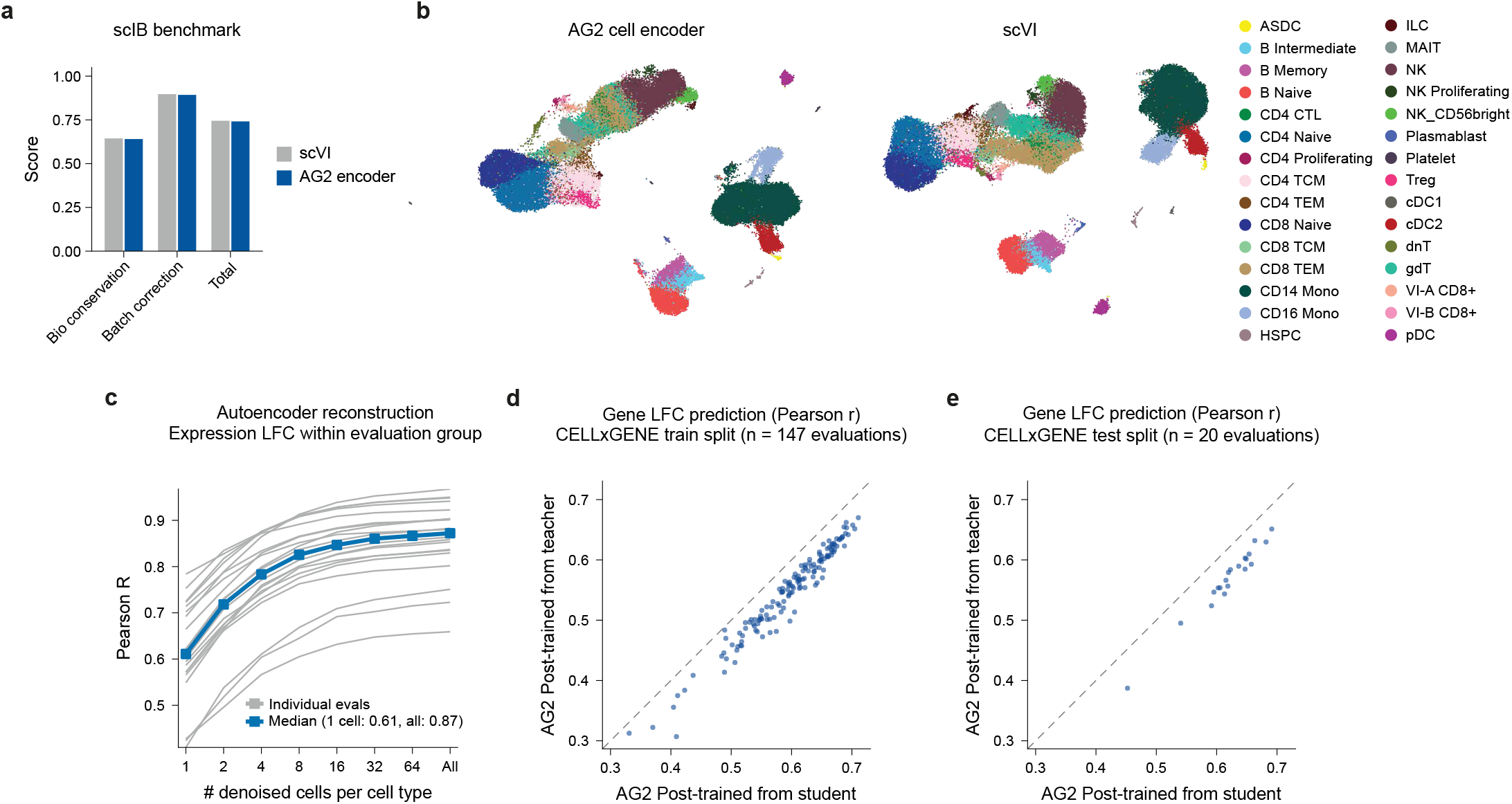
Evaluation of the single-cell expression autoencoder and post-trained AlphaGenome 2. **(a)** Single-cell integration benchmarking (scIB) of cell embeddings on the CITE-seq RNA component of a SARS-CoV-2 vaccination PBMC dataset^1^. The AG2 cell encoder with no dataset-specific training or batch covariates is compared with an scVI model trained on this dataset with donor as the batch covariate. Bars show the composite biological conservation, batch correction and total scores from scib-metrics^2^. **(b)** Uniform manifold approximation and projection (UMAP) of AG2 cell encoder embeddings (left) and scVI latent representations (right) for the same cells, colored by level-2 cell type. **(c)** Recovery of cell-type-specific expression by autoencoder reconstructions across 16 held-out CELLxGENE test-split datasets. For each cell type, cell embeddings were averaged over *k* randomly sampled cells (*k* = 1 to 64, or all cells; 20 trials) and decoded into a denoised profile, which was compared with the observed pseudobulk by cross-gene LFC Pearson *r* across cell types. Individual curves show each dataset alongside the median across datasets. **(d,e)** Gene LFC Pearson *r* for AG2 post-trained from a single pre-trained model prior to condition refinement (y-axis) versus from the distilled student model (x-axis), across evaluation groups from CELLxGENE training-split (**d**; *n* = 147) and test-split (**e**; *n* = 20). Dashed line, *y* = *x*.

**Supplementary Figure 10.**
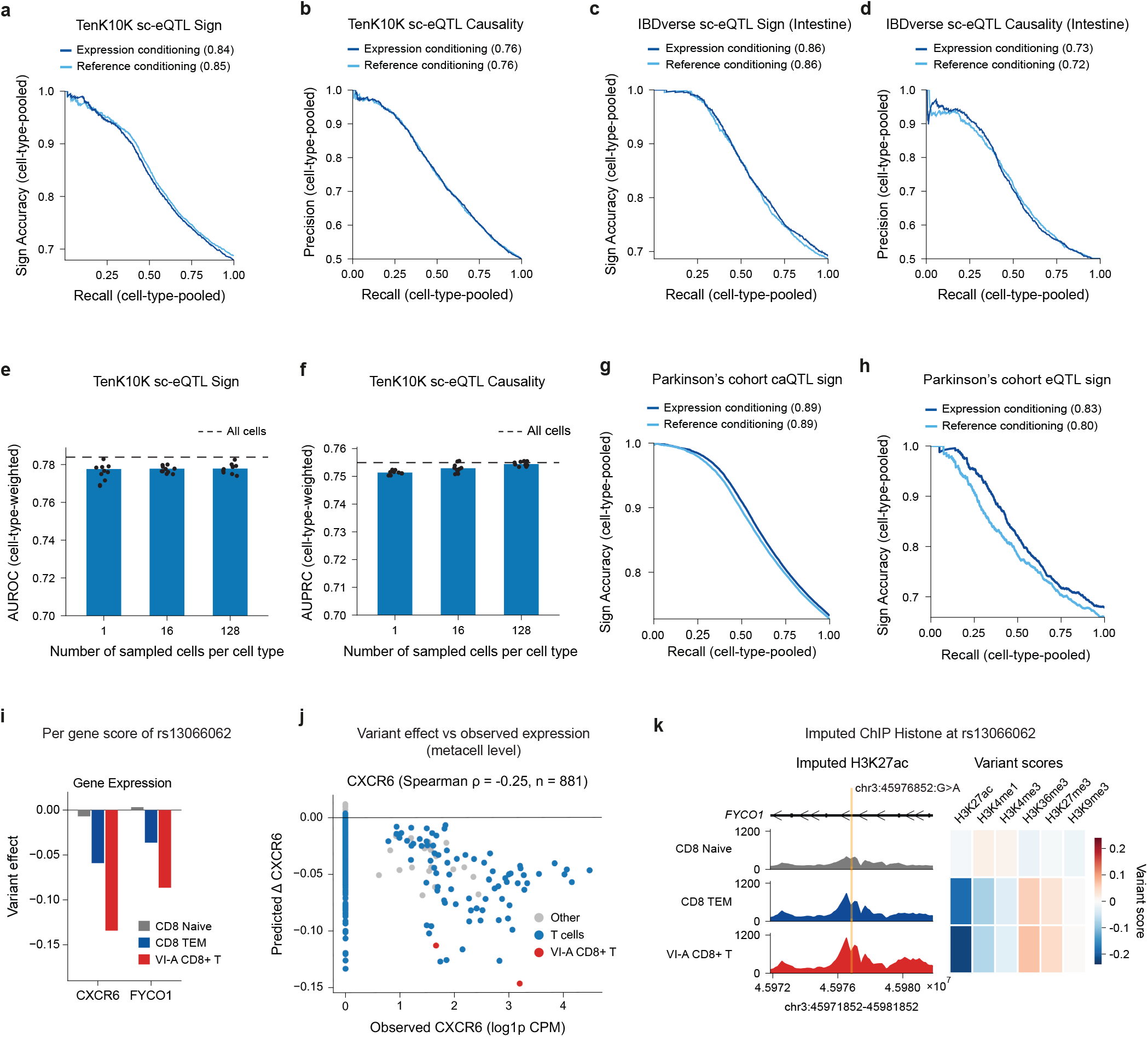
Evaluation of post-trained AlphaGenome 2. (a,b) Cell-type-pooled TenK10K single-cell expression QTL (sc-eQTL) sign accuracy versus recall in **a** and causality precision versus recall in **b** across the 25 cell types matched in an independent PBMC dataset^3,4^. Curves compare expression conditioning on mean cell embeddings from that dataset with reference conditioning on matched reference conditions. Legend values give area under the curve (AUC) in **a** and area under the precision–recall curve (AUPRC) in **b**. **(c,d)** Cell-type-pooled IBDverse sc-eQTL sign accuracy versus recall in **c** and causality precision versus recall in **d** across the 47 intestinal cell types^5^. Curves compare expression conditioning on mean cell embeddings from each cell type’s donor-majority intestinal tissue in the IBDverse dataset with reference conditioning on matched reference conditions. Legend values give AUC in **c** and AUPRC in **d**. **(e,f)** Cell-type-weighted TenK10K sc-eQTL sign AUROC in **e** and causality AUPRC in **f** for post-trained AG2 conditioned on cell embeddings averaged over 1, 16 or 128 randomly sampled cells per cell type^3,4^. Bars show the mean across 10 random trials and points show individual trials. Dashed lines indicate conditioning on all cells. **(g,h)** Sign accuracy versus recall pooled across cell types and brain regions in the Parkinson’s disease cohort^6^ for variants with both chromatin accessibility QTL and allele-specific chromatin accessibility effects in **g** and for overlapping expression QTLs in **h**. Curves compare expression conditioning on mean cell embeddings for each cell type and brain region with reference conditioning on matched reference conditions. Legend values give AUC. **(i)** Predicted effect of rs13066062 on *CXCR6* and *FYCO1* expression for post-trained AG2 conditioned on the mean cell embeddings of naive CD8^+^ T (CD8 Naive), effector memory CD8^+^ T (CD8 TEM) and vaccine-induced group A CD8^+^ T (VI-A CD8^+^ T) cells from the SARS-CoV-2 vaccination dataset^1^. **(j)** Predicted change in *CXCR6* expression caused by rs13066062 versus observed *CXCR6* expression (log(CPM + 1)) for 881 metacells from the same dataset (Spearman *ρ* = *−*0.25). Points indicate VI-A CD8^+^ T, other T-cell and remaining metacells. **(k)** Imputed histone ChIP-seq profiles and variant effects at rs13066062 for the three CD8^+^ T-cell states in **i** (Methods). Left: imputed H3K27ac signal across a 10-kb window at *FYCO1*, centered on rs13066062 (`chr3:45976852:G>A`; orange line). Right: predicted variant scores across H3K27ac, H3K4me1, H3K4me3, H3K36me3, H3K27me3 and H3K9me3.

**Supplementary Figure 11.**
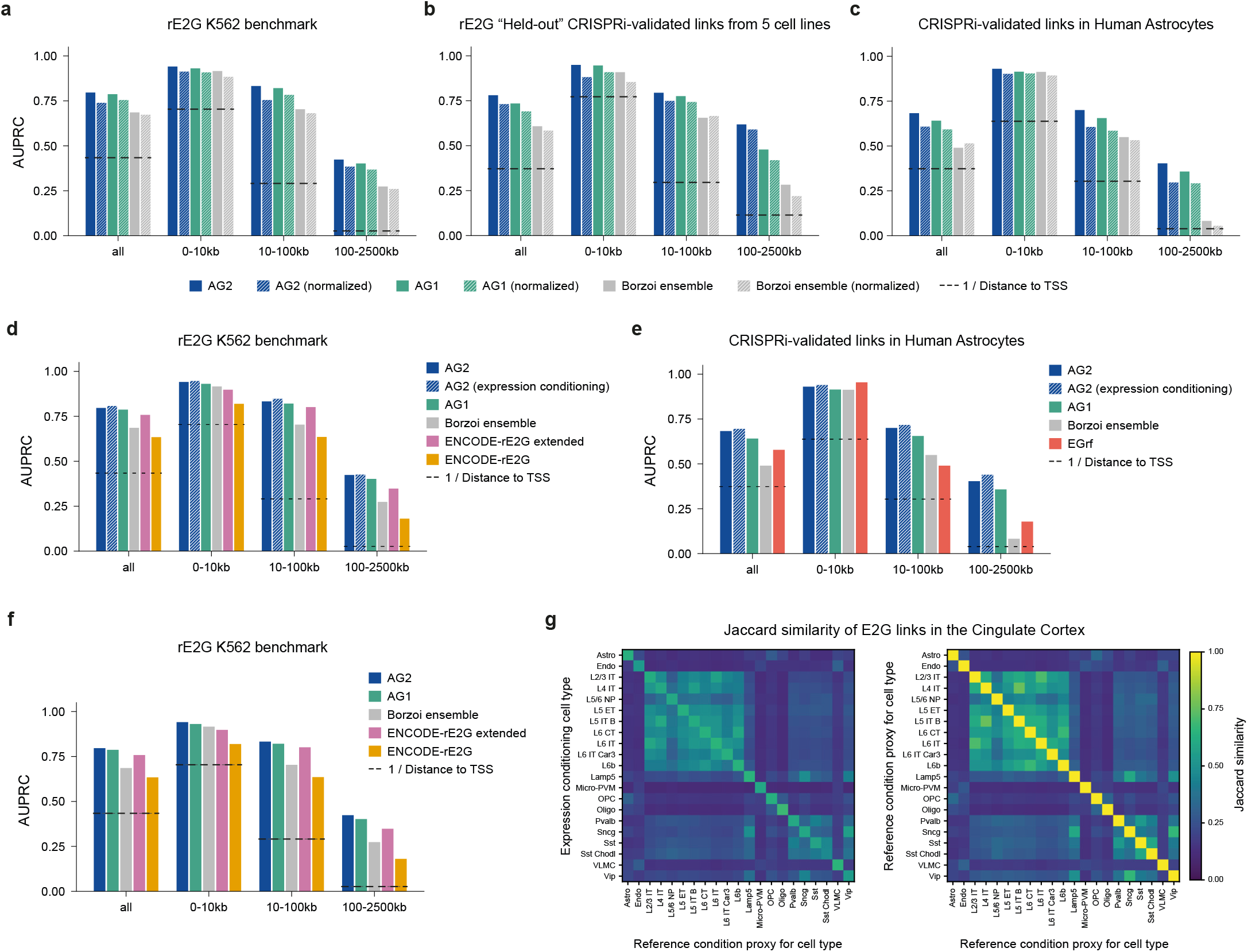
Extended evaluation and ablations of enhancer–gene linking across CRISPRi benchmarks and primary brain cell types. **(a–c)** Comparison of unnormalized (solid bars) versus window-normalized (hatched bars; divided by the mean absolute attribution across the input window) gradient attribution scores on the ENCODE-rE2G K562 CRISPRi benchmark (**a**)^7^, the held-out ENCODE-rE2G CRISPRi benchmark across five cell lines (**b**)^7^, and the human astrocyte CRISPRi benchmark (**c**)^8^ for AG2, AG1^9^, and the four-replicate Borzoi ensemble^10^. **(d,e)** Comparison of reference-conditioned AG2 (solid bars) and post-trained AG2 using expression conditioning on mean cell embeddings from the corresponding scRNA-seq dataset (hatched bars) on the ENCODE-rE2G K562 benchmark (**d**)^7^ and the human astrocyte benchmark (**e**)^8^, alongside AG1, the Borzoi ensemble, and ENCODE-rE2G extended and ENCODE-rE2G (**d**) or EGrf (**e**). **(g)** Enhancer–gene (E2G) linking performance on the full ENCODE-rE2G K562 CRISPRi benchmark^7^, comparing AG2, AG1, the Borzoi ensemble, ENCODE-rE2G extended, and ENCODE-rE2G. In **(a–f)**, bars show area under the precision–recall curve (AUPRC) across all tested element–gene pairs and stratified by enhancer distance to the target gene transcription start site (TSS; 0–10 kb, 10–100 kb, and 100–2500 kb); horizontal dotted lines indicate the inverse TSS distance baseline. **(g)** Pairwise Jaccard similarity of predicted E2G links for protein-coding genes at caQTL-ASCA peaks in the cingulate cortex of the Parkinson’s disease cohort^6^, comparing post-trained AG2 conditioned on mean cell embeddings for each cell type (rows) with AG2 conditioned on the corresponding reference condition proxies from the training corpus (columns; left), and comparing the reference condition proxies with one another across the same layout (right).

**Supplementary Figure 12.**
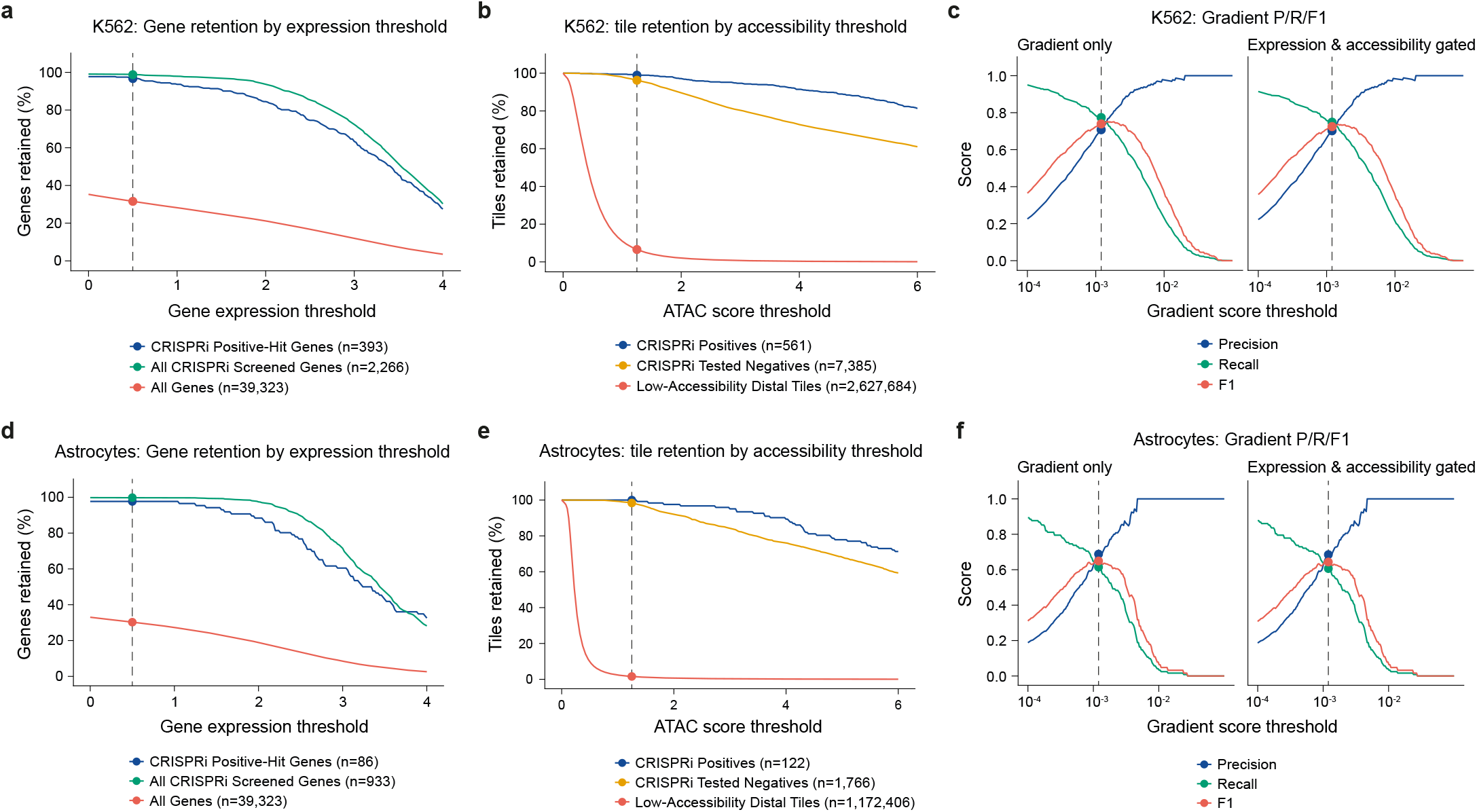
Threshold selection and gating diagnostics for the AlphaGenome 2 enhancer–gene linking atlas. **(a)** Percentage of genes retained in K562 as a function of the predicted log-transformed gene expression threshold across CRISPRi positive-hit target genes, all CRISPRi-screened genes, and all protein-coding and long non-coding RNA (lncRNA) genes in GENCODE v46; dashed vertical line indicates the selected target gene expression threshold (*ŷg ≥* 0.5; Methods). **(b)** Percentage of candidate enhancer tiles retained in K562 as a function of the predicted local chromatin accessibility threshold across CRISPRi-validated positive enhancers, CRISPRi tested negative elements, and low-accessibility distal tiles (located >2 kb from the target gene TSS and >2 kb from any ENCODE ATAC-seq narrowPeak in K562); dashed vertical line indicates the selected accessibility threshold (scATAC *≥* 1.25). **(c)** Precision, recall, and *F*­ score across gradient attribution score thresholds on the K562 ENCODE CRISPRi enhancer–gene (E2G) benchmark (with each CRISPRi element–gene pair mapped to its best-overlapping 500-bp atlas tile), shown before and after applying the target gene expression and local chromatin accessibility gates; dashed vertical line indicates the selected gradient attribution threshold (score *≥* 0.0012). **(d–f)** Same as **(a–c)**, evaluated on the astrocytes CRISPRi E2G benchmark.

**Supplementary Figure 13.**
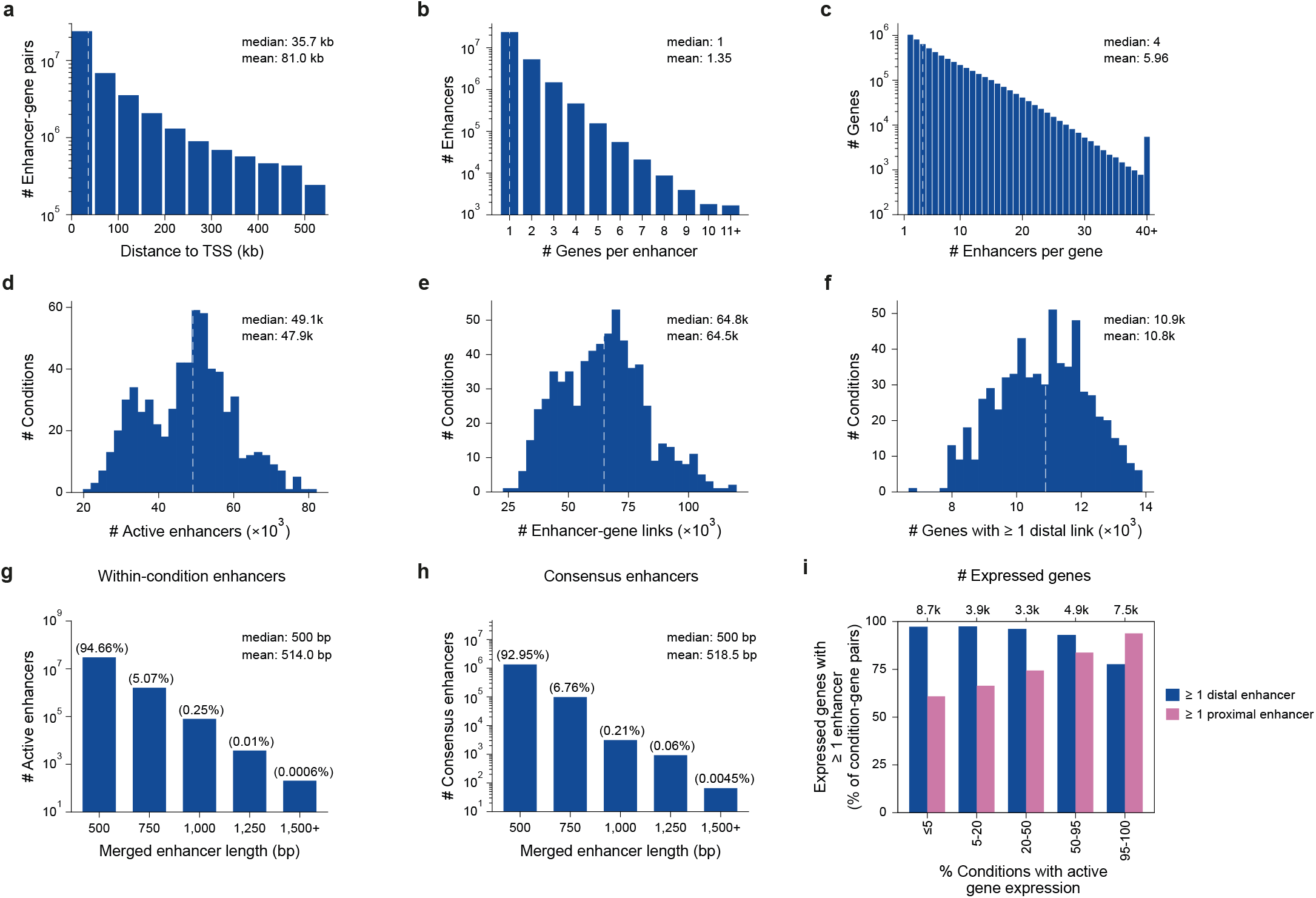
Summary statistics and regulatory properties of the AlphaGenome 2 enhancer–gene linking atlas across 640 primary single-cell conditions. **(a)** Distribution of genomic distances (in kb) between active distal enhancer midpoints and the representative transcription start site (TSS) of their linked target genes across all predicted distal enhancer–gene (E2G) links (>500 bp from the TSS; median = 35.7 kb, mean = 81.0 kb). **(b)** Distribution of the number of target genes linked per active distal enhancer interval within each condition (median = 1, mean = 1.35). **(c)** Distribution of the number of linked distal enhancers per regulated target gene within each condition (median = 4, mean = 5.96). **(d)** Distribution of the number of active distal enhancer intervals per condition across the 640 primary single-cell conditions (median = 49,100, mean = 47,900). **(e)** Distribution of the number of predicted distal E2G links per condition across the 640 primary single-cell conditions (median = 64,800, mean = 64,500). **(f)** Distribution of the number of target genes with at least one linked distal enhancer per condition across the 640 primary single-cell conditions (median = 10,900, mean = 10,800). In **(a–f)**, dashed vertical lines indicate the median. **(g,h)** Distribution of merged distal enhancer interval lengths (in bp) resulting from the 250-bp-stepped summit-merging procedure for within-condition active enhancers (**g**), and cross-condition consensus enhancers (**h**). Percentages of enhancers are indicated above each bar. **(i)** Percentage of expressed condition–gene pairs (*ŷg ≥* 0.5; Methods) with at least one linked distal enhancer (>500 bp from the TSS) or at least one promoter-proximal enhancer (*≤*500 bp from the TSS), stratified by the percentage of conditions in which the gene is actively expressed (top axis shows total expressed genes per bin).

**Supplementary Figure 14.**
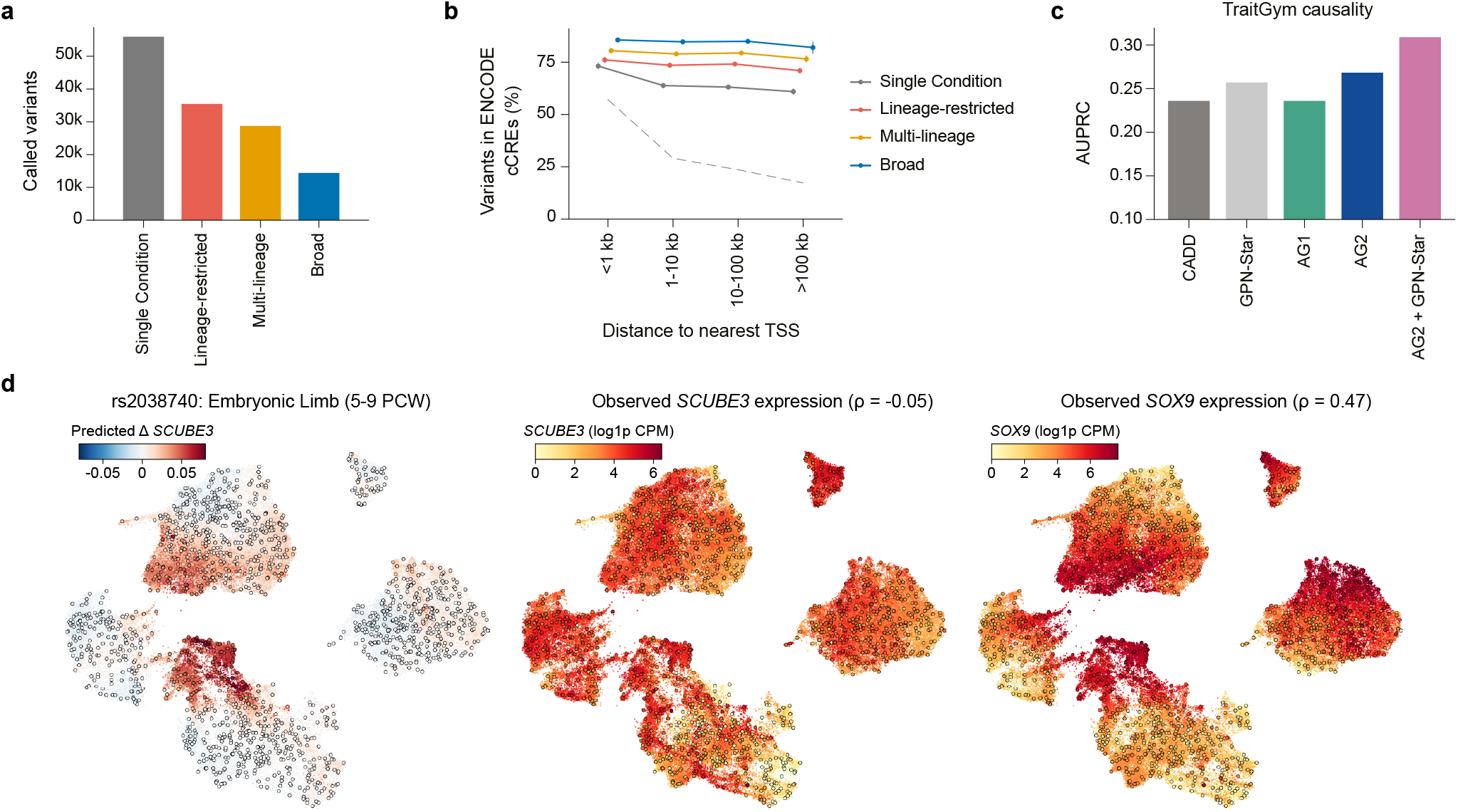
Properties of AlphaGenome 2 GWAS variant calls, complex-trait variant causality benchmarking and developmental expression at the *SCUBE3* locus. **(a)** Number of called variants (*n* = 134,813 across the GWAS panel) stratified by cell-type specificity across non-negative matrix factorization (NMF) transcriptional identity clusters (Methods): single condition (*n* = 56,189), lineage-restricted (1 lineage cluster; *n* = 35,447), multi-lineage (2–3 lineage clusters; *n* = 28,751) and broad (*≥*4 lineage clusters; *n* = 14,426). Variants with multiple calls are assigned the broadest category among their calls. **(b)** Percentage of variants overlapping an ENCODE v4 candidate *cis*-regulatory element (cCRE) across bins of distance to the nearest protein-coding transcription start site (TSS), for called variants stratified by cell-type specificity (solid lines) versus uncalled variants (dashed gray line) (Methods). Error bars, 95% confidence intervals (CIs) from a 1-Mb genomic block bootstrap. **(c)** Zero-shot complex-trait variant causality evaluation on the TraitGym benchmark (*n* = 11,400 variants; fine-mapped causal variants with PIP > 0.9 versus control variants matched on TSS distance, minor allele frequency and linkage disequilibrium (LD) score at a 1:9 ratio)^11^. Bars show area under the precision–recall curve (AUPRC) for CADD^12^, GPN-Star^13^, AG1^9^, AlphaGenome 2 (AG2; maximum zero-shot score across observed tracks from 500 conditions and nine modalities) and an equal-weight average of min–max normalized AG2 and GPN-Star scores (AG2 + GPN-Star) (Methods). **(d)** Predicted effect of the standing-height GWAS variant rs2038740 (`chr6:35146765:C>T`) on *SCUBE3* expression (left) compared with observed pseudobulk expression (log(CPM + 1)) of *SCUBE3* (middle) and *SOX9* (right; Spearman *ρ* = 0.47) across 1575 metacells from the human embryonic limb lateral plate mesoderm (5–9 post-conception weeks (PCW); *n* = 100,935 cells across 34 cell types)^14^. Each metacell is conditioned on the mean embedding of its cells; cells are shown on a UMAP of cell-encoder embeddings and colored by their metacell’s value; dots, metacell mean embeddings.

**Supplementary Figure 15.**
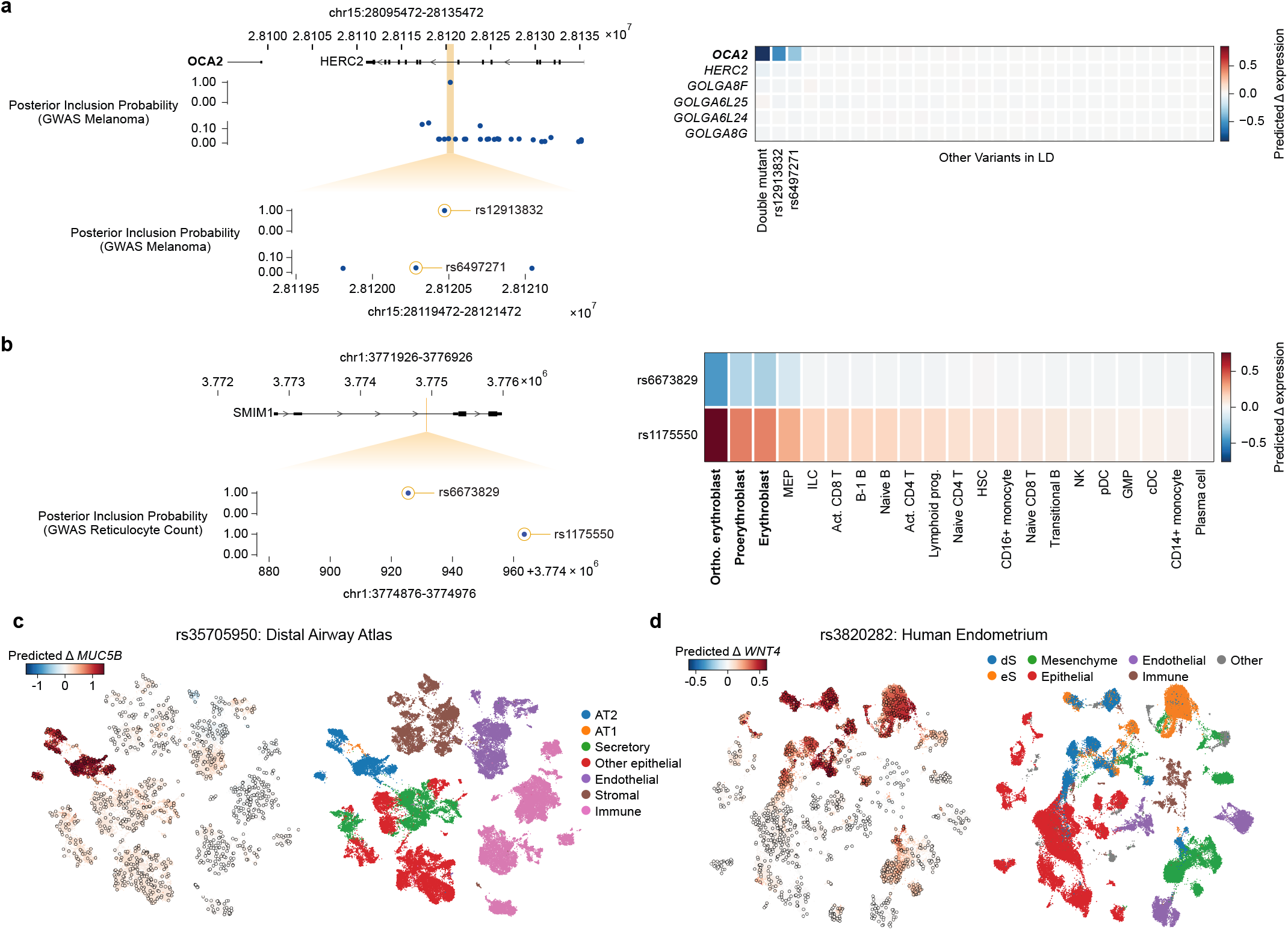
AlphaGenome 2 resolves multiple causal variants, allelic series and disease-relevant cell types at GWAS loci. **(a)** Dissection of two adjacent melanoma GWAS credible sets at the *HERC2*–*OCA2* locus^15^. Top left: 40-kb window showing *OCA2* and *HERC2* and posterior inclusion probabilities (PIPs) for the two credible sets. Bottom left: Zoomed 2-kb window within the *HERC2* intronic enhancer showing the single-variant credible set (rs12913832, `chr15:28120472:A>G`, PIP = 1.00; top track, circled) and the 27-variant credible set containing rs6497271 (`chr15:28120285:A>G`, PIP = 0.03, 187 bp away; bottom track, circled). Right: AG2-predicted expression effects (log fold change) in primary skin melanocytes across nearby protein-coding genes for the haplotype carrying both alternative alleles (“Double mutant”), rs12913832, rs6497271 and the remaining 26 credible-set variants in LD (“Other Variants in LD”). **(b)** Allelic series of two fine-mapped reticulocyte-count GWAS variants with opposing effects at the *SMIM1* locus^16,17^. Top left: 5-kb window around *SMIM1*. Bottom left: Zoomed 100-bp window showing two single-variant credible sets 38 bp apart: rs6673829 (`chr1:3774926:G>A`, PIP > 0.99; top track, circled) and rs1175550 (`chr1:3774964:A>G`, PIP > 0.99; bottom track, circled). Right: AG2-predicted effects on *SMIM1* expression (log fold change) across 21 primary bone-marrow hematopoietic cell types. **(c)** Cell-type resolution of the idiopathic pulmonary fibrosis risk variant rs35705950 (`chr11:1219991:G>T`) at the *MUC5B* locus across 1088 metacells from a human distal airway single-cell atlas (*n* = 69,830 cells across 19 cell types)^18^. Left: Predicted effect on *MUC5B* expression (log fold change). Right: Coarse cell-type labels, highlighting alveolar type 2 (AT2) and type 1 (AT1) cells. **(d)** Cell-type resolution of the endometriosis and reproductive-trait variant rs3820282 (`chr1:22141722: C>T`) at the *WNT4* locus across 1183 metacells from a human endometrial single-cell atlas (*n* = 75,844 cells across 16 cell types)^19^. Left: Predicted effect on *WNT4* expression (log fold change). Right: Coarse cell-type labels, highlighting decidualized (dS) and non-decidualized (eS) endometrial stromal cells. In **(c,d)**, each metacell is conditioned on the mean embedding of its cells; cells are shown on a UMAP of cell-encoder embeddings and colored by their metacell’s value; dots, metacell mean embeddings.

### Supplementary Note 1 *In silico* study of individual GWAS loci

Beyond large-scale analyses, AG2 can also be used to study individual loci *in silico*. For example, the *HERC2*– *OCA2* locus contains two adjacent credible sets associated with melanoma^15^. The first is fine-mapped to a single variant, rs12913832. This variant is a major determinant of eye and skin pigmentation in individuals of European ancestry^20^. It reduces *OCA2* expression by weakening a chromatin loop between an enhancer and the *OCA2* promoter^21^. The second locus contains 27 variants, none with a posterior inclusion probability (PIP) above 0.15. This includes rs6497271, located 187 bp from rs12913832. In melanocytes, AG2 predicts that both rs12913832 and rs6497271 reduce expression of the pigmentation gene *OCA2*, with little or no effect predicted for the other variants in linkage disequilibrium (Supplementary Fig. 15a). This agrees with massively parallel reporter assay, chromatin interaction and CRISPR interference experiments that identified rs6497271 as a functional variant in the same *OCA2* enhancer^22^. Carrying both derived alleles is common in individuals of European ancestry. AG2 predicts that the effect of this haplotype is approximately the sum of the two individual effects, in line with its lowest enhancer activity in reporter assays^22^.

By contrast, at the *SMIM1* locus, AG2 predicts opposite effects in erythroblasts for two variants only 38 bp apart (Supplementary Fig. 15b). Each variant is fine-mapped (PIP > 0.99) in a separate credible set for reticulocyte count^16,17^. The associations match these predictions: rs1175550 is predicted to increase *SMIM1* expression and is associated with increased reticulocyte count, whereas rs6673829 is predicted to decrease expression and is associated with decreased reticulocyte count. We anticipate that AG2 will help to identify further loci with multiple causal variants and to derive allelic series.

AG2 can also resolve the cell type in which a variant acts. The pulmonary fibrosis risk variant rs35705950 is known from bulk lung tissue to increase *MUC5B* expression^23^. *MUC5B* is expressed mainly by airway secretory cells, whereas pulmonary fibrosis primarily affects the distal lung and alveolar epithelium. We applied AG2 to a single-cell atlas of the human distal airways^18^. AG2 predicts that the largest effect of the variant is activation of *MUC5B* in alveolar type 2 (AT2) cells, which express the gene at very low levels at baseline (Supplementary Fig. 15c). Experimental evidence supports this prediction. *MUC5B* is co-expressed with the AT2 marker *SFTPC* in the lungs of individuals with idiopathic pulmonary fibrosis. Moreover, *Muc5b* overexpression in the distal airways and alveoli of mice exacerbates bleomycin-induced lung fibrosis^24^.

Similarly, rs3820282 is associated with endometriosis, uterine leiomyoma and gestational duration^25,26^. In an endometrial cell atlas^19^, AG2 predicts that this variant increases *WNT4* expression, most strongly in endometrial stromal cells. This agrees with mice in which the alternative allele was introduced by CRISPR–Cas9, leading to upregulation of transcription of *Wnt4* in endometrial stroma^25^ (Supplementary Fig. 15d).

### Supplementary Tables

**Supplementary Table 1.** References for single-cell pre-training studies in the AlphaGenome 2 corpus. Summary of the 36 single-cell and single-nucleus RNA-seq, ATAC-seq, and paired multiome studies used during AG2 pre-training; repository accessions, processing details, and curation notes are provided in Supplementary Table 3.

| Study / Dataset | Paper DOI | Reference |
|---|---|---|
| ENCODE Multiome | 10.1038/s41586-025-09909-9;<br>10.1038/s41467-025-64343-9 | Moore *et al.* (2026)<sup>27</sup>;<br>Kagda *et al.* (2025)<sup>28</sup> |
| ENCODE Single-Cell (Unpaired) | 10.1038/s41586-025-09909-9;<br>10.1038/s41467-025-64343-9 | Moore *et al.* (2026)<sup>27</sup>;<br>Kagda *et al.* (2025)<sup>28</sup> |
| Bone Marrow Multiome | openreview.net/forum?id=gN35BGa1Rt;<br>10.1038/s41592-025-02854-5 | Luecken *et al.* (2021)<sup>29</sup>;<br>Hingerl *et al.* (2025)<sup>30</sup> |
| K562 Multiome / ISSAAC-seq | 10.1038/s41592-022-01601-4 | Xu *et al.* (2022)<sup>31</sup> |
| WTC11 iPSC Differentiation | 10.1016/j.stem.2018.09.009 | Friedman *et al.* (2018)<sup>32</sup> |
| E2G Primary Astrocytes | 10.1038/s41593-025-02154-3 | Green *et al.* (2026)<sup>8</sup> |
| mESC Multiome | 10.1016/j.celrep.2025.116089 | Nariya *et al.* (2025)<sup>33</sup> |
| Human Developmental Multiomic Atlas / HDMA | 10.1038/s41586-026-10326-9 | Liu *et al.* (2026)<sup>34</sup> |
| Mouse Gastrulation Multiome | 10.1101/2022.06.15.496239 | Argelaguet *et al.* (2022)<sup>35</sup> |
| Human Gut Cell Atlas | 10.1038/s41586-021-03852-1 | Elmentaite *et al.* (2021)<sup>36</sup> |
| Human CNS Glia Atlas | 10.1186/s40478-023-01568-z | Seeker *et al.* (2023)<sup>37</sup> |
| Cross-Tissue Immune Atlas | 10.1126/science.abl5197 | Domínguez Conde *et al.* (2022)<sup>38</sup> |
| Developing Human Immune Atlas | 10.1126/science.abo0510 | Suo *et al.* (2022)<sup>39</sup> |
| Developing Human Cerebral Cortex Multiome | 10.1126/sciadv.adg3754 | Zhu *et al.* (2023)<sup>40</sup> |
| Human Breast Multiome | 10.1038/s41591-024-03011-9 | Bhat-Nakshatri *et al.* (2024)<sup>41</sup> |
| Human Early Pregnancy Multiome | 10.1038/s41586-023-05869-0 | Arutyunyan *et al.* (2023)<sup>42</sup> |
| Human Heart Multiome | 10.1038/s41586-023-06311-1 | Kanemaru *et al.* (2023)<sup>43</sup> |
| Human Liver Atlas | 10.1016/j.cell.2021.12.018 | Guilliams *et al.* (2022)<sup>44</sup> |
| Human Lung Cell Atlas | 10.1038/s41588-022-01243-4 | Madissoon *et al.* (2023)<sup>45</sup> |
| SEA-AD Human Brain Cortex | 10.1038/s41593-024-01774-5 | Gabitto *et al.* (2024)<sup>46</sup> |
| Human Subcutaneous & Visceral Adipose Atlas | 10.1038/s41588-024-02048-3 | Lazarescu *et al.* (2025)<sup>47</sup> |
| Tabula Sapiens v2 Bladder | 10.1101/2024.12.03.626516 | The Tabula Sapiens Consortium & Quake (2024)<sup>48</sup> |
| Single-Nucleus Cross-Tissue GTEx Atlas | 10.1126/science.abl4290 | Eraslan *et al.* (2022)<sup>49</sup> |
| KPMP Human Kidney Atlas | 10.1101/2025.09.26.678707 | Lake *et al.* (2025)<sup>50</sup> |
| Asian Immune Diversity Atlas PBMC | 10.1016/j.cell.2025.02.017 | Kock *et al.* (2025)<sup>51</sup> |
| PBMC snATAC-seq | 10.1371/journal.pgen.1010759 | Benaglio *et al.* (2023)<sup>52</sup> |
| Human Retina Multi-Omics Atlas | 10.1101/2023.11.07.566105 | Li *et al.* (2023)<sup>53</sup> |
| Mouse Retina Atlas | 10.1016/j.isci.2024.109916 | Li *et al.* (2024)<sup>54</sup> |
| Human First-Trimester Neurodevelopment | 10.1038/s41586-024-07234-1 | Mannens *et al.* (2025)<sup>55</sup> |
| Human Thymus Cell Atlas | 10.1038/s41586-024-07944-6 | Yayon *et al.* (2024)<sup>56</sup> |
| Mammalian Neocortex Multiome | 10.1038/s41586-023-06819-6 | Zemke *et al.* (2023)<sup>57</sup> |
| Human Skeletal Muscle Aging Atlas | 10.1038/s43587-024-00613-3 | Kedlian *et al.* (2024)<sup>58</sup> |
| PanSci Mouse Aging scRNA Atlas | 10.1126/science.adn3949 | Zhang *et al.* (2025)<sup>59</sup> |
| Mouse Aging EasySci-ATAC Atlas | 10.1126/science.adw6273 | Lu *et al.* (2026)<sup>60</sup> |
| Mouse Brain Aging EasySci Atlas | 10.1038/s41588-023-01572-y | Sziraki *et al.* (2023)<sup>61</sup> |
| ChromBPNet Reference scATAC (Microglia & Coronary Artery SMC) | 10.1101/2024.12.25.630221;<br>10.1038/s41588-020-00721-x;<br>10.1038/s41588-022-01069-0 | Pampari *et al.* (2024)<sup>62</sup>;<br>Corces *et al.* (2020)<sup>63</sup>;<br>Turner *et al.* (2022)<sup>64</sup> |

